# Polymerase Evolution Enables Access to Glycosylated Xeno-Nucleic Acids with Expanded Chemical Functionality

**DOI:** 10.64898/2026.07.20.739590

**Authors:** Eloi Vincent, Victoria Maola, Clément Lopez, Marc Borie-Guichot, Maria Dalla Pozza, Mohammad Hajjar, Rémi Sieskind, Soizick Lucas-Staat, Rafael Navaza, Laurence A. Mulard, Marcel Hollenstein, John C. Chaput, Marc Delarue

**Affiliations:** Institut Pasteur, Université Paris Cité, CNRS UMR 3528, Unit of Architecture and Dynamics of Biological Macromolecules, 25 rue du Docteur Roux, 75015 Paris, France; Institut Pasteur, Université Paris Cité, CNRS UMR 3523, Unit of Chemistry of Biomolecules, 28 rue du Docteur Roux, 75724 Paris Cedex 15, France; Institut Pasteur, Université Paris Cité, CNRS UMR 3523, Unit of Bioorganic Chemistry of Nucleic Acids, 28 rue du Docteur Roux, 75724 Paris Cedex 15, France; Institut Pasteur, Université Paris Cité, CNRS UMR 3528, Plate-forme de Cristallographie-C2RT, 28 rue du Docteur Roux, 75724 Paris Cedex 15, France; Department of Pharmaceutical Sciences, University of California, Irvine, CA, USA; Department of Chemistry, University of California, Irvine, CA, USA; Department of Molecular Biology and Biochemistry, University of California, Irvine, CA, USA; Department of Chemical and Biomolecular Engineering, University of California, Irvine, CA, USA

## Abstract

Chemical modifications expand the functional repertoire of nucleic acids, but the extent to which complex glycans can be integrated into genetic polymers remains largely unexplored. Glycosylated nucleic acids, including glycoRNAs and bacteriophage genomes, are emerging as key mediators of host-pathogen interactions, yet their synthetic accessibility is severely limited by the inability of polymerases to process substrates bearing simultaneous base and backbone modifications. Here we report the directed evolution of a family B DNA polymerase that enables the synthesis of glycosylated xeno-nucleic acids (XNAs). Using fluorescence-activated droplet sorting, we identified an engineered polymerase, termed G2, that efficiently incorporates nucleotides bearing mono- to trisaccharide base-appendages together with C2’ substitutions, yielding XNAs containing up to 30% glycosylated bases. Deep sequencing of >93,000 chimeras reveals that this activity arises from epistatic networks concentrated within catalytic domains. These results establish a route to glycan-encoded nucleic acids and advance polymerase engineering for chemically expanded therapeutics.

## Introduction

Nucleic acids are frequently chemically modified to regulate gene expression or evade cellular defense systems, with methylation, mono-glycosylation or oligosaccharides representing promising examples^1–4^. Bacteriophages provide a striking illustration of this principle, as T2 or T4 phages, for instance, incorporate extensively glycosylated nucleobases into their genomic DNA, enabling resistance to host endonucleases and CRISPR-mediated immunity^1,5,6^ **(Figure 1a)**. While glycosylation is well established in the context of DNA, lipids, and membrane proteins^7^, its role in eukaryotic RNA has only recently come into focus. Striking among these are glycosylated RNAs (glycoRNAs), an emerging new class of post-transcriptional modifications in which small non-coding RNAs are conjugated to N-linked glycans displayed on the cell surface^8,9^. These structures have been implicated in cellular communication, immune regulation, and disease progression^10^. From a therapeutic perspective, glycosylation offers a compelling strategy to expand the functional landscape of nucleic acids, with the potential to enhance target specificity, modulate immunogenicity, and improve delivery^11^. Indeed, analogous approaches are already used in protein-based vaccines and in oligonucleotide therapies as impressively demonstrated by the conjugation of trivalent *N*-acetyl-D-galactosamine ligand (D-GalNAc) to siRNAs for hepatocyte targeting^12,13^.

**Figure 1.**
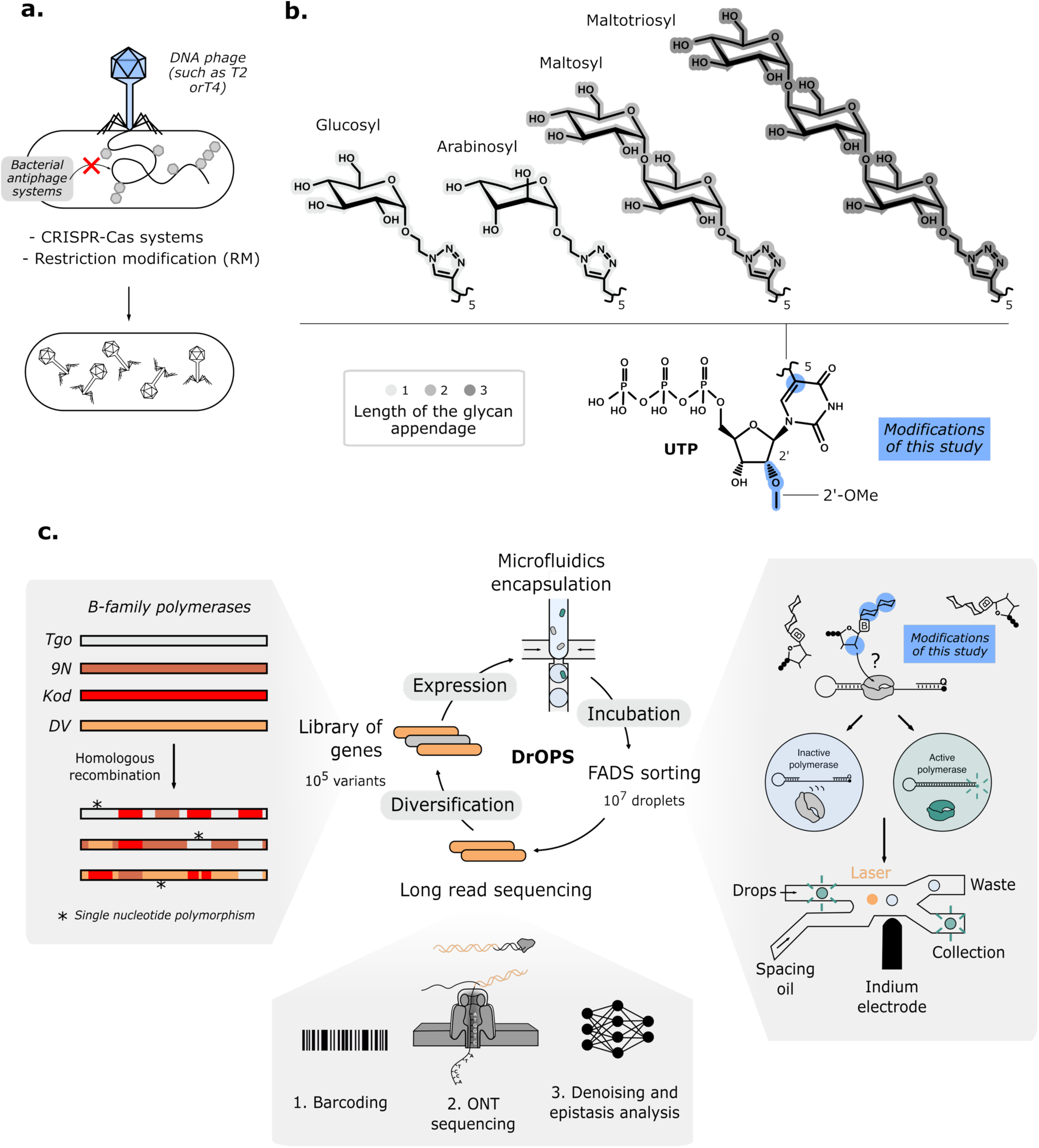
Directed evolution strategy for engineering family-B polymerases synthesizing glycosylated 2’-OMe-XNA. **(a)** Some phages such as T2 or T4 glycosylate their DNA to evade bacterial defense systems such as CRISPR-Cas and restriction-modification (RM) mechanisms. Mimicking these glycosylations for RNA strands could expand the chemical spectrum for RNA therapeutics. **(b)** Chemical structures of modified nucleotides that bear mono- and short oligosaccharides(up to maltotriose) grafted via click chemistry onto the uridine base, along with 2’ modifications (2’-OMe). In blue are the positions modified in this study. **(c)** Directed evolution experiments applied in this study use library diversification and screening methods described in^27,60^. A gene library was generated via homologous recombination from four B-family polymerase parents carrying RNA polymerase-enhancing mutations (V93Q, Y410G, A484L, E664K) and exo(-) inactivating mutations (D141A, E143A). Expressed in bacteria and encapsulated via microfluidics with a DNA substrate containing a fluorophore/quencher along with hyper-modified nucleotides, active polymerases displace the quencher to induce droplet fluorescence. This enables recovering of active mutants by fluorescence-activated droplet sorting (FADS). Enriched libraries undergo further sorting cycles or are analyzed with long-read Oxford Nanopore sequencing.

Enzymatic tools for the synthesis of hypermodified RNAs remain limited, as canonical RNA polymerases are often poorly suited to accommodate bulky chemical modifications^14,15^. In contrast, DNA-dependent DNA polymerases (DNAP) provide a more tractable engineering platform owing to their high fidelity, monomeric architecture, and intrinsic thermostability. These enzymes, classified into families A to D, RT, X and Y based on sequence homology^16^, share common catalytic motifs (A-C) that coordinate DNA synthesis via a primer extension mechanism^17,18^. Substrate selectivity is governed in part by a conserved “steric gate” residue that discriminates against ribonucleotides (NTPs) by sterically excluding the 2ʹ hydroxyl group found in NTPs^19,20^. Mutation of this residue to a smaller side chain (for example, Y409G), relaxes C2’ selectivity and enables the incorporation of multiple ribonucleotides. However, efficient extension of long RNA products requires additional mutations at secondary checkpoint, such as E664K, often referred to as a second steric gate, highlighting the complex layers of constraints that limit polymerase specificity^21,22^.

Thermophilic and processive family B DNA polymerases have emerged as leading platforms for the synthesis of RNA and XNA through extensive protein engineering^23,24^. Foundational efforts identified a set of mutations in *Thermococcus gorgonarius* (Tgo) polymerase, commonly referred to as Tgo-GQLK, that collectively relax substrate specificity and enhance RNA synthesis. These include V93Q, which alleviates uracil-stalling^25^, the steric gate mutation Y409G, the Therminator mutation A485L^26^ and E664K, an auxiliary site mutation that improves RNA extension efficiency^22^. Building on this framework, subsequent variants have achieved substantial gains in activity and substrate scope. Notably, the chimeric C28 polymerase developed in Chaput’s team exhibits a higher RNA synthesis rate of up to 3 nts^-1^ ^27^. Other design efforts identified I521L and F545L as mutations that broaden substrate tolerance^24,28^ or T541G and K592A (the so-called “2M” mutations), which enable the synthesis of longer 2ʹ-modified RNA^29^. Together, these studies reveal both the progress and remaining limitations in engineering polymerases with expanded substrates specificities.

Despite extensive functional and structural characterization of family B polymerases^30–32^, rational design efforts remain inefficient, with only a limited number of successful engineering campaigns reported^33,34^, and none capable of supporting the incorporation of highly modified nucleotides. In this context, directed evolution has emerged as an effective strategy for generating polymerases with expanded substrate scope. A range of selection platforms have been developed, including phage display^35–37^, self-selection systems such as compartmentalized self-replication (CSR)^38,39^, and fluorescence-based methods including compartmentalized bead labeling (CBL)^40^ and fluorescence-activated droplet sorting (FADS)^41^. While these approaches have enabled the isolation of polymerases capable of synthesizing diverse XNAs, extending this capability to substrates bearing multiple, structurally complex modifications remains a major unmet challenge in the field.

Here, we report a directed evolution campaign that relies on microfluidic-based droplet sorting to engineer polymerases that are capable of incorporating nucleotides bearing base- and C2’-sugar modifications on the same substrate **(Figure 1b)**. Starting from a recombined library of four family B polymerases harboring the QGLK mutations (Tgo, *Thermococcus sp. 9°N* (9°N), *Pyrococcus sp. GB-D Deep Vent* (DV), *Thermococcus Kodakarensis* (Kod)) **(Figure 1c)**, we identified a highly efficient variant, termed G2. This enzyme enables the synthesis of hypermodified RNAs containing both 2ʹ-O-methyl substitutions and nucleobased decorated with variable-length and composition glycans, including up to trisaccharides. Deep sequencing of the evolving populations reveals the recombination-driven trajectories underlying the emergence of this activity, providing mechanistic insight into how distributed mutations cooperate to expand substrate scope. These findings establish a general strategy for engineering polymerases with access to chemically complex substrates and highlight the potential for recombination-based evolution to explore the functional landscape of protein sequence space.

## Results

### Fluorescence activated droplet sorting

Screening of eight previously characterized polymerases from families A^35,42,43^ and B^29,44^, including the recently evolved Tgo-2M, revealed detectable but very low incorporation activity toward the hypermodified nucleotides bearing both C5-base glycosylation and C2’-ribose modifications employed in this study **(Supplementary Figure 2, Supplementary Figure 3)**. These nucleotides were synthesized by application of a recent approach developed for their corresponding dNTPs **(Supplementary Figure 1)**^45^. Briefly, the requisite nucleotide building blocks were prepared from a common 5-ethynyl-2’-*O*-methyl-UTP intermediate in 10 steps from ribose **(Supplementary Figure 1a)**. This synthon was then subjected to a copper-catalyzed azide-alkyne cycloaddition (CuAAC) reaction using a variety of mono-, di-, and tri-saccharides equipped with azide moieties. The compounds obtained are the 5-glucosyl-triazolylmethyl-2’-*O*-methyl-UTP, 5-arabinosyl-triazolylmethyl-2’-*O*-methyl-UTP, 5-maltosyl-triazolylmethyl-2’-*O*-methyl-UTP and 5-maltotriosyl-triazolylmethyl-2’-*O*-methyl-UTP. For clarity, these compounds are referred to here as 5-glycosyl-2’-*O*-methyl-UTP derivatives (sugar precised if necessary). **(Supplementary Figure 1b)**.

Iterative directed evolution rounds of high-throughput screening were performed starting from a library derived from the C28 RNA polymerase campaign^27^. This library was generated by DNA shuffling of four hyperthermophilic B-family DNA polymerases (Tgo, 9°N, Kod and DV) which share pairwise sequence identities from 82 to 93%. Each of the four 2.4 kb genes were divided in 11 segments of variable lengths and the resulting segments were shuffled by homologous recombination as described in^46^. The shuffled library was further designed to partly replace (around 30% of the library) a specific segment with one derived from Tgo and encoding the TGLLK and 2M mutations (I521L, T541G, F545L and K592A). These mutations have been described as another secondary steric gate of B-family polymerases and help suppress the discrimination between deoxyribo- and ribo-nucleotides^29^.

Library variants were compartmentalized in droplets with a polymerase activity assay mixture, containing primers enabling optical detection (fluorescent template and quencher), the 5-maltosyl-2ʹ-*O*-methyl-UTP **(Figure 1b)** as the modified substrate, and the remaining natural NTP substrates. The substrate was selected based on its extended glycosylation chain and superior synthesis yield. Thermal lysis released polymerases from expressing cells and provided a selective pressure for thermostability. Active variants extended a fluorescent self-priming hairpin template, disrupting fluorescence quenching and generating a signal proportional to XNA synthesis. Encapsulation is followed by screening rounds with FADS chips^47,48^, enabling massively parallel screening at 1 kHz (1,000 drops per second) **(Figure 1c)**. Four rounds were conducted, sorting 4.7 to 10.9 × 10^6^ droplets each, with the final round using a shortened 2 h extension time (versus 6 h initially) **(Supplementary Table 1)**.

Over the course of evolution, the fraction of droplets with high fluorescence and considered to be active polymerases increased from 5.4% (initial library, λ = 0.1 occupancy) to 54% (final library), representing a 10-fold average improvement **(Figure 2)**. The initial and final libraries were then sequenced to follow the evolutionary paths leading to this unnatural activity. Oxford Nanopore long-read sequencing yielded 1.8 × 10^5^ error-corrected reads for the initial library and 1.4 × 10^5^ for the final library which is close to the expected size of the library around 10^5^ variants. Long-read sequencing was necessary due to the length of the genes (≈2,400 nts) and the need to unambiguously know the identity of the residues in each sequence. The proportion of unique variants dropped from 43% to 9.6% along the evolution, indicating enrichment of a high-performing subset from the initial diversity. Screening 96 clones randomly chosen from the final library identified a lead variant, called G2, with a significantly enhanced activity for incorporating the full set of 5-glycosylated-2ʹ-*O*-methyl uridine into RNA matrices. After sequencing, G2 was detected in around 18% of the library, whereas the previous C28 was completely absent.

**Figure 2.**
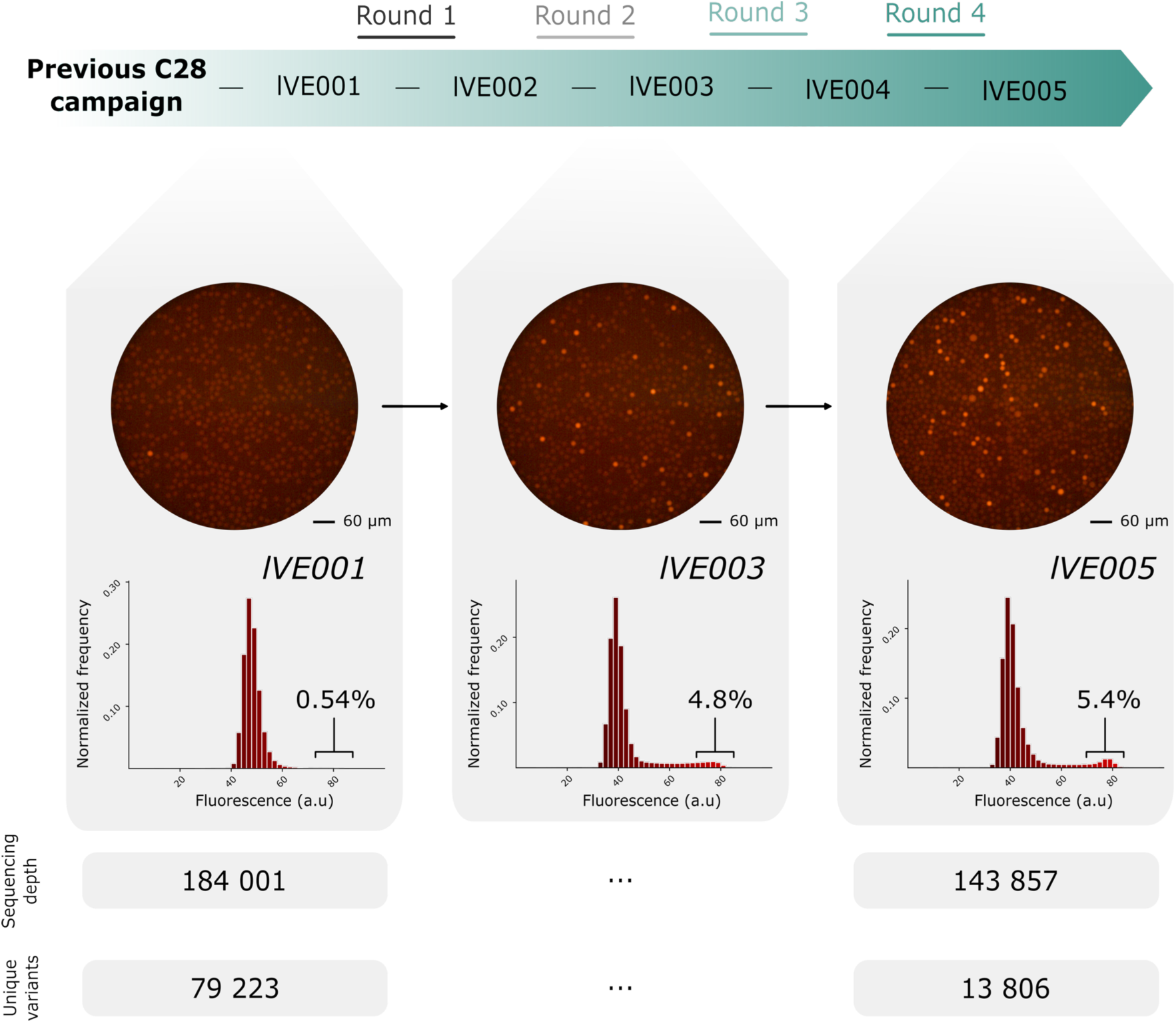
Experimental evolution of B-family polymerase population. Four cycles of directed evolution were performed starting from a shuffled B-family polymerase library derived from the campaign that yielded the C28 RNA polymerase^27^. Library activity was assessed throughout evolution by measuring the fraction of droplets exceeding 70 a.u. fluorescence. This fraction increased from 0.54% (equivalent to 5.4% at λ=0.1 occupancy) to 5.4% (54%), reflecting a 10-fold average enrichment in activity. Initial and final libraries were sequenced, yielding more than 184,000 and 143,000 reads, respectively. The proportion of unique variants dropped from 43% to 9.6%, indicating enrichment of a small subset of active polymerases from the starting library.

### Variant G2 outperforms C28 RNA polymerase for XNA synthesis

Activity of variant G2, isolated after four rounds of directed evolution, was quantified using a primer-extension assay with various glycosylated ribonucleotides. In the first assay **(Figure 3a)**, time courses experiments performed on DNA templates containing four adenosine nucleotides were analyzed by capillary electrophoresis. Conversion was measured as the fraction of fully extended product (+10 nts) over stalled intermediate species. Apparent constant *k_obs_* rates were then estimated. Canonical dNTPs, NTPs, and NTPs in which UTP was substituted by 5-glucosyl-2’-*O*-methyl-UTP or by 5-maltosyl-2’-*O*-methyl-UTP (the substrate used for selection) were tested. G2 showed slightly slower kinetics for both DNA and RNA synthesis compared with the highly active C28 RNA polymerase, generated in previous campaigns aimed at improving its RNA synthesis ability^27^. For the incorporation of 5-glucosyl-2’-*O*-methyl-UTP, the difference was more pronounced, wherein the evolved G2 variant exhibited an 11-fold higher apparent rate constant *k_obs_* and 87% conversion after 30 min versus 55% for C28 (0.95s^-1^ versus 0.083s^-1^). For the 5-maltosyl-2ʹ-*O*-methyl substrate, G2 displayed an almost sixfold increase in *k_obs_* and achieved more than 85% conversion after 30 min, whereas C28 stalled at 16% conversion (0.10s^-1^ versus 0.018s^-1^). The fidelity of G2 was also estimated to be ≈2-fold higher. The selected variant indeed exhibited a 2-fold reduced misincorporation rate that would correspond to a >99% fidelity. **(Supplementary Figure 4).**

**Figure 3.**
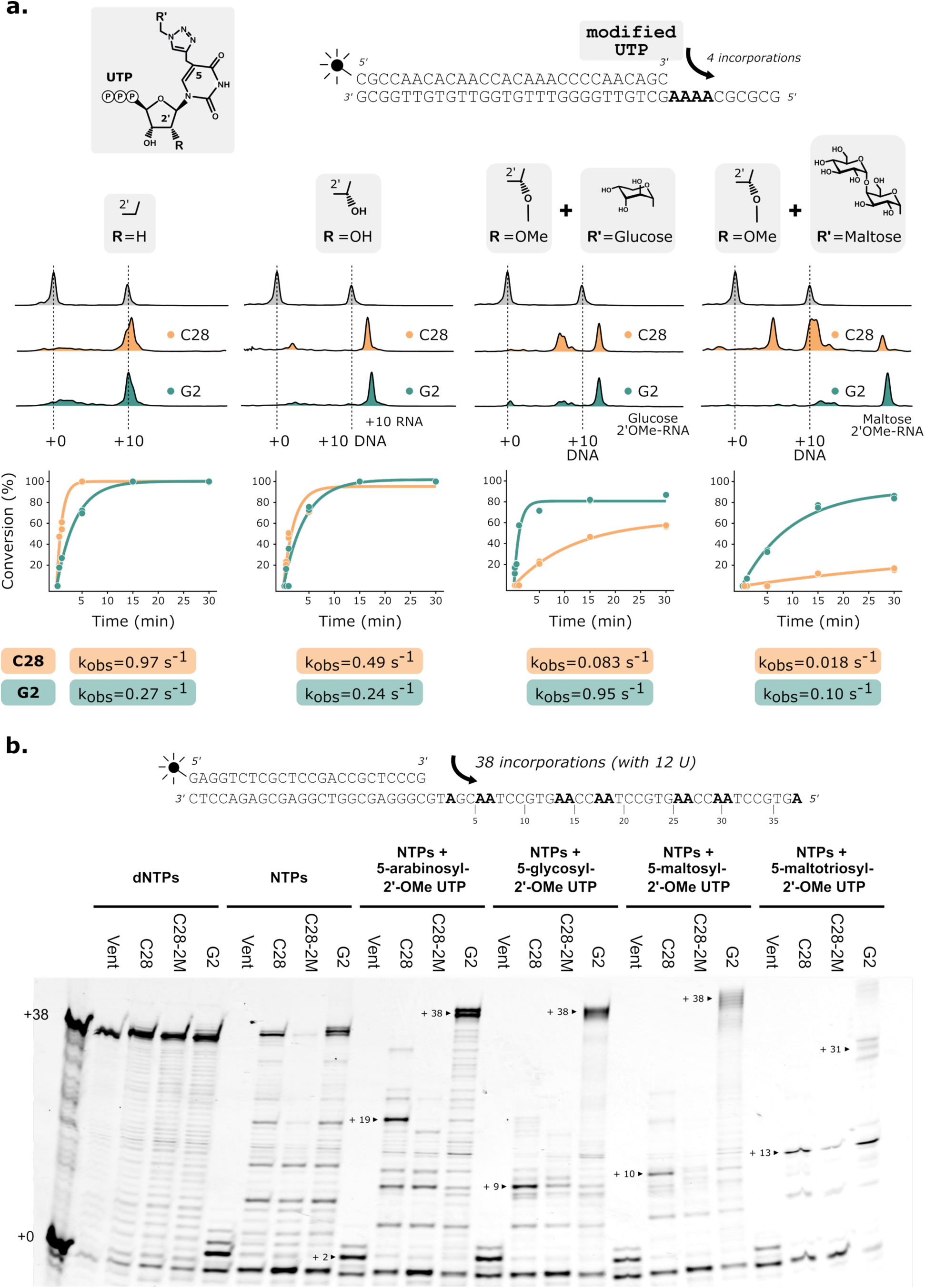
Functional characterisation for G2 polymerase activity for hyper-modified nucleotide incroporation. **(a)** Capillary electrophoresis of primer extension assays shows incorporation of four nucleotides by C28 (yellow) and G2 (green) polymerases using three sets of substrates: dNTPs, NTPs, and NTPs with UTP replaced by 5-glucosyl-2’-*O*-methyl-UTP or 5-maltosyl-2’-*O*-methyl-UTP. **Top:** primer-template duplex used. **Middle:** Capillary profiles after 30 min extension reveal unextended primer (+0), DNA product, RNA, and slower-migrating glycosylated RNA (glucosyl or maltosyl). Bottom: Full-length conversion (duplicates) fits single-exponential model (solid lines) for extracting apparent rate constants (*k_obs_*). G2 is slower for DNA or RNA synthesis but 6-fold faster for 5-maltosyl-2’-OMe-UTP and 11-fold faster for 5-glucosyl-2’-*O*-methyl-UTP nucleotide incorporation. **(b)** Primer extension assays for the incorporation of four increasingly glycosylated 2’-OMe UTP (5-arabinosyl, 5-glucosyl, 5-maltosyl and 5-maltotriosyl) with four different family-B DNA polymerases Vent(exo⁻), C28, C28-2M (from^29^), and G2. G2 completes incorporation of 12 modified nucleotides with 5-arabinosyl, 5-glucosyl or 5-maltosyl modified 2’-*O*-methyl-UTP (vs. 7, 3 and 3 for C28) and 11 on 5-maltotriosyl-2’-*O*-methyl-UTP (vs. 3 for C28).

G2 was benchmarked across all glycosylated substrates of this study mixed with NTPs against Vent(exo^-^), C28, and C28 bearing the 2M mutations as previously described^29^. The Vent(exo^-^) DNAP was chosen as the commercially available B-family control since this enzyme has been shown to tolerate the corresponding modified dNTPs^45^ and bulky modifications^49^. For three of the four glycosyl-nucleotides tested in **Figure 3b**, G2 reached essentially complete conversion to fully extended RNA product (+38 nt), which includes 12 sites for glycosylated nucleotide incorporation. Even for the most challenging trisaccharide appendage, G2 produced 31-nucleotide RNA, representing 11 of the 12 possible glycosylated nucleotide incorporations. These conversion levels substantially exceeded those of C28, which stalled for the 5-arabinosyl-modified nucleotide at position +19 (7 of 12 U analogues incorporated) and for maltotriose-modified nucleotide at position +13 (3 of 12U). On the other hand, the Vent(exo^-^) DNAP failed to incorporate more than two NTPs under the same conditions. Despite the expectation that a C5-linked sugar residue pointing away from the DNA major groove would minimally affect nucleotide incorporation, the data reveal clear sugar-dependent differences in incorporation efficiency. A similar effect had been observed with similarly modified dNTPs on various templates^45^. Introducing the 2M mutations into C28 did not significantly enhance activity toward these nucleotides modified at both the 2ʹ position and at position 5 of the nucleobase.

To assess the substrate scope of G2, we evaluated its capacity to incorporate a broad panel of modified uridine analogs across an 38-nt template requiring 12 consecutive modified incorporations. G2 achieved complete full-length product synthesis on seven of eight substrates tested, including pseudouridine, 2’-fluoro (2’-F), 2’-amino (2’-NH_2_), locked nucleic acid (LNA), hexitol nucleic acid (HNA), 2’-*O*-methyl (2’-OMe), and 5-ethynyl-2’-*O*-methyl nucleotides **(Figure 4a-g)**. Notably, G2 demonstrated a superior incorporation efficiency relative to the parental enzyme C28 on all substrates except LNA, where C28 looks more tolerant to this modification **(Figure 4d)**. This broad substrate tolerance suggests that directed evolution reshaped the active site geometry of G2 to accommodate diverse 2’ and base modifications without imposing strict steric constraints. Out of the 8 non-natural substrates that were tested, the sole exception was the 2’-*O*-methoxyethyl (2’-MOE) modification pattern, on which G2 failed to reach completion to full-length product **(Figure 4h)**. The bulky aliphatic chain of the 2’-MOE modification likely imposes steric constraints that exceed the conformational flexibility acquired during the evolution campaign. Collectively, these results establish G2 as a highly versatile XNA polymerase with substantially expanded substrate scope relative to C28, capable of synthesizing heavily modified oligonucleotides of potential direct therapeutic relevance.

**Figure 4.**
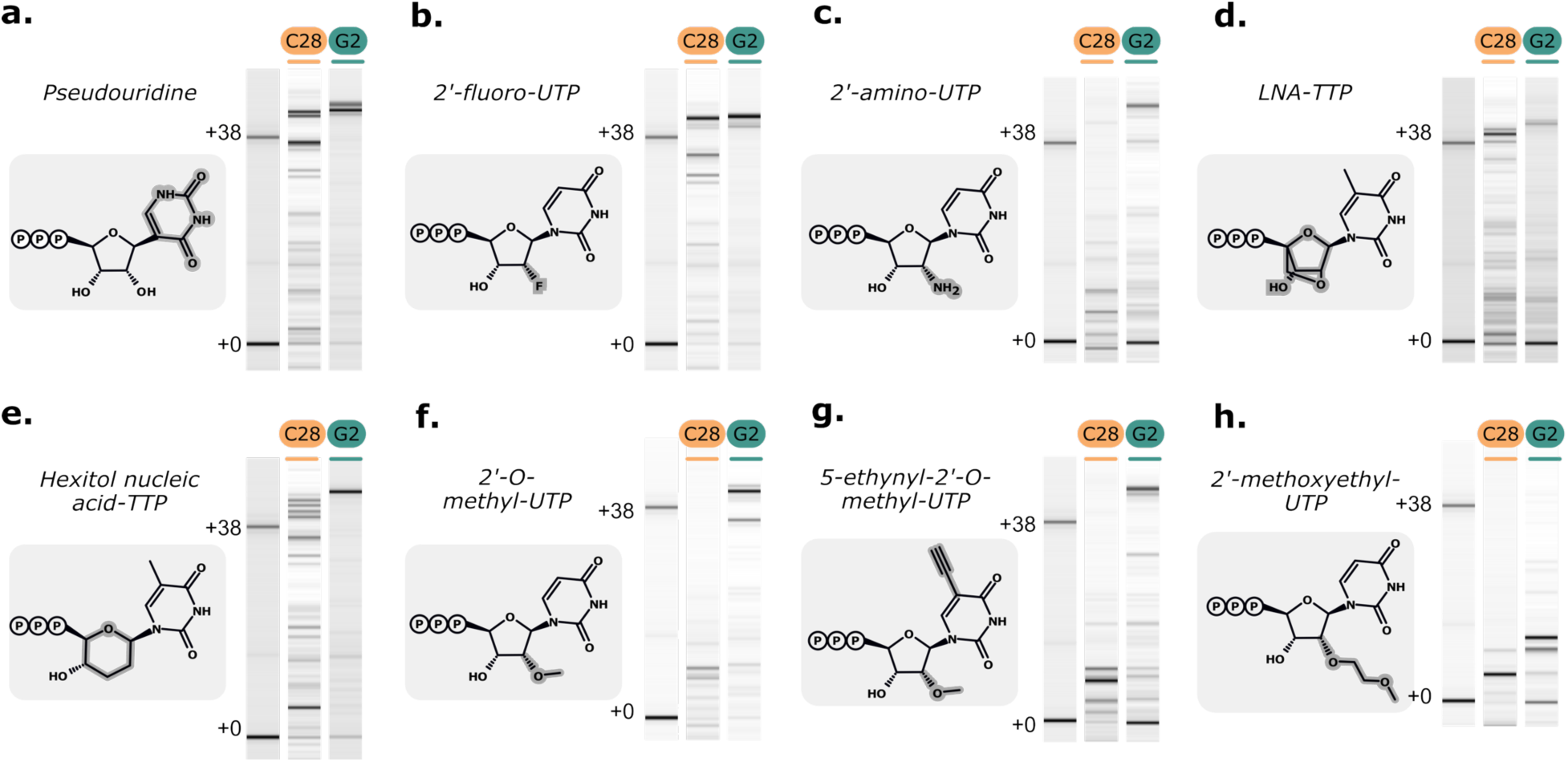
Biochemical characterization of evolved polymerase G2 versus C28 for XNA synthesis. Capillary electrophoresis analysis of primer extension reactions designed to incorporate 12 modified uridine residues using engineered C28 and G2 XNA polymerases. Modified nucleotides include: pseudoUTP **(a)**, 2’-fluoro (2’-F) **(b)**, 2’-amino (2’-NH_2_) **(c)**, LNA **(d)**, HNA **(e)**, 2’-*O*-methyl (2’-OMe) **(f)**, 2’-*O*-methyl with 5-ethynyl base-modification **(g)**, and 2’-*O*-methoxyethyl (2’-MOE) **(h)**. The marker lane consists of the expected DNA product (+38) and the initial primer (+0). G2 achieved full-length product synthesis on substrates **(a)** to **(g)** but failed to reach completion on the 2’-MOE substrate **(h)**. G2 outperformed C28 on all substrates except LNA **(d)**.

Variant G2 comprises nine parental segments and carries only the predefined mutations shared across parents: the RNA polymerase-promoting QGLK substitutions (V93Q, Y410G, A486L, E665K) and 3ʹ-5ʹ exonuclease-inactivating mutations (D141A, E143A). 62% of the G2 sequence was assigned to parent Tgo-QGLK, 7.5% to DV-QGLK, 25% to Kod-QGLK and 5.5% to 9°N-QGLK. The C28 was more balanced between parents (45.4% Tgo-QGLK, 14.8% DV-QGLK, 16.1% Kod-QGLK and 23.8% 9°N-QGLK). The shuffled segments were mapped onto the Kod polymerase structure (PDB:5OMF). Notably, the 9°N segment specifically replaces the fingers domain between residues 450 and 501. Domain transitions in G2 coincide with segment boundaries. For instance, the end of the N-terminal domain is a transition between DV and Tgo. Similarly the end of the 3ʹ-5ʹ exonuclease domain is a shift from Kod to Tgo. The transition between palm and fingers is also marked by a switch of parents **(Figure 5a,b)**. Critically, the catalytic motifs in the polymerase domain remain from a single parent and are never splitted between species. Analysis revealed that motif A originated from a Kod segment, motif B from 9°N, and motif C from a Tgo domain.

**Figure 5.**
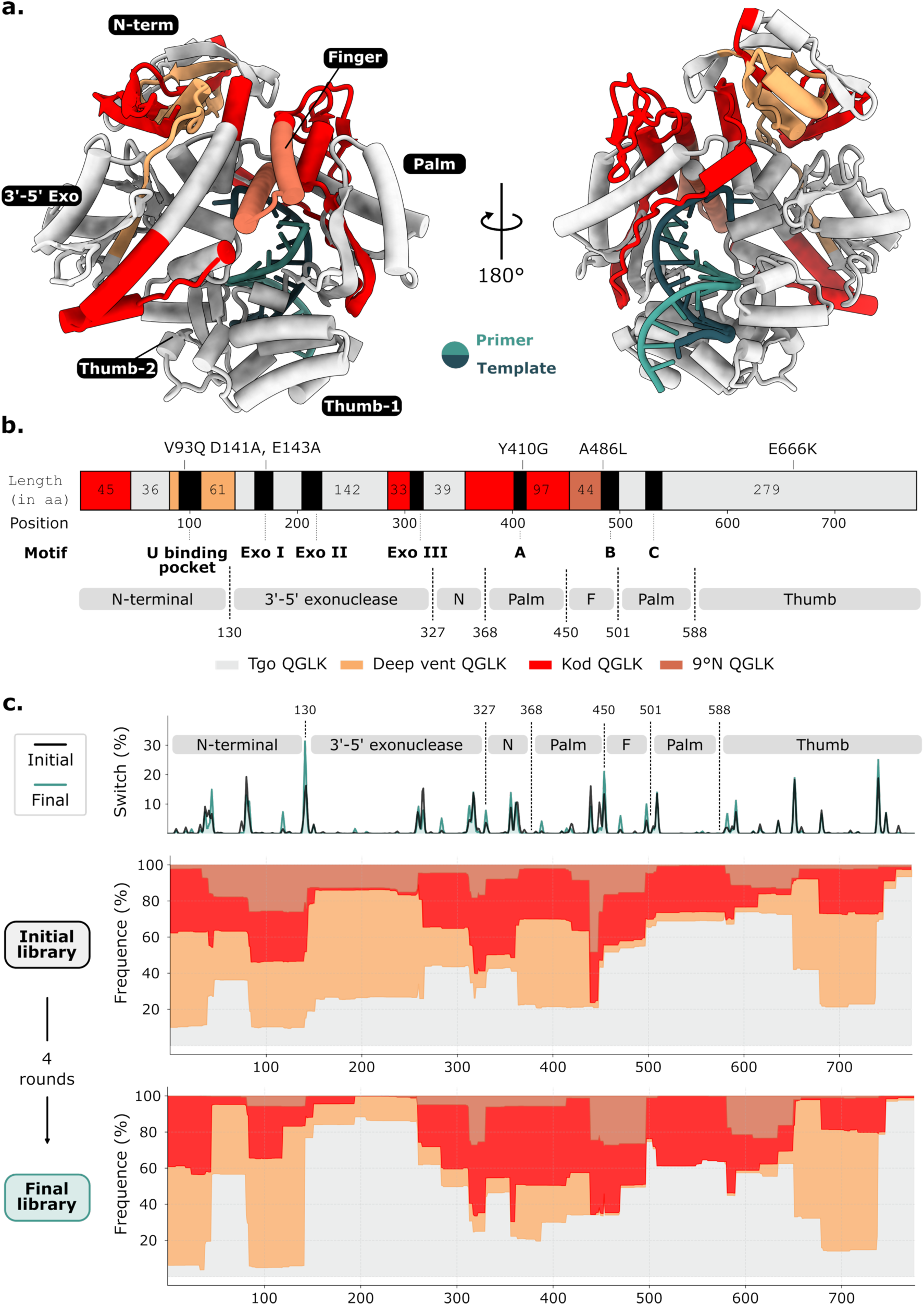
Recombinated fragments of the evolved polymerase G2 and selection dynamics. **(a)** The identified variant G2 comprises nine sequence blocks derived from the four parental polymerases used to generate the homologous recombination library. Blocks are mapped onto the structure of Kod DNA polymerase (PDB: 5OMF). Fingers domain originates from 9°N polymerase and recombination junctions generally avoid secondary-structure elements, with the exception of the longest α-helix in the 3ʹ-5ʹ exonuclease domain. **(b)** Annotated G2 sequence showing key catalytic motifs (uridine-binding pocket, Exo I–III, motifs A-C) and major structural domains. G2 is composed of 62% Tgo-QGLK, 7.5% DV-QGLK, 25% Kod-QGLK and 5.5% with 9°N-QGLK with no additional mutations. **(c)** Segment analysis in the sequenced initial and final libraries. **(top)** Frequency of recombination events at each amino acid position. **(middle)** Parental contributions along the alignment before and **(bottom)** after four rounds of screening.

### Parental segments dynamics

High-throughput sequencing of initial and final libraries generated sequences for tens of thousands of variants, enabling comparison to pinpoint those enriched because of enhanced glyco-RNA polymerase activity. Using a sliding-window analysis for every sequence, parental segment proportions can be mapped along the sequence **(Figure 5c)**. The initial library was inherently biased (not evenly partitioned at 25% per parent per segment) with some regions showing complete dominance of one parent such as Tgo in the second half of the palm domain. These biases can be explained by the fact that the library was obtained from a previous directed evolution campaign reducing the evolutionary gap to gain in XNA polymerase activity^27^.

Certain segments showed little evolution between initial and final libraries. For instance, the second palm domain stayed dominated by one third Kod and two thirds Tgo. Similarly, the thumb domain core remained predominantly DV (70%) throughout evolution. This evolutionary conservation could be attributed to the essential role these segments play in RNA polymerase activity, while having no discernible impact on XNA polymerase activity. Some regions were much more dynamic and so maybe more important for XNA polymerase activity. Notably, DV segments were enriched in some domains such as in the mid-N-terminal region where DV components largely supplanted parts from Kod and 9°N. In the 3ʹ-5ʹ exonuclease domain, the DV sequence is replaced massively as also is the 9°N sequence by the corresponding Tgo exonuclease modules. The first palm domain and second half of the N-terminal domain were enriched for Kod over DV. Significant enrichment occurred for Kod and 9°N segments at the begining of the thumb domain even if their fraction in the initial library was low.

### Non-parental mutational landscape

In addition to DNA shuffling, homologous recombination can also produce single point mutations. We quantified the enrichment of point mutations within our set of protein sequences. For each position, we calculated the frequency of each amino acid in the initial and final libraries, yielding an enrichment matrix per amino acid and per position. This matrix reveals that most amino acids per position are globally depleted, as shown by the maximal absolute variations being negative (blue squares in **Figure 6a**).

**Figure 6.**
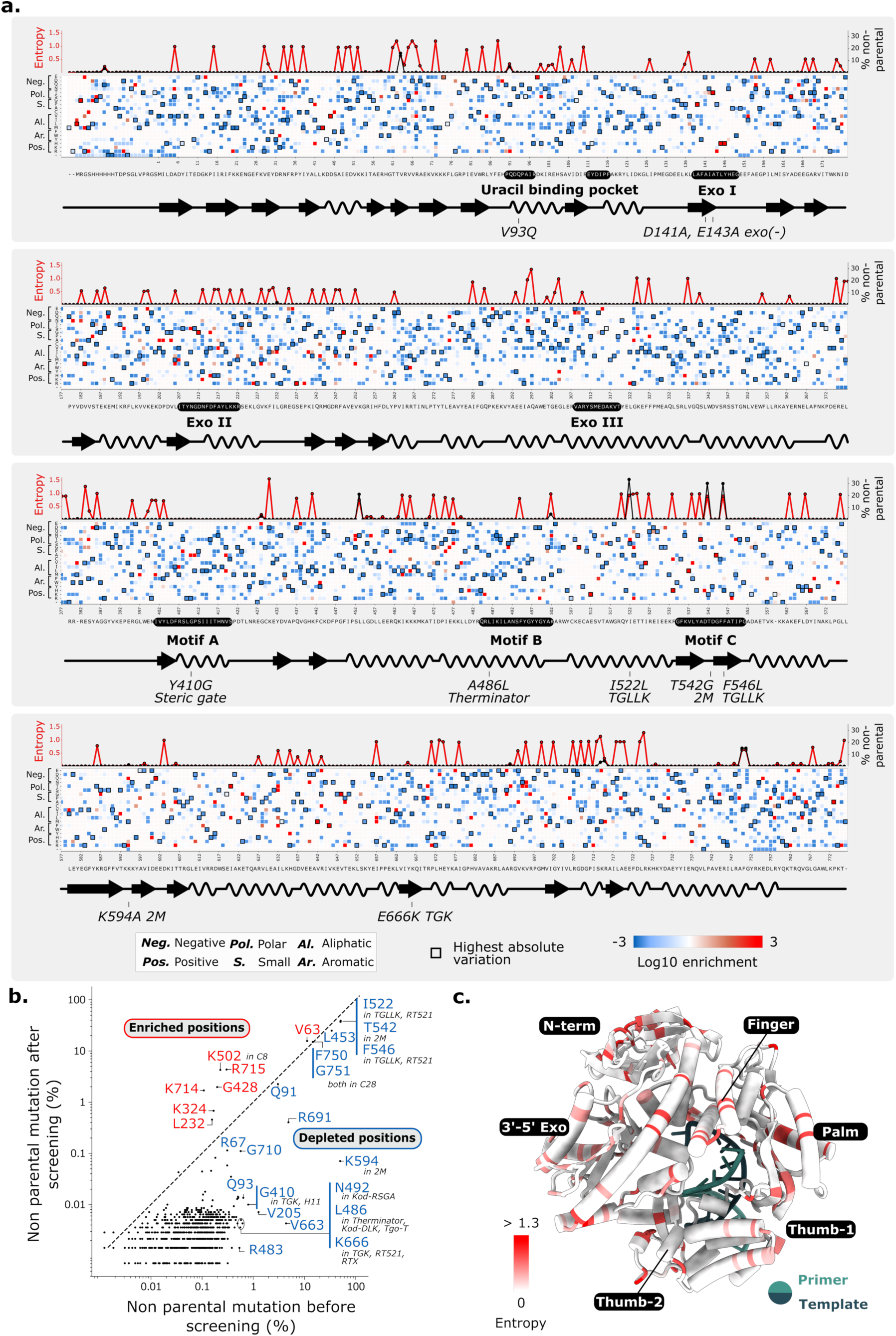
Deep mutational profiling of XNA polymerase directed evolution. **(a)** Enrichment for every amino-acid substitution over the directed evolution rounds. Most substitutions are strongly depleted and some regions appear largely neutral (Exo I, Exo III motifs). Shannon entropy approximating the mutational tolerance is represented **in red** on the top plot and fraction of variants carrying non-parental substitutions (i.e., not derived from the four parents) is **in black**. Catalytic motifs are largely intolerant to mutations (low to near-zero entropy). The printed sequence is the consensus of the alignment of the final library. Substitutions previously reported to modulate incorporation of ribonucleotides or 2ʹ-modified nucleotides are in *italic*. **(b)** Frequency of non-parental mutations are tracked before and after directed evolution. Non-parental mutations are mostly depleted even at sites described in the literature, such as the second residue of the steric gate reported by Freund et al.29 Few positions show enrichment (K502 and R715). TGLLK and the T542 substitution from 2M mutant are maintained at high frequency. **(c)** Shannon entropy is mapped onto the structure of Kod DNA polymerase (PDB: 5OMF). Positions with the highest entropy are distributed throughout the structure.

Certain regions, such as Exo I and Exo III motifs or the region following motif B, exhibit largely neutral enrichment. Consequently, we focused our analysis on positions known to enhance RNA polymerase activity and present in all four parents (QGLK), the uracil binding mutation V93Q, the steric gate Y410G in motif A, the Therminator A486L, and the Tgo-T E666K. All of them display distinct patterns. For V93Q, no alternative is strongly enriched, but V93S is heavily depleted, suggesting a preference for larger polar residues at this site. At the steric gate Y410G, only alanine (Y410A) is enriched, consistent with a small residue also known to avoid 2’-modified nucleotide rejection^34^ while the large and charged arginine shows a strong depletion. For Therminator mutation (A486L) facilitating incorporation of modified nucleotides, alanine is strongly depleted which indicates that reversion to the wild-type residue is unfavorable. At position E666, polar (Q, N, T) and aliphatic (I) residues are strongly depleted, reinforcing the need for a positively charged amino acid at this position (E666K). For 2M and TGLLK mutations introduced into the library, serine is depleted and threonine is enriched at position I522 even if both are polar whereas the intended L stays stable. T542G remains constant while charged residues (lysine, arginine, glutamate) are strongly depleted favoring small uncharged residues at this position. F546 disfavors another aromatic residue, as well as histidine, which is charged. At K594, positive arginine or negative glutamine are enriched which shows that a charged amino acid such as the wild-type lysine is important at this position. The other polars, aliphatics, and small residues (M, V, A) are depleted at this position.

Shannon entropy and non-parental mutational rates were mapped along the sequence. Shannon entropy scores in the final library indicate mutational tolerance per position. The mutational rate of non-parental mutations reflects intentionally introduced or randomly induced point mutations which are different from the four parents. Entropy localizes to subsets of positions (e.g., high values at position 430, 297, 725, 383, 88, none in catalytic motifs), with motifs generally intolerant to mutations except Exo II and C motifs (except for the catalytic Asp residues). We did not observe many final-library positions strongly mutated to non-parental amino acids except for the one introduced from the 2M variant and a few others such as positions V63, Q91 (in the uracil binding pocket), L453, F750 or G751.

Beyond literature-known sites, we identified novel enrichments by tracking frequencies of non-parental mutations pre- and post-evolution **(Figure 6b)**. Most positions depleted, in accordance with previous observations on the evolution of other enzymes by DNA shuffling^50^ explaining that pure recombination outperforms single point mutations to obtain functional variants. However, G428E (small to negatively charged, BLOSUM: -2), K502N (positively charged to polar, BLOSUM: 0), K714E (positively to negatively charged, BLOSUM: 1), and R715M/S (positively charged to aliphatic or small polar, BLOSUM: -1) reached >1% in the final library and were strongly enriched. None of them was strongly nonconservative as shown by a minimum BLOSUM score of -2. K502 marks motif B’s end.

Shannon entropy values were mapped onto the structure of Kod DNA polymerase. Positions exhibiting the highest entropy so the highest mutational tolerance are distributed across the structure sometimes clustering by structural proximity. For example, high-entropy residues are found on opposite faces of the α-helices in the fingers domain, as well as within the N-terminal domain, suggesting localized regions of structural flexibility or reduced functional constraints **(Figure 6c)**.

### Direct coupling analysis enables protein scoring

Analysis of the segment dynamics and point mutational rate along the evolutionary trajectory provides local information, but makes it difficult to capture covariation between positions and, therefore, co-evolving positions. To address this, we used a semi-physical Potts model based on the joint occurrence frequencies of amino acids (*a_i_*, *a*_j_) at positions *i* and *j*)^51^.This model yields a mean-field Direct Coupling Analysis (mfDCA) which is an energetic description of the sequence alignment, in which the probability of observing a given sequence of length *L* (*a*_1_, *a*_2_, …, *a_L_*) is expressed as the exponential of an energy term given by the sum of local fields ℎ*_i_*(*a_i_*) and pairwise couplings *J_i_*_j_(*a_i_*, *a*_j_). Lower energy corresponds to a higher probability of observing the sequence. For each pair of positions, the coupling strength can be summarized as a direct information (DI) score, a mutual-information-derived quantity computed from the Potts-model probabilities rather than from raw frequencies. This provides a complete network of direct information coupling strengths between all position pairs **(Figure 7a)**.

**Figure 7.**
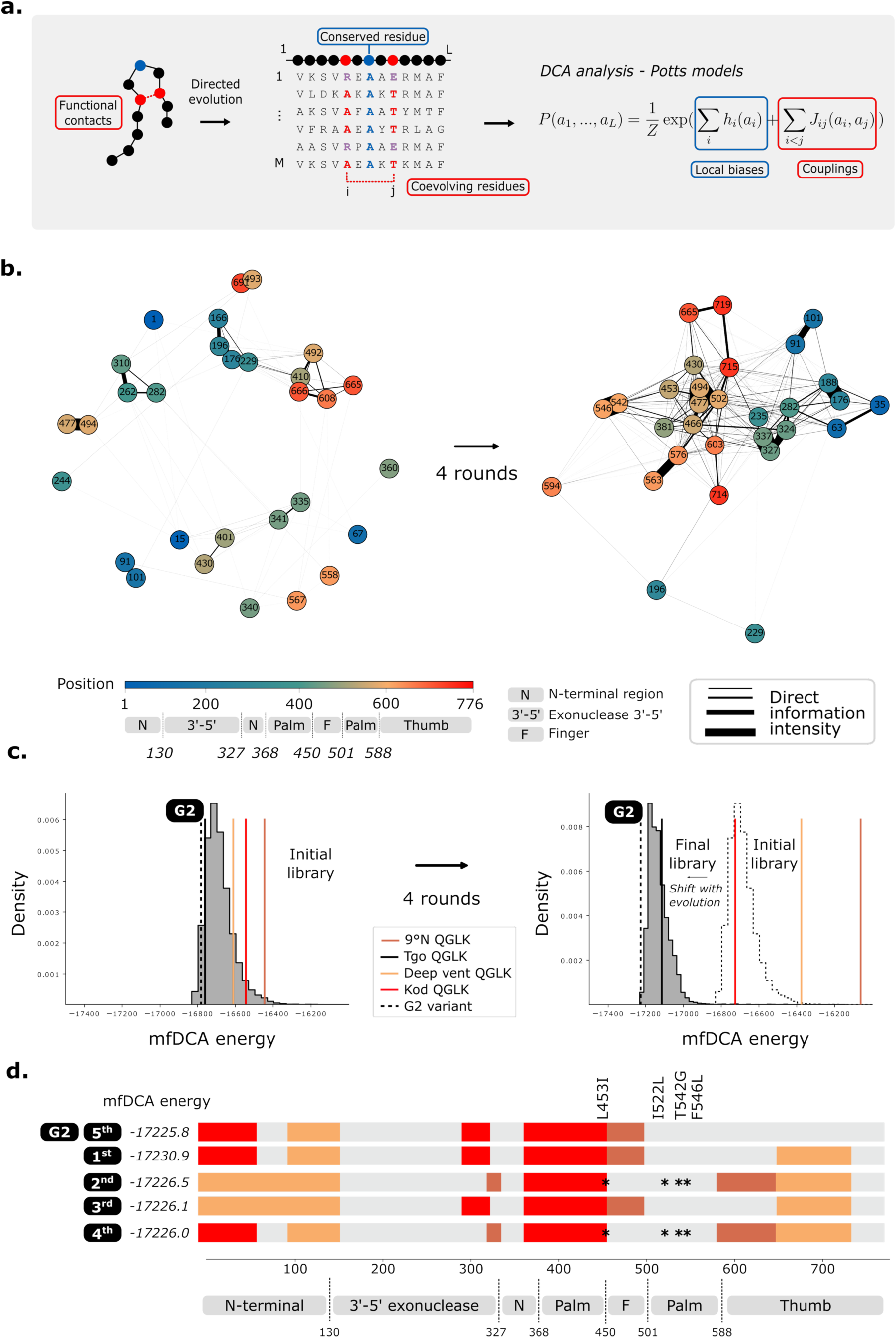
Coevolutionary analysis and Potts-model scoring of polymerase libraries. **(a)** Evolutionary constraints are inferred from MSA by analysing patterns of amino-acid conservation and co-variation. Residue-residue couplings produces correlated columns in MSA of homologous proteins. The MSA can be modelled using a Potts-model energy framework defined by inferred site-specific fields h_i_ (local biases) and pairwise coupling terms J_ij_. **(b)** A direct-information (DI) network is inferred from the initial and final MSA and nodes with the highest betweenness centrality are identified. The initial library shows weak couplings and limited interconnectivity whereas the couplings in the final library are strengthened. Couplings are particularly stronger between positions close in the sequence consistent with the recombination strategy based on relatively independent sequence blocks. **(c)** Histogram of energies assigned by the inferred Potts model to each sequence. Lower energies correspond to higher apparition probabilities. Across directed evolution, inferred energies shift towards lower values, indicating enrichment of more probable sequences according to the model. The model also distinguishes parental sequences from the evolved variant G2, assigning G2 a lower energy than the parents. **(d)** Sequences of the five lowest energies among the analysed sequences in the last library. Variant G2 ranks as the fifth. Its architecture is comparable to the four lower-energy variants, but unlike them it shows no fragmentation of the thumb domain. Segment junctions are shared across these top five variants, with strong conservation of the palm domain and the N-terminus.

To identify the most critical nodes in this network, we focused on positions with the highest betweenness centrality, i.e. those that most frequently act as bridges between strongly coupled clusters of positions^52^. Visualization of the coupling network before and after directed evolution **(Figure 7b)** shows that some strongly coupled positions at the start of evolution remain strongly coupled, such as positions 477 and 494 in motif B. This interaction becomes central in the final network by recruiting position 502 which is one of the sites whose mutation rate increases during evolution so that together they form the core of a dense interaction subnetwork. Additional strong couplings emerge during evolution, notably between positions 546 and 542, two sites previously implicated in the activity of variants TGLLK and 2M. By contrast, the second position highlighted in the 2M study, 594, is present in the final network but appears only weakly connected to the rest of the library, in line with its strong depletion during evolution **(Figure 6b)**. Other position pairs also show strong coupling, such as 563-576, 327-337, 176-188, and 91-101 which are all separated by a characteristic distance of around 10-13 residues. The network further reveals that coupled positions tend to be proximal along the sequence. Positions in the N-terminal domain couple predominantly among themselves, as do positions in the palm or thumb domain. This sharp separation between domains may either be explained by, or itself promote, homologous recombination, which (as shown above) tends to swap entire domains or subdomains between parental polymerases from different species **(Figure 5c)**.

The models inferred for the initial and final libraries allow us to score each sequence, including the four parents and our variant of interest G2. Over the course of evolution, the mean energy score (mfDCA score) decreases, with the histogram shifting towards lower energies, i.e. towards more probable sequences. The parental sequences cluster relatively closely in energy in the initial library, whereas in the final library only the parent Tgo_QGLK clearly stands out. In both libraries, the variant of interest G2 has a lower energy score than all parent sequences. In the final library, it ranks as the fifth-best variant **(Figure 7c)**. Focusing on the five top-scoring sequences **(Figure 7d)**, shows that G2 is distinguished from the other four by having a palm and thumb domain derived exclusively from Tgo, while for the other variants are a mosaic of several parents. Segment patterns are shared between variants such as the C-terminal region of 1^st^ and 3^rd^ but not the N-terminal region or the opposite for 1^st^ and 4^th^. Some regions are very conserved for instance, the 3ʹ-5ʹ exonuclease domain is always from Tgo, and similarly, the central part of the thumb domain is systematically derived from DV. Two of these top variants carry point mutations, notably the previously undescribed conservative L453I substitution and the I522L, T542G, and F546L mutations originating from the TGLLK and 2M variants.

### DCA scoring correlates with experimental fitness

To assess whether mfDCA energy scores correspond to experimental fitness, we computed a fitness score based on the log10 enrichment ratio between variant frequencies in the final and initial libraries. Plotting mfDCA energy against experimental fitness reveals a general trend: lower mfDCA scores associate with higher experimental fitness. However, the correlation is weak rather than strong (**Figure 8a**, Spearman correlation coefficient ρ = -0.36) as already observed in previous epistasis study^53^.

**Figure 8.**
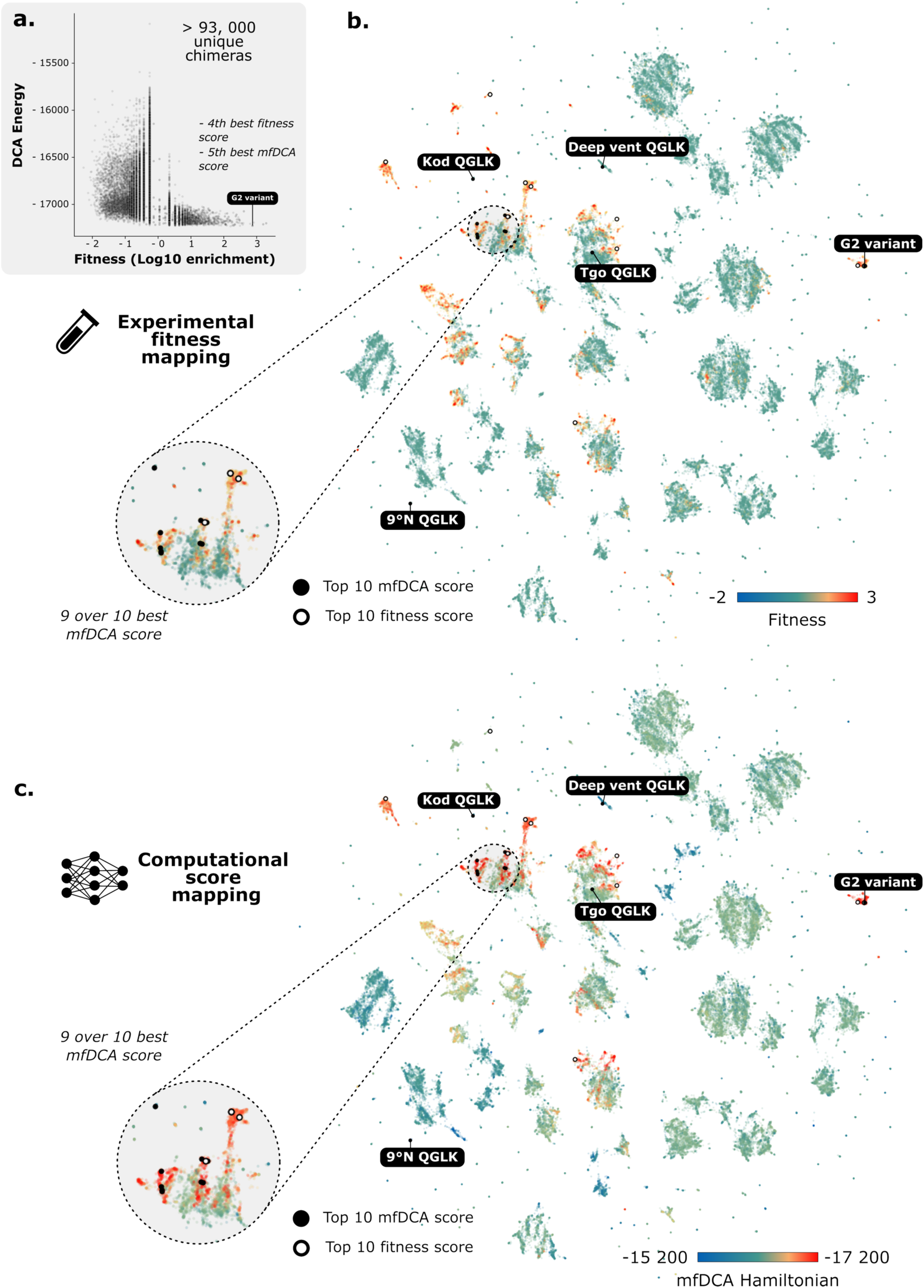
Fitness landscapes from protein language-model embeddings and comparison of experimental and mfDCA-predicted fitness. Unique chimeric sequences were embedded using a protein language model (ESM2-8M) and UMAP projected into two dimensions. Each sequence was annotated with either its experimental fitness estimated from variant enrichment between the initial and final libraries or its predicted fitness (mfDCA-derived energetical score). **(a)** The relationship between experimental and predicted fitness was assessed. Overall correlation is weak (Spearman ρ = -0.36) but the highest experimental-fitness variants tend to correspond to low mfDCA energies. This is exemplified by variant G2, which is both well scored by mfDCA and exhibits high experimental fitness, consistent with its strong enrichment in the final library. **(b)** The enrichment-based fitness landscape reveals that the highest-fitness variants are distributed across the sequence space. A zoomed region highlights the dense cluster of variants exhibiting maximal experimental fitness. Parental polymerases are part of isolated clusters and, as expected, are not associated with high experimental fitness. The cluster containing variant G2 is among those with the highest experimental fitness. **(c)** Hamiltonians derived from DCA are mapped onto the same embedded sequence space. The zoomed region shows that the fitness prediction (with 9 over 10 top mfDCA scores) is comparable with the experimental pattern. Some areas are predicted to be strongly unfavourable (high Hamiltonian), notably around the 9°N-derived region.

All unique variants were projected into a 2D embedding space using sequence representations from the ESM2-8M protein language model^54^ **(Figure 8)**. This yields fitness landscapes highlighting regions of interest strongly selected during evolution, with clusters showing local fitness maxima. All sequence clusters are separated by fitness valleys which are not explored during evolution. The four non-shuffled parents occupy small peripheral clusters which shows that few sequences of the libraries are unshuffled. There is one exception among the parents for Tgo which is part of a large cluster. This overrepresentation of Tgo-derived sequences is in accordance with its better mfDCA scoring in **Figure 7**. The top 10 experimental fitness variants are far more dispersed across sequence space than the top mfDCA hits. The G2 cluster shows good experimental and computational fitness but is shifted regarding the entire evolution. In this cluster, G2 is the sole member with a very strong (top 10) mfDCA energy score. The other nine top mfDCA variants (excluding G2) cluster in an intermediate region between Tgo, Kod and 9°N parents. This subregion shows three main enrichment clusters that are captured by both the experimental and computational fitness estimators. Comparing mfDCA and experimental fitness landscapes reveals many overlapping high-fitness zones. **(Figure 8b,c)**.

## Discussion

DNA-dependent DNA polymerases, despite their inherent stability and fidelity, are intrinsically optimized for natural deoxyribonucleotide substrates, making them difficult to reprogram for RNA synthesis and particularly for the incorporation of highly modified nucleotides. Although the rational design has identified key control points that enable limited RNA synthesis^22,55^, these modifications alone are insufficient to support the diverse chemistries required for therapeutic applications, including those under consideration for next-generation aptamers and small interfering RNAs.

Herein, using UTP as a model substrate, we describe the directed evolution of a DNA polymerase capable of synthesizing hypermodified RNA bearing modifications at both the ribose (position C2ʹ) and the nucleobase (position C5). Our aim was to expand the chemical scope available for RNA therapeutics by incorporating bio-inspired 2’-*O*-methyl glyconucleotides into RNA strands. Besides enhancing RNA stability toward endo- and exonucleases using C2’ modified RNA^56,57^, the targeted improvement relies primarily on the enrichment of the nucleotide library by original nucleotide bricks featuring a glycosyl-triazolmethyl appendage at C5. To modulated RNA properties such as affinity or stability, it was of interest to envision a diversity of glycan appendages differing in terms of nature and chain length. The efficient chemo-enzymatic synthesis of these 2’-masked glycosyl-modified RNA is unique to our knowledge. A modular approach to glycosylated 2’-OMe UTP is described that enables access to a large diversity glycosylated of 2’-OMe NTP analogues. Achievements on uridine, initially selected as a model nucleobase with the aim of subsequently extending this approach to other bases, pave the way to multiple variations relying on the four basic nucleotides combined to the large chemically available glycospace.

The evolved variant, G2, incorporates up to 12 modified nucleotides in a 38-nt RNA, including 2ʹ-*O*-methyl substitutions and complex glycosylations ranging from monosaccharides to maltotriose **(Figure 3b)**. This level of chemical diversity enables access to glycosylated RNA architectures with strong potential for enhanced stability, tunable pharmacokinetics, and selective recognition of glycosylated targets, extending beyond prior efforts that were largely limited to glycosylated DNA aptamers^58^. Moreover, while selected by screening a large family of variants against one single modified nucleotide substrat (5-maltosyl-2’-*O*-methyl-UTP), the selected variant was shown to be also highly relevant for the incorporation of a broad range of substrates specifically modified on the sugar moiety, from pseudouridines to chemically unnatural HNA nucleotides **(Figure 4)**. The 5-ethynyl-2’-*O*-methyl-UTP, which served as precursor to all glycosylated analogues, was also a very good substrate. This paves the way to strategies whereby RNA glycosylation by site selective click chemistry could be envisioned post RNA elongation instead of by incorporation of glycosylated nucleotides.

Directed evolution can be viewed as an exploration of sequence space in the vicinity of parental enzymes to identify variants with improved functional activity. This process is markedly enhanced by gene shuffling and homologous recombination^50,59^, which enable the rapid assembly of diverse sequence combinations. In our system, homologous recombination generated a substantially richer mosaic of parental segments than those predicted purely on the basis of idealized models of DNA shuffling. Rather than producing sharply defining fragments at designed crossover points, we observed a broad distribution of segment lengths **(Figure 5c)**, consistent with PCR-driven processes such as partial extension, template switching, and occasional point mutations. This dynamic segment behavior contributes more strongly to functional diversification than random mutagenesis, as evidenced by the depletion of mutations not inherited from the four parental polymerases. Notably, the previously described 2M mutations^29^, considered essential for 2ʹ-modified nucleotide incorporation, are only partially retained in the evolved library (even if present at more than 30% in the initial library) and are absent from several high-fitness variants. Of the two residues, only T542G is consistently conserved, whereas K594A is frequently lost, suggesting that their previously strong impact reflect additive contributions rather than strict epistatic coupling. **(Figure 6b)**

Segment dynamics are most pronounced at the boundaries between structural domains, including the N-terminal region, 3ʹ-5ʹ exonuclease, palm, fingers, and thumb subdomains. In contrast, the strongest epistatic couplings are largely confined within individual domains (**Figure 7b**), indicating that substrate specificity is governed primarily by intra-domain interactions rather than connections across domains. These observations underscores the importance of domain-level recombination between polymerases from different species for acquiring non-native functions, which is consistent with prior work demonstrating that recombination accelerates enzyme evolution by combining distant, pre-adapted structural modules^27,60^. Notably, the final G2 variant carries no additional point mutations beyond those obtained by natural diversity and is assembled from nine recombined segments, underscoring the dominant role of segmental reassortment in accessing new functional states.

Projection of the evolved sequences into a two-dimensional protein-language-model embedding reveals that homologous recombination explores sequence space more broadly than point mutagenesis alone **(Figure 8)**. While point mutations sample local neighborhoods, recombination enables large, non-local jumps that reorganize domains or multi-residue segments, giving rise to distinct functional clusters. Consistent with prior comparisons of DNA shuffling and error-prone PCR, recombination libraries are expected to contain a higher fraction of functional variants than error-prone PCR libraries, which predominantly generate deleterious mutations^61,62^. These results illustrate how recombination can efficiently traverse rugged fitness landscapes by reassembling pre-validated structural modules rather than relying solely on incremental single-point changes. This behavior is analogous to the Swendsen and Wang algorithm used to simulate phase transitions in 2D or 3D Ising spin systems, in which clusters of spins are flipped collectively rather than individually, enabling more efficient exploration of the energy landscape in Monte Carlo simulations^63^.

To generalize sequence behavior across the evolutionary trajectory, we inferred a semi-physical Potts (mfDCA) model that assigns an energy-like score to each sequence, commonly used as a proxy for fitness in natural protein families^64^. In our recombination-dominated libraries, the mfDCA score remains partially informative, as it distinguishes the active G2 variant from parental sequences. However it correlates only weakly with the experimentally measured fitness based on enrichment across screening rounds **(Figure 8a)**. This discrepancy likely arises from multiple factors. First, the experimental fitness metric is an indirect readout of catalytic activity and is affected by sequencing depth or bacterial growth noise. Second, Potts models capture only pairwise residue couplings, whereas our analysis indicates that evolution proceeds through higher-order interactions, including coordinated exchanges of entire domains or multi-residue segments that lie beyond a pairwise representations. In silico evolution for such systems will require generative models capable of capturing higher-order and segment-level epistasis, rather than relying on pairwise representations alone.

Together, these results demonstrate that engineering polymerases for XNA synthesis, particularly for densely modified glyco-RNAs, is driven very effectively by recombination of parental sequences. Achievements in the context of uridine analogues, are expected to be transposable to other ribonucleosides. A route to glycan-encoded nucleic acids and chemically expanded therapeutics is established.

## Methods

### Synthesis of 5-glycosyl-2’-*O*-methyl-UTP

The 5-glycosyl-2’-*O*-methyl-UTP analogs (**5-8**) were synthesized in two stages (**Supplementary Figure 1**, full procedures **in Supplementary Data**). First, the alkyne building block 5-ethynyl-2’-*O*-methyl-UTP (**4**) was assembled from D-ribose in nine steps through key intermediates **2** and **3**, employing protocols for methyl ribofuranoside formation and selective O-methylation^65,66^, Sonogashira cross-coupling to introduce a 5-ethynyl group onto 5-iodouracil^67,68^, Vorbrüggen nucleosidation, and Ludwig-Eckstein triphosphorylation in the final step. Second, azidoethyl glycosides of glucose, arabinose, maltose and maltotriose (**26-29**) were prepared from the parent saccharides by acid-catalyzed Fischer glycosidation with 2-bromoethanol, followed by S_N_2 azidation with sodium azide. Preparative RP-HPLC was employed to isolate the α-configured anomers. The glycosylated triphosphates **5-8** were then obtained by copper(I)-catalyzed azide-alkyne cycloaddition (CuAAC, CuI, DMF/H_2_O, 25 °C, 4 h) between **4** and the respective azidoethyl glycosides, and purified by anion-exchange HPLC on a DNAPac PA-100 column. Additionally, 2’-*O*-(2-methoxyethyl)-UTP (**21**) was prepared from commercial 2’-*O*-(2-methoxyethyl)uridine by sequential DMT protection, O-acetylation, DMT removal, and Ludwig-Eckstein triphosphorylation. All compounds were fully characterized by ^1^H, ^13^C, and ^31^P NMR spectroscopy when appropriate, high-resolution mass spectrometry (HRMS-ESI) and MALDI-TOF for triphosphates(**Supplementary Data**).

### Design and construction of libraries

Genes encoding B-family DNA polymerases from *Thermococcus gorgonarius* (Tgo; GenBank: KP682507), *Thermococcus sp. 9°N* (9°N; GenBank: KP682506), *Pyrococcus sp. GB-D DV* (DV; GenBank: KP682509) and *Thermococcus kodakarensis* (Kod; GenBank: KP682508) were designed with 3ʹ-5ʹ exonuclease-deficient mutations (D141A and E143A) and previously described RNA polymerase activity mutations (V93Q, Y409G, A485L and E664K)^22,24,26,28^. Eleven regions of high sequence identity (approximately 6-12 amino acids in length) were identified among these scaffolds and codon-swapped to generate perfectly homologous segments suitable for DNA shuffling. The resulting Tgo-QGLK, 9°N-QGLK, DV-QGLK and Kod-QGLK gene variants were obtained as synthetic gBlocks (Integrated DNA Technologies). Shuffled polymerase libraries were obtained after screening for a new function described previously^27^. The region corresponding to the secondary steric gate (2M variant), identified for B-family polymerases^29^, was subsequently replaced by Gibson assembly. This replacement fragment, designed on the Tgo background and carrying mutations I521L, T541G, F545L and K592A, was synthesized as a separate gBlock.

To construct the shuffled library vector, the pGDR11 backbone was amplified from 10 ng of pGDR11-C28 Round 3 evolution plasmid using Q5 high-fidelity DNA polymerase (New England Biolabs) with 1× Q5 reaction buffer, 1 μM forward primer, 1 μM reverse primer, 0.4 mM dNTPs and 1 μL Q5 polymerase in a 50 μL reaction. Thermocycling conditions were: 95 °C for 2.5 min; 30 cycles of 95 °C for 30 s, 68 °C for 45 s and 72 °C for 4 min; followed by a final extension at 72 °C for 1 min. This PCR generated the complete pGDR11 vector backbone containing the shuffled polymerase library, except for the segment to be replaced by the 2M steric gate gBlock. The PCR product was treated with DpnI (New England Biolabs, R0176S) to remove template plasmid, resolved on an agarose gel, purified using a commercial gel extraction kit and quantified by UV absorbance at 260 nm.

For Gibson assembly, 100 ng of the Tgo:2M gBlock and 100 ng of the purified vector backbone were combined with 2× Gibson Assembly Master Mix (New England Biolabs) in a 20 μL reaction and incubated at 50 °C for 1 h. A 4 μL aliquot of the assembly mixture was then transformed into 50 μL chemically competent NEB 5-alpha *E. coli*. Cells were plated on LB-agar containing carbenicillin (50 μg.mL^-1^) and incubated overnight at 37 °C to obtain single colonies. Individual colonies were screened by colony PCR.Plasmids from correctly sized clones were purified and subjected to whole plasmid sequencing to verify the integrity of the recombined polymerase genes. The validated Gibson-assembled library was transformed into NEB 5-alpha cells, grown on multiple plates, and colonies were scraped to prepare a pooled plasmid library, which was purified, quantified by UV absorbance and stored at -20 °C for subsequent rounds of transformation and directed evolution. The pGDR11 construction obtained is an IPTG-inducible *E. coli* expression vector with a T5/lac promoter regulated by plasmid-encoded LacI.

### Encapsulation of libraries microfluidics

A total of 100 ng plasmid library was transformed into 40 μl XL10-Gold ultracompetent *E. coli* cells (Agilent Technologies). SOC-recovered cells were used to inoculate 50 mL LB-carbenicillin (50 μg.mL⁻¹) and grown overnight at 37 °C, 250 rpm. This culture was diluted 1:100 (v/v) into 50 mL fresh LB-carbenicillin and grown to OD_600_ ≈ 0.6 before induction with 1 mM IPTG at 37 °C for 3 h.

Cultures (1 mL by 1 mL) were pelleted at 3,500 × g for 5 min, washed once in 500 μl 1× ThermoPol buffer (New England Biolabs), and resuspended in 500 μL 1× ThermoPol buffer to OD_600_ = 0.05 to enable encapsulation at occupancies of 0.1 cells per droplet according to a Poisson distribution. Cell suspensions were mixed with polymerase activity assay reagents: 1 μM 5ʹ-Cy3-labeled self-priming hairpin template (30merHP.V2), 2 μM 3ʹ-Iowa Black quencher (QP08.Iowa Black), 100 μM each dATP, dGTP, dCTP and glycosylated 2ʹ-OMe-UTP, and 1× ThermoPol buffer.

Aqueous and oil phases were loaded into 1.5-mL tubes. Tygon tubing was inserted through drilled tube caps and sealed: one line going to an SMC ITV0011-2UMS digital pressure regulator and another submerged for fluidic input. Water-in-oil single emulsions were generated in custom polydimethylsiloxan (PDMS) microfluidic chips using flow-focusing geometry where aqueous polymerase activity assay mix was sheared by HFE-7500 fluorinated oil (3M) containing 1% (w/w) Pico-Surf (Dolomite Microfluidics). Flow pressures were controlled by custom LabVIEW software (National Instruments). Droplet production was tuned to ≈20-22 μm diameters at 30-35 kHz (≈10^7^ droplets h⁻¹). Emulsions were collected under mineral oil in 1.5-mL tubes, incubated at 70 °C for 5 min to lyse cells, then at 55 °C for 6 h to enable polymerase extension reactions.

### Fluorescence-Activated Droplet Sorting and plasmid recovery

Post-incubation droplets were sorted using a FADS microfluidic chip as described in^48,69^. A 552-nm laser illuminated the droplets through a ×20 objective. Cy3 emission passed a 562-nm quad-band dichroic, long-pass dichroics and was detected by a photomultiplier tube (PMT) after a 563 nm band-pass filter. A high-speed camera (35,000 frames.s^-1^) imaged blue-light-excited droplets to avoid Cy3 spectral overlap. PMT signals were processed by field-programmable gate array (FPGA) hardware running custom LabVIEW software. Droplets exceeding fluorescence threshold were deflected by dielectrophoretic force into the collection channel by a liquid indium electrode (50 kHz, 50% duty cycle, 60 μs square-wave pulse, amplified to 600 V).

Plasmid DNA from sorted droplets was extracted using Pico-Break (Sphere Fluidics) according to the manufacturer instructions after vortexing and centrifugation (2,000 g, 60 s). The top aqueous layer was recovered and the bottom layer was re-extracted with nuclease-free water. Fractions were pooled, and concentrated with Zymo DNA Clean & Concentrator IC column (500 μL wash buffer ×2; elution in 20 μL nuclease-free water). The purified, concentrated plasmid was transformed into XL10-Gold ultracompetent *E. coli* for library propagation and the next evolution round.

### Polymerase activity screening

Libraries were transformed into NEB-5-apha^TM^ *E. coli* (C2987H) and plated onto ampicilin LB agar plate (100 µg.mL^-1^). Single colonies were transferred into 96-well plates (Eppendorf) containing 1 mL LB-ampicilin medium and cultivated overnight at 37 °C with shaking at 800 rpm using an Eppendorf ThermoMixer incubator. 10 μL of overnight culture was inoculated into fresh 1 mL ampicilin LB medium (100 µg.mL^-1^), and cells were grown to OD_600_≈0.6. Expression was induced with 1 mM IPTG, and cultures were incubated at 20 °C, 800 rpm overnight. Cells were pelleted by centrifugation (3,200 × g, 25 °C, 20 min), resuspended in 50 μL 1× ThermoPol buffer (20 mM Tris-HCl pH8.8, 10 mM (NH_4_)_2_SO_4_, 10 mM KCl, 2 mM MgSO_4_,0.1% Triton X-100), and transferred to 96-well PCR plates. Samples were incubated at 75 °C in an Eppendorf ThermoMixer for 30 min, cooled on ice for 15 min, and centrifuged (3,200 × g, 25 °C, 1 h). To assess polymerase activity, reactions were assembled to a final volume of 10 μL containing 1× ThermoPol buffer, 0.5 μM DNA template, 0.5 μM 5’-FAM-labelled primer, 0.25 mM NTPs (incuding 5-glycosyl-2’-*O*-methyl-UTP), and 1 μL lysate, and incubated at 60 °C for 60 min. Reactions were terminated by the addition of two volumes of stop buffer (10 mM EDTA, 98% formamide, 1 mg.mL^-1^ bromophenol blue, 10 μM unlabelled complementary primer). Prior to analysis, samples were preheated at 95 °C for 10 min, resolved by denaturing 18% urea-polyacrylamide gel electrophoresis, and visualized using a Typhoon FLA 9000 biomolecular imager (Cytiva). The selected clones were reevaluated by verifying their fidelity through the elimination of 5-glycosyl-2’-*O*-methyl-UTP in order to check whether overextension was observed.

### Polymerase purification

NEB-5-apha^TM^ *E. coli* competent cells were transformed with the pGDR11 PolB expression plasmid (C28, C28-2M and G2 variant of interest). A selected colony was inoculated into LB medium and cultured at 37 °C. The preculture was diluted 1:100 and protein expression was induced by adding 1 mM IPTG when the culture reached OD_600_ 0.6, followed by an overnight incubation of the 2 L culture at 20 °C. Cells were harvested by centrifugation at 3,300 × g for 30 min, and the pellet was resuspended in 40 mL lysis buffer (50 mM Tris-HCl pH 7.5, 300 mM NaCl, 1 mM DTT, 0.05% Triton X-100). One tablet of protease inhibitor cocktail (Thermo Scientific) and 1 μL of benzonase nuclease (Millipore) per 30 mL of lysate were added. Cells were disrupted by sonication with five cycles (60s on - 60s off). Lysates were clarified by centrifugation at 30,000 × g for 30 min. Polyethylene glycol (PEG) 8000 was added to the supernatant to reach a 10% final concentration. The solution was heated at 70 °C for 15 min, cooled for 5 min at room temperature, and then on ice for 15 min. A white/yellow precipitate formed in the lysate, which was removed by centrifugation at 15,000 × g for 40 min. The supernatant was diluted to adjust the salt concentration to approximately 90 mM before HiTrap Heparin purification (Cytiva), performed using a gradient between buffer A (100 mM NaCl, 50 mM Tris-Cl pH 7, 1 mM DTT, 0.05% Trion X-100) and buffer B (2M mM NaCl, 50 mM Tris-Cl pH 7, 1 mM DTT, 0.05% Trion X-100). Eluted fractions were collected and concentrated.

Protein purity was analyzed by SDS-PAGE using 4-15% or 4-12% gels with a Precision Plus Protein molecular weight ladder (BioRad). Enzyme samples were concentrated with Amicon Ultra centrifugal filters (30 kDa MWCO, Merck), flash frozen in liquid nitrogen, and stored at -20 °C.

### Kinetic analysis

G2 and C28 polymerases were purified as described above. To quantify their kinetics for incorporation of modified UTP, primer-extension reactions were analyzed by capillary electrophoresis. Reactions (10 µL) contained 1× ThermoPol buffer, 0.5 µM 5ʹ-ATTO488-labelled primer annealed to a DNA template, 0.25 mM nucleotides (dNTPs, NTPs with and without 5-glycosyl-2’-*O*-methyl-UTP), and 1 µM purified polymerase (G2 or C28), and were incubated at 60 °C for 10 seconds, 30 seconds, 1, 5, 15 and 30 minutes. Each time course was performed in duplicate. Reactions were quenched by adding 40 µL 50 mM EDTA and then diluted 1:100 in miliQ water. Samples were submitted to Microsynth for fragment-length analysis by capillary electrophoresis (Filter set : DS-33_G5, Size standard : Genescan 120 LIZ). Traces were aligned and peak areas were used to calculate the fraction of fully extended primer relative to partially extended and unreacted species, yielding the extent of product formation at each time point.

### Polymerases activity assays

Polymerases G2, C28 and C28-2M were purified as described above. Vent(exo^-^) DNA polymerase (New England Biolabs, M0257S) was diluted by 10 before use. Polymerase activity was assayed in 10 μL reactions containing 1× ThermoPol buffer (20 mM Tris-HCl pH 8.8, 10 mM KCl, 10 mM (NH₄)₂SO₄, 2 mM MgSO₄, 0.1% Triton X-100), 0.5 μM DNA template, 0.5 μM 5ʹ-FAM primer, 0.25 mM each NTP (including 5-glycosyl-2ʹ-OMe-UTP), and 1 μM polymerase (or 1 μL diluted Vent(exo^-^)). Reactions were incubated at 60 °C for 30 min. Reactions were quenched by adding two volumes of formamide stop buffer (10 mM EDTA, 98% formamide, 1 mg.mL^-1^ bromophenol blue, 10 μM unlabelled complementary primer) and heated at 95 °C for 10 min prior to loading. Products were resolved by denaturing 18% urea–polyacrylamide gel electrophoresis in 1× TBE and visualized on a Typhoon FLA 9000 biomolecular imager (Cytiva).

### Libraries tagging and sequencing

A Q5 PCR was performed to introduce 32-nt semi-structured unique molecular identifiers (UMIs) flanking the region of interest such that all reads sharing the same pair of UMIs derive from a single input molecule. The product obtained was gel purified with NucleoSpin^TM^ Gel and PCR Clean-up (740609.50). PCR products were recloned into pGDR11 using 2X NEBuilder^TM^ HiFi DNA assembly master mix (100 ng insert : 100 ng backbone, E2621S). The tagged library was transformed into NEB 10-beta electrocompetent cells (C3020) cells by electroporation and transferred into 100 mL LB medium supplemented with ampicilin (100 µg.mL^-1^) for overnight growth at 37 °C. Following library amplification, PCR was performed to generate sequencing-ready DNA using the following conditions: Q5 PCR buffer 1x, (NEB B9027), GC Enhancer 1x (NEB B9028), 200 μM dNTPs (N0447), 200 nM forward primer, 200 nM reverse primer, 1-5 ng tagged plasmid DNA, and 10 units Q5 polymerase (M0491S). Cycling conditions were as follows: initial denaturation at 98 °C for 2 min; 27 cycles of 98 °C for 10 s, 68 °C for 30 s, and 72 °C for 3 min. 500 μL reaction mix was made and column purified using the MinElute kit (Qiagen 28004). A low cycle number combined with prolonged extension times was used to minimize PCR-mediated chimera formation between variants, which would otherwise distort quantitative analysis of variant representation in the library.

Sequencing libraries were prepared using the Ligation Sequencing Amplicons V14 protocol (SQK-LSK114). Sequencing was stopped once the sequencing read count reached 50-fold sample coverage (assuming 10^5^ variants and 2 kb amplicon lengths) corresponding to >10 Gb sequencing data and around 5 million reads.

### Nanopore sequencing denoising

The following variant pipeline adapts previously described denoising long-read strategies^70–73^ to much larger datasets (here, several gigabases corresponding to millions of 1.9-kb reads). Briefly, raw ONT signals are basecalled with Dorado (SUP v5), and reads are aligned to the consensus sequence of parental polymerases using Minimap2^74^ (keeping PCR duplicates) as it is the base of the denoising method. UMI sequences at both ends of each read are extracted from the alignments and used to cluster reads with CD-HIT-EST^75^ at 91.5% identity, thereby grouping reads derived from the same molecule. Clusters supported by fewer than five reads are discarded. Within each remaining cluster, variants relative to the reference are called with freebayes^76^ and restricted to single-allele substitution calls that pass filters on allele fraction (>60%) while sites outside the coding region, positions with ambiguous reference bases, indels and multi-allelic calls are discarded, so that only high-confidence mutations consistently observed across all reads of a cluster are retained as true variants.

The sequencing experiments are summarized in **Supplementary Table 3**. The table reports, for each sequenced library, the library name, the number of raw reads before clustering, the number of reads aligned to the reference, the number of clusters (defined as unique UMI pairs with ≥5 supporting reads) and their mean cluster size, the number of unique variants recovered, and the number of unique variants with no premature stop codons together with the fraction of reads carrying stop codons in the library. Each amplicon is typically covered by an average of ≃10 reads (mean cluster size across libraries), providing multiple independent observations of each sequence and enabling robust consensus calling and accurate per-variant depth estimates.

### Sequence data analysis

All unique variant sequences from initial and final libraries obtained from Oxford Nanopore sequencing were aligned with the four parental B-family polymerases (Tgo-QGLK, 9°N-QGLK, DV-QGLK, Kod-QGLK) with Clustal-Omega^77^. This multiple sequence alignment (MSA) is splitted per library to obtain MSA with the same lengths. Protein sequences were filtered to remove sequences containing an internal stop codon, which are most likely inactive. To identify the parental segments, the RDP2 code from^78^ was adapated. Local identity (with each parents) was scored in 10 amino acids sliding window and Gaussian-smoothed to reduce noise effects leading to a first score matrix. Parental segments were identified by computing maximum (depending of the previous parent) cumulative similarity scores across positions. The score favored continuation of a parent by adding a switch penalty (=0.1) and we enforced a minimum segment length of 5 amino acids. Positions with no parental origin were identified as non-natural mutations. Parental proportions per position were visualized as heatmaps **(Figure 5c)**.

For each alignment position *i*, the Shannon entropy was computed from the empirical amino-acid frequency distribution *p_i_*(*a*).

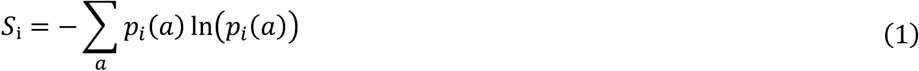

The obtained MSA were used directly for mean-field Direct Coupling Analysis (mfDCA). Calculations were performed in Python using the py-mfdca implementation (Ref) of the Matlab method published by Morcos et al.^51^ with default parameters. The DCA model assigns a probability to a sequence of length L based on the observed single-site *a_i_* and pairwise 7*a_i_*, *a*_j_8 statistics of amino acids across positions.

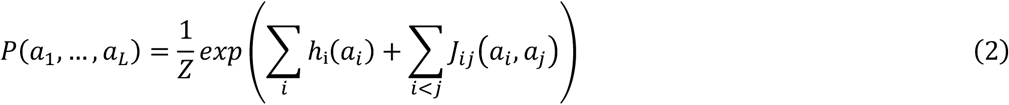

From the inferred model parameters, the direct information between residue pairs was computed to quantify direct coevolutionary couplings with the same py-mfdca package.

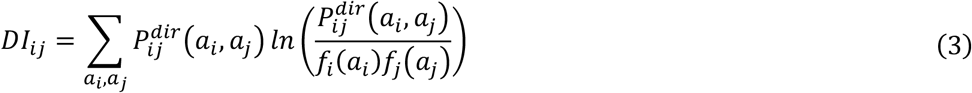

Finally, networks were constructed from these couplings (python package : networkx), and nodes were ranked by their betweenness centrality, defined as the fraction of all shortest paths between any pair of nodes that pass through a given node. Nodes with the highest betweenness act as major intermediates to connect amino acids clusters and were identified as key communication hubs (**Figure 7**).

Sequence counts were obtained for each variant across the libraries, and a frequency was assigned to each variant at its final appearance in the evolutionary trajectory. Enrichment was then calculated for each variant relative to the initial, non-selected library lVE001. The fitness score was expressed as the log10 of this enrichment, as previously described^79^ (**Figure 8**).

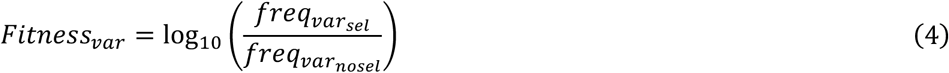

All the unique sequences were then isolated and encoded with the esm.pretrained.esm2_t48_8M_UR50D() protein language model^54^. For the analysis, representations from the 6^th^ encoder layer were extracted. After extracting each variant’s embedding, these were reduced in dimensionality through UMAP (python package: sklearn, default settings, n_components = 2). In **Figure 8**, each variant was then colored according to the magnitude of its calculated fitness inspired from^80^ or with the estimated energy obtained from mfDCA.

## Reporting summary

Further information on research design is available in the Nature Portfolio Reporting Summary linked to this article.

## Data Availability

The sequence datasets will be made available from the authors upon reasonable request.

## Code Availability

Unique Molecular Identifier pairs were extracted from filtered alignments using our in-house tool xumi v1.0.3 deposited in https://github.com/Fravadona/xumi.

## Acknowledgements

The authors acknowledge enlightening discussions with all past and present members of the Delarue, Sauguet (IP) and Chaput (UCI) teams. We thank Piet Herdewijn for the gift of the HNA nucleotides and Simona Cocco for insightful remarks on the manuscript.

## Contributions

M.D., J.C.C. and E.V. conceived the project and designed the experiments. E.V. and V.M. evolved the enzyme. E.V., C.L. and S.L.S. performed the biochemical characterization. M.D.P and M.B.G. synthetized the nucleotides. M.H. designed and obtained the initial library. E.V, C.L., R.N. and R.S. performed sequencing experiments and analysis. M.D., J.C.C. and E.V. wrote the paper. All authors reviewed and commented on the paper.

## Ethics declarations

Competing interests: M.D., J.C.C, V.M., M.H. and E.V. have filed a patent application on the composition and activity of the G2 XNA polymerase (US64/054,572).

## References

1. Mahler, M. et al. Phage arabinosyl-hydroxy-cytosine DNA modifications result in distinct evasion and sensitivity responses to phage defense systems. Cell Host Microbe 33, 1173–1190.e9 (2025).

2. Dhara, D., Mulard, L. A. & Hollenstein, M. Natural, modified and conjugated carbohydrates in nucleic acids. Chem. Soc. Rev. 54, 2948–2983 (2025).

3. Pyle, J. D., Lund, S. R., O’Toole, K. H. & Saleh, L. Virus-encoded glycosyltransferases hypermodify DNA with diverse glycans. Cell Rep. 43, 114631 (2024).

4. Yang, Q. et al. Carboxymethylcytosine is a natural base modification and a handle for bacteriophage DNA hypermodification. Nat. Commun. 17, 281 (2025).

5. H.Gommers-Ampt, J. & Borst, P. Hypermodified bases in DNA. The FASEB Journal 9, 1034–1042 (1995).

6. Lee, Y.-J. et al. Identification and biosynthesis of thymidine hypermodifications in the genomic DNA of widespread bacterial viruses. Proc. Natl. Acad. Sci. U. S. A. 115, E3116–E3125 (2018).

7. Varki, A. Biological roles of glycans. Glycobiology 27, 3–49 (2017).

8. Flynn, R. A. et al. Small RNAs are modified with N-glycans and displayed on the surface of living cells. Cell 184, 3109–3124.e22 (2021).

9. Xie, Y. et al. The modified RNA base acp3U is an attachment site for N-glycans in glycoRNA. Cell 187, 5228–5237.e12 (2024).

10. Yi, L., Zhou, Y., Zhang, C., Lu, H. & Xie, Y. GlycoRNA research: from unknown unknowns to known unknowns. Protein Cell 17, 1–20 (2026).

11. Dalziel, M., Crispin, M., Scanlan, C. N., Zitzmann, N. & Dwek, R. A. Emerging principles for the therapeutic exploitation of glycosylation. Science 343, 1235681 (2014).

12. Janas, M. M. et al. Selection of GalNAc-conjugated siRNAs with limited off-target-driven rat hepatotoxicity. Nat. Commun. 9, 723 (2018).

13. Nair, J. K. et al. Multivalent N-Acetylgalactosamine-Conjugated siRNA Localizes in Hepatocytes and Elicits Robust RNAi-Mediated Gene Silencing. J. Am. Chem. Soc. 136, 16958–16961 (2014).

14. Werner, F. & Grohmann, D. Evolution of multisubunit RNA polymerases in the three domains of life. Nat. Rev. Microbiol. 9, 85–98 (2011).

15. Cermakian, N. et al. On the evolution of the single-subunit RNA polymerases. J. Mol. Evol. 45, 671–681 (1997).

16. Patel, P. H. & Loeb, L. A. Getting a grip on how DNA polymerases function. Nat. Struct. Mol. Biol. 8, 656–9 (2001).

17. Braithwaite, D. K. & Ito, J. Compilation, alignment, and phylogenetic relationships of DNA polymerases. Nucleic Acids Res. 21, 787–802 (1993).

18. Delarue, M., Poch, O., Tordo, N., Moras, D. & Argos, P. An attempt to unify the structure of polymerases. Protein Eng. Des. Sel. 3, 461–467 (1990).

19. Astatke, M., Ng, K., Grindley, N. D. F. & Joyce, C. M. A single side chain prevents Escherichia coli DNA polymerase I (Klenow fragment) from incorporating ribonucleotides. Proc. Natl. Acad. Sci. U. S. A. 95, 3402–3407 (1998).

20. Bonnin, A., Lázaro, J. M., Blanco, L. & Salas, M. A single tyrosine prevents insertion of ribonucleotides in the eukaryotic-type φ29 DNA polymerase 1 1Edited by A. R. Fersht. J. Mol. Biol. 290, 241–251 (1999).

21. Medina, E., Yik, E. J., Herdewijn, P. & Chaput, J. C. Functional Comparison of Laboratory-Evolved XNA Polymerases for Synthetic Biology. ACS Synth. Biol. 10, 1429–1437 (2021).

22. Cozens, C., Pinheiro, V. B., Vaisman, A., Woodgate, R. & Holliger, P. A short adaptive path from DNA to RNA polymerases. Proc. Natl. Acad. Sci. U. S. A. 109, 8067–8072 (2012).

23. Wang, G., et al. Thermophilic Nucleic Acid Polymerases and Their Application in Xenobiology. Int. J. Mol. Sci. 23, 14969 (2022).

24. Pinheiro, V. B. et al. Synthetic genetic polymers capable of heredity and evolution. Science 336, 341–4 (2012).

25. Fogg, M. J., Pearl, L. H. & Connolly, B. A. Structural basis for uracil recognition by archaeal family B DNA polymerases. Nat. Struct. Biol. 9, 922–927 (2002).

26. Gardner, A. Determinants of nucleotide sugar recognition in an archaeon DNA polymerase. Nucleic Acids Res. 27, 2545–2553 (1999).

27. Medina, E. L. et al. Rapid evolution of a highly efficient RNA polymerase by homologous recombination. Nat. Chem. Biol. 1–9 (2026).

28. Ramsay, N. et al. CyDNA: Synthesis and Replication of Highly Cy-Dye Substituted DNA by an Evolved Polymerase. J. Am. Chem. Soc. 132, 5096–5104 (2010).

29. Freund, N. et al. A two-residue nascent-strand steric gate controls synthesis of 2ʹ-O-methyl- and 2ʹ-O-(2-methoxyethyl)-RNA. Nat. Chem. 15, 91–100 (2023).

30. Kropp, H. M., Betz, K., Wirth, J., Diederichs, K. & Marx, A. Crystal structures of ternary complexes of archaeal B-family DNA polymerases. PLoS One 12, e0188005 (2017).

31. Samson, C. et al. Structural Studies of HNA Substrate Specificity in Mutants of an Archaeal DNA Polymerase Obtained by Directed Evolution. Biomolecules 10, 1647 (2020).

32. Gutfreund, C. et al. Structural insights into a DNA polymerase reading the xeno nucleic acid HNA. Nucleic Acids Res. 53, 1156 (2025).

33. Gardner, A. F. et al. Therminator DNA Polymerase: Modified nucleotides and unnatural substrates. Front. Mol. Biosci. 6, 28 (2019).

34. Zhai, L. et al. Semi-rational evolution of a recombinant DNA polymerase for modified nucleotide incorporation efficiency. PLoS One 20, e0316531 (2025).

35. Chen, T. et al. Evolution of thermophilic DNA polymerases for the recognition and amplification of C2ʹ-modified DNA. Nat. Chem. 8, 556–562 (2016).

36. Xia, G. et al. Directed evolution of novel polymerase activities: Mutation of a DNA polymerase into an efficient RNA polymerase. Proc. Natl. Acad. Sci. U. S. A. 99, 6597–6602 (2002).

37. Jestin, J.-L., Kristensen, P. & Winter, G. A Method for the Selection of Catalytic Activity Using Phage Display and Proximity Coupling. Angew. Chem. Int. Ed. 38, 1124–1127 (1999).

38. Ghadessy, F. J., Ong, J. L. & Holliger, P. Directed evolution of polymerase function by compartmentalized self-replication. Proc. Natl. Acad. Sci. U. S. A. 98, 4552–7 (2001).

39. Ellefson, J. W. et al. Synthetic evolutionary origin of a proofreading reverse transcriptase. Science 352, 1590–3 (2016).

40. Houlihan, G. et al. Discovery and evolution of RNA and XNA reverse transcriptase function and fidelity. Nat. Chem. 12, 683–690 (2020).

41. Larsen, A. C. et al. A general strategy for expanding polymerase function by droplet microfluidics. Nat. Commun. 7, 11235 (2016).

42. Czernecki, D., Nourisson, A., Legrand, P. & Delarue, M. Reclassification of family A DNA polymerases reveals novel functional subfamilies and distinctive structural features. Nucleic Acids Res. 51, 4488–4507 (2023).

43. Randrianjatovo-Gbalou, I. et al. Enzymatic synthesis of random sequences of RNA and RNA analogues by DNA polymerase theta mutants for the generation of aptamer libraries. Nucleic Acids Res. 46, 6271–6284 (2018).

44. Ralec, C., Henry, E., Lemor, M., Killelea, T. & Henneke, G. Calcium-driven DNA synthesis by a high-fidelity DNA polymerase. Nucleic Acids Res. 45, 12425–12440 (2017).

45. Dalla Pozza, M., et al. Enzymatic synthesis of glyco-DNA equipped with oligosaccharides and charged monosaccharides. Preprint at 10.26434/chemrxiv.15001602/v1 (2026).

46. Yik, E. J., Maola, V. A. & Chaput, J. C. Engineering TNA polymerases through iterative cycles of directed evolution. in Methods in Enzymology vol. 691 29–59 (Academic Press, 2023).

47. Sciambi, A. & Abate, A. R. Accurate microfluidic sorting of droplets at 30 kHz. Lab Chip 15, 47–51 (2015).

48. Vallejo, D., Nikoomanzar, A., Paegel, B. M. & Chaput, J. C. Fluorescence-Activated Droplet Sorting for Single-Cell Directed Evolution. ACS Synth. Biol. 8, 1430–1440 (2019).

49. Flamme, M. et al. Selection of Ruthenium Polypyridyl Complex-Modified Aptamers for Photodynamic Therapy against Streptococcus Pneumonia. J. Am. Chem. Soc. 147, 43612–43628 (2025).

50. Crameri, A., Raillard, S.-A., Bermudez, E. & Stemmer, W. P. C. DNA shuffling of a family of genes from diverse species accelerates directed evolution. Nature 391, 288–291 (1998).

51. Morcos, F. et al. Direct-coupling analysis of residue coevolution captures native contacts across many protein families. Proc. Natl. Acad. Sci. U. S. A. 108, E1293–301 (2011).

52. Zhou, Z. & Hu, G. Applications of graph theory in studying protein structure, dynamics, and interactions. J. Math. Chem. 62, 2562–2580 (2024).

53. D’Costa, S., Hinds, E. C., Freschlin, C. R., Song, H. & Romero, P. A. Inferring protein fitness landscapes from laboratory evolution experiments. PLoS Comput. Biol. 19, e1010956 (2023).

54. Lin, Z. et al. Evolutionary-scale prediction of atomic-level protein structure with a language model. Science 379, 1123–1130 (2023).

55. Ong, J. L., Loakes, D., Jaroslawski, S., Too, K. & Holliger, P. Directed Evolution of DNA Polymerase, RNA Polymerase and Reverse Transcriptase Activity in a Single Polypeptide. J. Mol. Biol. 361, 537–550 (2006).

56. Maio, G. E. et al. Systematic Optimization and Modification of a DNA Aptamer with 2’-O-Methyl RNA Analogues. ChemistrySelect 2, 2335–2340 (2017).

57. Kratschmer, C. & Levy, M. Effect of Chemical Modifications on Aptamer Stability in Serum. Nucleic Acid Ther. 27, 335–344 (2017).

58. Krömer, M., Poštová Slavětínská, L. & Hocek, M. Glyco-DNA: Enzymatic Synthesis of Base-Modified and Hypermodified DNA Displaying up to Four Different Monosaccharide Units in the Major Groove. Chemistry 30, e202402318 (2024).

59. Stemmer, W. P. C. Rapid evolution of a protein in vitro by DNA shuffling. Nature 370, 389–391 (1994).

60. Maola, V. A. et al. Directed evolution of a highly efficient TNA polymerase achieved by homologous recombination. Nat. Catal. 7, 1173–1185 (2024).

61. Tokuriki, N., Stricher, F., Serrano, L. & Tawfik, D. S. How Protein Stability and New Functions Trade Off. PLoS Comput. Biol. 4, e1000002 (2008).

62. Bloom, J. D., Labthavikul, S. T., Otey, C. R. & Arnold, F. H. Protein stability promotes evolvability. Proc. Natl. Acad. Sci. U. S. A. 103, 5869–5874 (2006).

63. Krauth, W. Statistical Mechanics: Algorithms and Computations. (Oxford University Press Oxford, 2006). doi:10.1093/oso/9780198515357.001.0001.

64. Alvarez, S. et al. In vivo functional phenotypes from a computational epistatic model of evolution. Proc. Natl. Acad. Sci. U. S. A. 121, e2308895121 (2024).

65. Parmentier, G., Schmitt, G., Dolle, F. & Luu, B. A convergent synthesis of 2ʹ-o-methyl uridine. Tetrahedron 50, 5361–5368 (1994).

66. Barandun, L. J. et al. Replacement of Water Molecules in a Phosphate Binding Site by Furanoside-Appended *lin* -Benzoguanine Ligands of tRNA-Guanine Transglycosylase (TGT). Chemistry 21, 126–135 (2015).

67. Cooke, J. W. B., Bright, R., Coleman, M. J. & Jenkins, K. P. Process Research and Development of a Dihydropyrimidine Dehydrogenase Inactivator: Large-Scale Preparation of Eniluracil Using a Sonogashira Coupling. Org. Process Res. Dev. 5, 383–386 (2001).

68. Wang, Q., Ma, X., Chen, Y., Jiang, C. & Xu, Y. Electrochemical Synthesis of 5-Selenouracil Derivatives by Selenylation of Uracils. European J. Org. Chem. 2020, 4384–4388 (2020).

69. Vallejo, D., Nikoomanzar, A. & Chaput, J. C. Directed evolution of custom polymerases using droplet microfluidics. Methods Enzymol. 644, 227–253 (2020).

70. Espada, R., Zarevski, N., Dramé-Maigné, A. & Rondelez, Y. Accurate gene consensus at low nanopore coverage. Gigascience 11, 1–8 (2022).

71. Zurek, P. J., Knyphausen, P., Neufeld, K., Pushpanath, A. & Hollfelder, F. UMI-linked consensus sequencing enables phylogenetic analysis of directed evolution. Nat. Commun. 11, 6023 (2020).

72. Long, Y. et al. LevSeq: Rapid Generation of Sequence-Function Data for Directed Evolution and Machine Learning. ACS Synth. Biol. 14, 230–238 (2025).

73. Karst, S. M. et al. High-accuracy long-read amplicon sequences using unique molecular identifiers with Nanopore or PacBio sequencing. Nat. Methods 18, 165–169 (2021).

74. Li, H. Minimap2: pairwise alignment for nucleotide sequences. Bioinformatics 34, 3094–3100 (2018).

75. Fu, L., Niu, B., Zhu, Z., Wu, S. & Li, W. CD-HIT: accelerated for clustering the next-generation sequencing data. Bioinformatics 28, 3150–3152 (2012).

76. Garrison, E. & Marth, G. Haplotype-based variant detection from short-read sequencing. ArXiv http://arxiv.org/abs/1207.3907 (2012).

77. Sievers, F. et al. Fast, scalable generation of high-quality protein multiple sequence alignments using Clustal Omega. Mol. Syst. Biol. 7, 539 (2011).

78. Martin, D. P., Williamson, C. & Posada, D. RDP2: recombination detection and analysis from sequence alignments. Bioinformatics 21, 260–262 (2005).

79. Chen, J. Z. et al. Understanding epistatic networks in the B1 β-lactamases through coevolutionary statistical modeling and deep mutational scanning. Nat. Commun. 15, 8441 (2024).

80. Patsch, D. et al. Enriching productive mutational paths accelerates enzyme evolution. Nat. Chem. Biol. 20, 1662–1669 (2024).

