## Supplementary Data for "Polymerase Evolution Enables Access to Glycosylated Xeno-Nucleic Acids with Expanded Chemical Functionality"

#### Supplementary Information

Eloi Vincent[1], Victoria Maola[5], Clément Lopez[1], Marc Borie-Guichot[2#], Maria Dalla Pozza[3#], Mohammad Hajjar[5], Rémi Sieskind[1], Soizick Lucas-Staat[1], Rafael Navaza[4], Laurence A. Mulard[2], Marcel Hollenstein[3], John C. Chaput[5-8*], Marc Delarue[1*]

#### Affiliations

1. Institut Pasteur, Université Paris Cité, CNRS UMR 3528, Unit of Architecture and Dynamics of Biological Macromolecules, 25 rue du Docteur Roux, 75015 Paris, France
2. Institut Pasteur, Université Paris Cité, CNRS UMR 3523, Unit of Chemistry of Biomolecules, 28 rue du Docteur Roux, 75724 Paris Cedex 15, France
3. Institut Pasteur, Université Paris Cité, CNRS UMR 3523, Unit of Bioorganic Chemistry of Nucleic Acids, 28 rue du Docteur Roux, 75724 Paris Cedex 15, France.
4. Institut Pasteur, Université Paris Cité, CNRS UMR 3528, Plate-forme de Cristallographie-C2RT, 28 rue du Docteur Roux, 75724 Paris Cedex 15, France
5. Department of Pharmaceutical Sciences, University of California, Irvine, CA, USA
6. Department of Chemistry, University of California, Irvine, CA, USA
7. Department of Molecular Biology and Biochemistry, University of California, Irvine, CA, USA
8. Department of Chemical and Biomolecular Engineering, University of California, Irvine, CA, USA

### These authors contributed equally.

### Table of contents

#### Supplementary data

##### Synthesis of 5-glycosyl-2’-*O*-methyl-NTPs


1. General information
2. General procedures
3. Synthesis & characterization of 5-glycosyl-2’-*O*-methyl-nucleotides
4. NMR spectra ($^{1}H$, $^{13}C$, $^{31}P$) of all compounds

**Protein sequences**

**Nanopore sequencing data processing**


#### Supplementary Figures


**Supplementary Figure 1 | Synthesis of modified nucleotides.**

**Supplementary Figure 2 | Basal activity for glycosylated nucleotides incorporation by various family B DNA polymerases from the literature.**

**Supplementary Figure 3 | Basal activity for 5-glucosyl-2'-*O*-methyl-UTP incorporation by various family A DNA polymerases from the literature.**

**Supplementary Figure 4 | Fidelity assessment of C28 and G2 polymerases by misincorporation assay.**


#### Supplementary Tables


**Supplementary Table 1 | Throughput of droplet-based directed evolution.**

**Supplementary Table 2 | Primers and templates for all polymerase studies, mutagenesis and directed evolution.**

**Supplementary Table 3 | Summary of the high-throughput sequencing (MinION Oxford Nanopore Technology) experiments performed throughout the evolution campaign.**

### Supplementary data

#### Synthesis of 5-glycosyl-2'-*O*-methyl-NTPs

##### 1. General information

All reagents and solvents were purchased from commercial suppliers and used without further purification unless otherwise stated. Reaction progress was monitored by analytical thin-layer chromatography (TLC) on glass plates pre-coated with silica gel 60 F254 (Merck KGaA). TLC plates were visualized under UV light at 254 nm and/or by staining with potassium permanganate followed by heating. Crude reaction mixtures were purified by pre-packed flash column chromatography on silica gel (Interchim) using a Reveleris™ X2-UV system (BUCHI). Solvents were removed under reduced pressure at temperatures below 40 °C using a rotary evaporator, and the resulting residues were further dried under high vacuum.

**HPLC Purification**

Triphosphorylated compounds were purified by anion-exchange high-performance liquid chromatography (HPLC) on an ÄKTA Pure system DNAPacTM PA-100 BioLCTM (Thermo Scientific), 22 x 250mm, Flow: 10mL/min; gradient 0 % B for 5min then 0 to 100 % B in 25min and 100 % B for 5min (A: 10mM TEAB; B: 1M TEAB) at rt. For further purification, when needed, a semi-preparative reverse-phase (RP) column (Phenomenex Luna 5 µm C18 100 Å) was used (water (20 mM of TEAA)/ACN, 100:0 to 90:10). Saccharide and glyco-nucleotides were purified on an Agilent 1260 Infinity II system equipped with a preparative RP column (Kromasil 100-5-C18).

* The gradient is compound-dependent, with a maximum of 10% solvent B.

**Mass Spectrometry**

High-resolution mass spectrometry (HRMS) was performed on a Thermofisher Scientific Q Exactive mass spectrometer with electrospray ionization (H-ESI II probe). MALDI-TOF analyses of triphosphates were performed on a Bruker UltrafleXtreme instrument using 9-aminoacridine as the matrix and operating in linear negative mode.

**Nuclear Magnetic Resonance (NMR) Spectroscopy**

$^{1}$H, $^{13}$C and $^{31}$P NMR spectra were recorded at 298 K on a Bruker Avance III (400 MHz) , a Bruker Avance III HD (500 MHz), or a Bruker Avance Neo (600 et 800 MHz) spectrometers. Chemical shifts (δ) are reported in parts per million (ppm) relative to residual solvent signals: $^{1}$H δ = 7.26 ($CDCl_3$), 4.76 ($D_2O$), 3.31($CD_3OD$), 2.50 (DMSO-d6) ppm; ($^{13}$C δ = 77.16 ($CDCl_3$), 49.00 ($CD_3OD$), 39.52 (DMSO-d6). Data are reported as follows: chemical shift, multiplicity (s = singlet, br s = broad singlet, d = doublet, dd = doublet of doublets, dt = doublet of triplets, t = triplet, q = quartet, m = multiplet), coupling constants J (Hz), and integration. Carbon multiplicities were assigned using DEPT experiments. When necessary, $^{1}$H and $^{13}$C resonances were assigned using 2D NMR techniques such as COSY, HSQC and HMBC.

**PCR and Primer Extension reactions**

PCR and primer extension experiments were performed on a SimpliAmp™ thermal cycler (Thermo Fisher Scientific).

**Polyacrylamide Gel Electrophoresis (PAGE)**

Acrylamide/bisacrylamide (29:1, 40%) solutions were obtained from Fisher Scientific. PAGE analyses were performed and visualized by fluorescence imaging using a Typhoon Trio scanner (GE Healthcare). The gels were run using 1x TBE buffer (89 mM Tris-borate, 2 mM EDTA) as buffer, 1.25x formamide loading buffer (70% formamide, 50 mM EDTA, 0.1% bromophenol, and 0.1% xylenecyanol). After loading, the gel was run at rt for 4h at 200S-6 250 V. Gel electrophoresis was run with a trisborate–EDTA (TBE) buffer at a 1× concentration (pH 8, 7 M urea) at 40W for 4 h or at 4W for 12 h and visualized afterwards by UV light (255 and 350 nm).

**UV-Vis Spectroscopy**

UV–Vis analyses were performed using a Cary 3500 Compact Peltier spectrophotometer (Agilent Technologies) with 1 mL quartz cuvettes for quantification, and 70 µL quartz cuvettes for the melting temperatures curves. Melting temperatures have been calculated by triplicate experiments, calculating each derivative value of the different curves.

#### 2. General procedures

**Protocol A**: Dowex ($H^+$) (1:1 w/w with the starting material) was added to a mixture of saccharide (1.0 equiv.) and bromoethanol (18.0 equiv.) in a microwave vial reactor. The reaction mixture was stirred for 15 min at 120 °C (20 W). After completion, the resulting brown/dark liquid was diluted with EtOAc (10 mL) and filtered through a cotton pad. The filtrate was concentrated under reduced pressure. The crude product was purified by automated flash chromatography on prepacked silica gel columns (EtOAc/MeOH 98:2 to 90:10), affording a mixture of α/β-anomers.

**Protocol B**: Sodium azide (2 equiv.) was added to a solution of bromoethyl saccharide (1 equiv.) in anhydrous DMF (5 $mL.mmol^{-1}$). The reaction mixture was stirred overnight at 80 °C. After completion, the resulting brown liquid was diluted with EtOAc (10 mL) and then concentrated under reduced pressure. The crude product was purified by RP-HPLC (100/0 to 98-90/2-10; water/ACN), to separate α-anomers.

**Protocol C**: Acetic acid (4 equiv. / *O*-acetyl protected alcohol) was added to a solution of oligosaccharide (1 equiv.) in dry pyridine (8 mL.mmol-1) at 0 °C. A catalytic amount of DMAP 4-dimethylaminopyridine was then added, and the reaction mixture was stirred overnight at room temperature. At completion, the mixture was diluted with EtOAc (50 mL) and washed successively with 1 M HCl (20 mL) and saturated aqueous $NaHCO_3$ (20 mL). The organic phase was dried over anhydrous sodium sulfate, filtered, and concentrated with toluene (3 × 20 mL) under reduced pressure to afford a white solid. The crude product obtained as a mixture of α/β-anomers was used as such in the next step.

$BF_3 \cdot Et_2O$ (6 equiv.) was added dropwise to a solution of bromoethanol (3 equiv.), molecular sieves (1 equiv. w/w) and crude (1 equiv.) in DCE (10 $mL.mmol^{-1}$), previously stirred for 15 min at 0 °C in a microwave vial reactor. The reaction mixture was then stirred for 10 min at 80 °C under microwave irradiation. The resulting yellow liquid was diluted with EtOAc (20 mL) and filtered through a cotton pad. The filtrate was then concentrated under reduced pressure. The crude product was passed through a pre-packed silica gel column eluting with a cyclohexane/EtOAc gradient (70:30 to 0:100), automated flash chromatography) to afford a mixture of partially deacetylated α/β-anomers.

This mixture was then taken up with aqueous $NH_4OH_{aq}$ (10 $mL.mmol^{-1}$) in MeOH (20 $mL.mmol^{-1}$) and stirred overnight at room temperature. At completion, the reaction mixture was concentrated and co-evaporated first with water (50 mL) and then with MeOH (20 mL). The crude product was isolated as a mixture of α/β-anomers and used as such in the next step.

**Protocol D**: CuI (2 equiv., 8 $mg.mL^{-1}$) in degassed DMF/water (1:1) was added to a mixture of azidoethyl saccharide (2–4 equiv.) and 5-ethynyl-2'-*O*-methyl-uridine triphosphate **4** (1 equiv.) in degassed water (0.3 $mL.mmol^{-1}$) under an inert atmosphere in an 2 mL eppendorf tube. Then, N,N-diisopropylethylamine (6 equiv.) was added, and the reaction mixture was stirred at 25 °C for 4 h. At completion, the reaction mixture was poured into cold acetone (10 mL), and the resulting precipitate was collected by centrifugation, then lyophilized. The precipitate was taken in an $EDTA_{aq}$ solution and was purified by RP-HPLC (20 mM TEAA in water)/ACN, 100 to 97-93:3-7), affording the desired triphosphate as a white solid after multiple lyophilizations.

#### 3. Synthesis & characterization of 5-glycosyl-2'-*O*-methyl-nucleotides

***Compound 9*[1,2]**

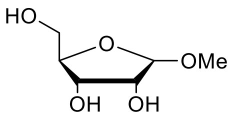


Chemical Formula: $C_6H_{12}O_5$
Exact Mass: 164,07
Molecular Weight: 164,16 g.mol$^{-1}$

Dowex ($H^+$) (1.00 g) was added to a solution of D-ribose (1.00 g; 1.0 equiv.) in MeOH (5.38 mL; 20.0 equiv.) in a microwave tube reactor vial. The reaction mixture was stirred at 100 °C (20 W) for 40 minutes. The conversion of the starting material was monitored by TLC (eluent: EtOAc/MeOH 90:10), showing the formation of a less polar compound. At completion, the resulting orange solution was filtered and washed with MeOH (100 mL). An aqueous ammonia solution (30%) (10 mL) was then added to the filtrate, which was subsequently concentrated under reduced pressure. The crude product was purified by flash column chromatography on silica gel (EtOAc/MeOH; 98:2 to 92:8) to afford a mixture of α/β anomers (1.05 g, 98%).

**$^1$H NMR (400 MHz, MeOD)** δ: 4.77 (s, 1H), 4.61 (d, 1H, *J* = 3.7 Hz), 4.04 (dd, 1H, *J* = 6.6, 4.7 Hz), 3.94 (td, 1H, *J* = 6.6, 3.5 Hz), 3.88 (brd, 1H, *J* = 4.7 Hz), 3.83-3.79 (m, 1H), 3.78-3.63 (m, 4H), 3.58-3.41 (m, 2H), 3.41 (s, 3H), 3.35 (s, 3H).

**$^{13}$C NMR (100 MHz, MeOD)** δ: 109.8, 103.3, 84.8, 76.1, 72.6, 72.3, 70.5, 68.0, 64.9, 64.7, 56.0, 55.4.

***Compound 10*[1,2]**

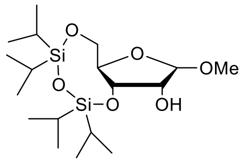


Chemical Formula: $C_{18}H_{38}O_6Si_2$
Exact Mass: 406,22
Molecular Weight: 406,66 g.mol$^{-1}$

1,3-Dichloro-1,1,3,3-tetraisopropyldisiloxane (11.43 g, 1.0 equiv.) was added dropwise over 30 minutes to a solution of **9** (5.80 g, 1.0 equiv.) in pyridine cooled to −15 °C. After an additional 20 minutes, the cooling bath was removed, and the reaction mixture was stirred at room temperature for 3 hours. The conversion of the starting material was monitored by TLC (eluent: EtOAc/MeOH 90:10), showing the formation of a less polar compound. EtOAc (150 mL) was then added, and the organic phase was washed with water (2 × 80 mL). The organic layer was dried, filtered, and concentrated under reduced pressure. The crude product was purified by flash column chromatography on silica gel (cyclohexane/EtOAc gradient from 98:2 to 92:8) to afford a mixture of α/β anomers (13.93 g, 97%).

**$^1$H NMR (400 MHz, DMSO-d6)** δ: 5.00 (d, 1H, *J* = 3.9 Hz), 4.81 (d, 1H, *J* = 6.8 Hz), 4.60 (s, 1H), 4.36 (d, 1H, *J* = 7.3 Hz), 4.31-4.27 (m, 1H), 4.26 (dd, 1H, *J* = 7.4, 4.4 Hz), 4.02 (ddd, 1H, *J* = 9.3, 4.5,2.3 Hz), 3.88 (dd, 1H, *J* = 11.6, 3.0 Hz), 3.85-3.71 (m, 2H), 3.60 (dd, 1H, *J* = 10.9, 4.6 Hz), 3.49 (dd, 1H, *J* = 10.5, 9.7 Hz), 3.34 (s, 3H), 3.18 (s, 3H), 1.09-0.92 (m, 28H).

**$^{13}$C NMR (100 MHz, DMSO-d6)** δ: 107.1, 101.4, 80.1, 74.6, 73.2, 73.0, 71.0, 69.6, 63.8, 63.0, 55.7, 53.9, 17.3, 11.7.

***Compound 2[1,2]***

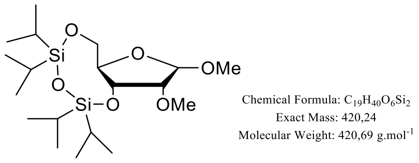


To a solution of **10** (13.80 g, 1.0 equiv.) in MeI (60 mL) under an argon atmosphere, NaH (4.75 g, 6.0 equiv.) was added over 30 minutes. The reaction mixture was then stirred at room temperature for 2 hours. EtOAc (150 mL) was added, and the resulting organic phase was washed with water (2 × 80 mL). The organic layer was dried, filtered, and concentrated under reduced pressure. The crude product was purified by flash column chromatography on silica gel (cyclohexane/EtOAc 98:2 to 95:5) to afford the desired compound as a mixture of a mixture of α/β anomers (5.21 g, 36%).

**$^1$H NMR (400 MHz, $CDCl_3$)** δ: 4.78 (s, 1H), 4.61 (d, 1H, *J* = 7.4 Hz), 4.55-4.53 (m, 1H), 4.50 (dd, 1H, *J* = 7.9, 4.3 Hz), 4.05-4.00 (m, 3H), 3.93-3.87 (m, 1H), 3.77-3.67 (m, 2H), 3.62 (brd, 1H, *J* = 4.3 Hz), 3.59 (s, 3H), 3.54 (s, 3H), 3.49 (s, 3H), 3.43 (s, 3H), 3.01 (dd, 1H, *J* = 7.4, 2.7 Hz), 1.15-1.01 (m, 28H).

**$^{13}$C NMR (100 MHz, $CDCl_3$)** δ: 105.7, 101.1, 84.5, 81.0, 80.2, 73.7, 71.7, 71.2, 63.9, 63.8, 58.7, 56.7, 54.7, 17.5, 12.6.

***Compound 11[1,2]***

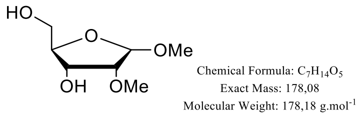


To a solution of **2** (5.06 g, 1.0 equiv.) in THF (350 mL) cooled to 0 °C, TBAF·3$H_2$O (11.95 g, 3.2 equiv.) was added over 20 min. The reaction mixture was stirred for 2 h while allowing the temperature to gradually rise to room temperature. The crude product was purified by flash column chromatography on silica gel (cyclohexane/EtOAc 90:10 to 0:100) to afford a mixture of α/β anomers (1.77 g, 83%).The mixture was then concentrated under reduced pressure.

**$^1$H NMR (400 MHz, MeOD)** δ: 4.86 (d, 1H, , *J* = 0.9 Hz), 4.70 (d, 1H, *J* = 4.2 Hz), 4.10 (dd, 1H, *J* = 6.5, 5.0 Hz), 3.96-3.93 (m, 1H), 3.90 (dt, 1H, *J* = 6.5, 3.6Hz), 3.76-3.64 (m, 4H), 3.62 (t, 1H, *J* = 5.3 Hz), 3.56-3.50 (m, 2H), 3.48 (s, 6H), 3.42 (s, 3H), 3.37 (s, 3H), 3.21 (ddd, 1H, *J* = 4.2, 3.3, 0.8 Hz).

**$^{13}$C NMR (100 MHz, MeOD)** δ: 107.0, 100.8, 85.6, 85.2, 81.7, 72.2, 70.2, 67.7, 64.8, 64.7, 59.0, 58.8, 56.1, 55.5.

***Compound 12[2]***

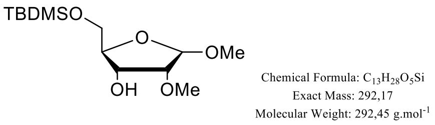


To a solution of **11** (1.70 g, 1.0 equiv.), Triethylamine (3.3 mL, 2.5 equiv.), and 4-Dimethylaminopyridine (cat.) in DMF (350 mL) cooled to 0 °C, *tert*-Butyldimethylsilyl chloride (1.44 gr, 1.0 equiv.) was added dropwise over 1 hour. The reaction mixture was then stirred overnight at room temperature. The mixture was concentrated under reduced pressure. The crude product was purified by flash column chromatography on silica gel (cyclohexane/EtOAc 99:1 to 80:20, then 0:100) to afford a mixture of α/β anomers (1.55 g, 55%).

**$^1$H NMR (400 MHz, $CDCl_3$)** δ: 4.91 (d, 1H, *J* = 1.3 Hz), 4.62 (d, 1H, *J* = 7.0 Hz), 4.26-4.20 (m, 1H), 4.18-4.15 (m, 1H), 3.96-3.92 (m, 1H), 3.83 (ddd, 1H, *J* = 8.7, 5.3, 2.9 Hz), 3.80-3.70 (m, 1H), 3.78 (dd, 1H, *J* = 10.9, 4.4 Hz), 3.72 (dd, 1H, *J* = 10.9, 5.1 Hz), 3.67-3.63 (m, 2H), 3.55 (s, 3H), 3.54 (s, 3H), 3.53 (s, 3H), 3.39 (s, 3H), 3.07 (ddd, 1H, *J* = 7.2, 2.6, 0.7 Hz), 2.53 (d, 1H, *J* = 8.1 Hz), 2.52-2.50 (m, 1H), 0.96-0.88 (m, 9H), 0.16-0.08 (m, 6H).

**$^{13}$C NMR (100 MHz, $CDCl_3$)** δ: 105.2, 100.8, 84.6, 84.1, 79.5, 71.1, 69.1, 68.5, 64.2, 63.5, 58.4, 58.4, 56.7, 55.2, 25.9, 25.7, 18.4, 18.1.

***Compound 3***

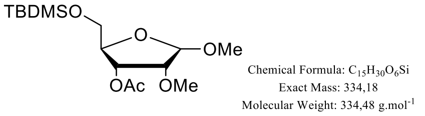


To a solution of **12** (1.70 g, 1.0 equiv.), Triethylamine (3.3 mL, 2.5 equiv.), and DMAP (cat.) in DMF (350 mL) cooled to 0 °C, *tert*-butyldimethylsilyl chloride (1.44 gr, 1.0 equiv.) was added dropwise over 1 hour. The reaction mixture was then stirred overnight at room temperature. The crude product was purified by flash column chromatography on silica gel (cyclohexane/EtOAc 99:1 to 80:20, then 0:100) to afford a mixture of α/β anomers (1.55 g, 55%). The mixture was concentrated under reduced pressure.

**$^1$H NMR (400 MHz, $CDCl_3$)** δ: 5.62-5.58 (m, 1H), 5.2 ($t_{app}$, *J* = 5.1 Hz), 4.92 (d, 1H, *J* = 2.3 Hz), 4.49 (d, 1H, *J* = 7.6 Hz), 4.16 ($q_{app}$, 1H, *J* = 5.0 Hz), 3.88 (dd, 1H, *J* = 5.0, 2.2 Hz), 3.88-3.82 (m, 1H), 3.75 (dd, 1H, *J* = 10.9, 4.7 Hz), 3.70 (dd, 1H, *J* = 11.2, 5.4 Hz), 3.69 (dd, 1H, *J* = 10.9, 5.1 Hz), 3.60 (dd, 1H, *J* = 11.2, 10.6 Hz), 3.54 (s, 3H), 3.44 (s, 3H), 3.43 (s, 3H), 3.41 (s, 3H), 3.12 (dd, 1H, *J* = 7.7, 3.0 Hz), 2.13 (s, 6H), 0.96-0.88 (m, 9H), 0.16-0.08 (m, 6H).

**$^{13}$C NMR (100 MHz, $CDCl_3$)** δ: 170.3 (C=O), 106.3, 101.1, 82.8, 81.7, 80.2, 73.3, 70.0, 68.3, 64.1, 60.9, 58.8, 58.8, 56.8, 55.5, 25.8, 25.8, 20.8, 20.8.

**HRMS (ESI)**: *m/z* calcd. For $C_{15}H_{34}NO_6Si$ $[M+NH_4]^+$: 352.2150; found: 352.2141.

***Compound 13[3]***

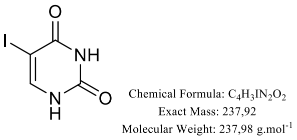


*N*-Iodosuccinimide (8.74 g, 1.6 equiv.) was added to a solution of uracil (2.70 g, 1.0 equiv.) in anhydrous DMF (60 mL) under an argon atmosphere in a microwave reactor vial. The reaction mixture was stirred at 130 °C (15 W) for 30 min. The mixture was then concentrated under reduced pressure, and the resulting solid was filtered and washed with EtOAc (800 mL) to afford a white solid. The product required no further purification (4.67 g, 81%).

**$^1$H NMR (400 MHz, DMSO-d6)** δ: 11.42 (s, 1H), 11.17 (s, 1H), 7.89 (s, 1H).

Spectral data are in accordance with literature.

***Compound 14[4]***

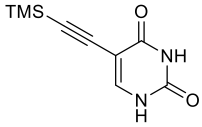


Chemical Formula: $C_9H_{12}N_2O_2Si$
Exact Mass: 208,07
Molecular Weight: 208,29 g.mol$^{-1}$

Compound **13** (4.65 g, 1.0 equiv.) was suspended in anhydrous THF (90 mL) under an argon atmosphere. $PdCl_2$ (0.34 g, 0.1 equiv.), $PPh_3$ (0.75 g, 0.15 equiv.), and CuI (0.75 g, 0.2 equiv.) were added, and the reaction mixture was degassed under argon. Then, Triethylamine (5.44 mL, 2.0 equiv.) and Trimethylsilylacetylene (6.80 g, 3.5 equiv.) were added, and the mixture was stirred at 40 °C in the dark for 18 h. THF was added, and the suspension was filtered through Celite pad. The filtrate was concentrated under reduced pressure, and the residue was taken up in DCM/MeOH (97:3). The suspension was filtered to afford a white solid (2.69 g, 66%).

**$^1$H NMR (400 MHz, DMSO-d6)** δ: 11.39-11.26 (m, 2H), 7.81 (s, 1H), 0.19 (s, 9H).

Spectral data are in accordance with literature.

***Compound 15[4]***

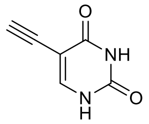


Chemical Formula: $C_6H_4N_2O_2$
Exact Mass: 136,03
Molecular Weight: 136,11 g.mol$^{-1}$

Compound **14** (1.0 equiv.) was suspended in MeOH (30 mL), and a solution of $NaOH_{aq}$ (1 M; 72.0 mL) was added dropwise. The reaction mixture was stirred at room temperature for 2 hours. The mixture was then concentrated under reduced pressure, and the residue was taken up in DCM/MeOH (95:5). The suspension was filtered and washed successively with DCM/MeOH (95:5) and DCM/MeOH (90:10) to afford a white solid (1.34 g, 88%).

**$^1$H NMR (400 MHz, DMSO-d6)** δ: 11.47-11.22 (m, 2H), 7.83 (s, 1H), 4.05 (s, 1H).

Spectral data are in accordance with literature.

***Compound 16***

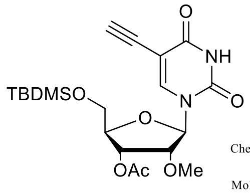


Chemical Formula: $C_{20}H_{30}N_2O_7Si$
Exact Mass: 438,18
Molecular Weight: 438,55 g.mol$^{-1}$

N,O-bis(trimethylsilyl)acetamide (2.5 mL, 4.5 equiv.) was added to a solution of **15** (0.626g, 2.2 equiv.) in ACN (20 mL) under an argon atmosphere. The reaction mixture was stirred at room temperature for 30 min. Then, a solution of **3** (0.700g, 1 equiv.) and Trimethylsilyl trifluoromethanesulfonate (0.70 mL, 2 equiv.) in MeCN (30 mL) was added dropwise, and the reaction mixture was stirred at 80 °C for 24 hours. The reaction mixture was quenched with an aqueous solution of $NaHCO_3$ (80 mL), and the organic phase was extracted with DCM (3 × 80 mL). The combined organic layers were dried, filtered, and concentrated under reduced pressure. The crude product was purified by flash column chromatography on silica gel (cyclohexane/EtOAc 90:10 to 50:50,) to afford a colorless oil (0.52 g, 57%).

**$^1$H NMR (400 MHz, MeOD)** δ: 8.12 (s, 1H), 6.00 (d, 1H, *J* = 5.5 Hz), 5.25 (dd, 1H, *J* = 5.0, 3.9 Hz), 4.24 ($td_{app}$, 1H, *J* = 3.7, 1.9 Hz), 4.04 ($t_{app}$, 1H, *J* = 5.2 Hz), 3.99 (dd, 1H, *J* = 11.8, 2.0 Hz), 3.87 (dd, 1H, *J* = 11.9, 2.0 Hz), 3.63 (s, 1H), 3.40 (s, 3H), 2.13 (s, 3H), 0.98 (s, 9H), 0.19 (s, 3H), 0.19 (s, 3H).

**$^{13}$C NMR (100 MHz, MeOD)** δ: 171.6, 163.8, 151.1, 144.4, 100.5, 88.0, 84.5, 84.0, 83.8, 75.9, 71.7, 63.7, 59.4, 26.6, 20.6, 19.3.

***<u>Compound 17</u>***

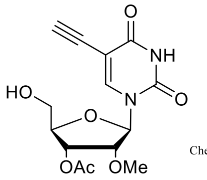


Chemical Formula: $C_{14}H_{16}N_2O_7$
Exact Mass: 324,10
Molecular Weight: 324,29 g.mol$^{-1}$

Tetrabutylammonium fluoride (1 M/THF) (2.50 mL) was added dropwise to a solution of **16** (0.500 g, 1.0 equiv.) in anhydrous THF under an argon atmosphere at 0 °C. The reaction mixture was then allowed to warm to room temperature and stirred for 3 hours. The mixture was quenched with an aqueous solution of $NaHCO_3$ (50 mL), and the organic phase was extracted with EtOAc (3 × 50 mL). The combined organic layers were dried, filtered, and concentrated under reduced pressure. The crude product was purified by flash column chromatography on silica gel (DCM/EtOAc 90:10 to 30:70) to afford a white solid (0.26 g, 71%).

**$^1$H NMR (400 MHz, MeOD)** δ: 8.47 (s, 1H), 5.98 (d, 1H, *J* = 5.1 Hz), 5.26 (t, 1H, *J* = 4.7 Hz), 4.19 ($td_{app}$, 1H, *J* = 4.6, 2.4 Hz), 4.14 ($t_{app}$, 1H, *J* = 5.1 Hz), 3.87 (dd, 1H, *J* = 12.3, 2.5 Hz), 3.75 (dd, 1H, *J* = 12.3, 2.4 Hz), 3.57 (s, 1H), 3.41 (s, 3H), 2.15 (s, 3H).

**$^{13}$C NMR (100 MHz, MeOD)** δ: 171.8, 164.1, 151.3, 145.8, 100.2, 88.7, 84.7, 83.5, 83.1, 75.8, 72.0, 61.7, 59.3, 20.6.

**HRMS (ESI)**: *m/z* calcd. For $C_{14}H_{15}N_2O_7$ $[M-H]^-$: 323.0885; found: 323.0882.

***<u>Compound 4: 5-ethynyl-2'-O-methyl-UTP</u>***

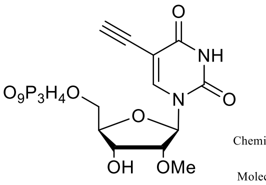


Chemical Formula: $C_{12}H_{17}N_2O_{15}P_3$
Exact Mass: 521,98
Molecular Weight: 522,19 g.mol$^{-1}$

Nucleoside **16** (30.0 mg, 1.0 equiv.) was dried under reduced pressure overnight prior reaction, then dissolved in the mixture (1:2) of dry pyridine (0.20 mL) and dioxane (0.40 mL) under argon. To this solution, 2-chloro-1,3,2-benzodioxaphosphorin-4-one (49.0 mg, 2.6 equiv.) dissolved in dry dioxane (0.15 mL) was added at 0 °C and the reaction mixture was stirred for 4.5 hours at room temperature under argon. Next, the solution of tributylammonium pyrophosphate (100.0 mg, 2.5 equiv.), in the mixture (1:3) of dry tributylamine (0.15 mL) and DMF (0.45 mL) was added dropwise at 0 °C and the reaction mixture was stirred at room temperature for 2 hours under argon. The mixture was then oxidized by the addition of iodine (23.0 mg, 0.8 equiv.) in pyridine (0.50 mL) and $H_2O$ (0.25 mL) at 0 °C. After 20 min of stirring, the excess iodine was quenched with sodium thiosulfate solution (10% w/v in water) and the reaction mixture was concentrated under reduced pressure at 30 °C. The residue was dissolved in $H_2O$ (1 mL)and precipitated by adding it dropwise to a 2% $NaClO_4$ solution in acetone (20 mL) then centrifuged and acetone was decanted. The precipitate was dried under reduced pressure.

The crude product was purified by HPLC using a preparative ion exchange column (Buffer A: 10 mM TEAB, Buffer B: 1M TEAB) to afford a yellow solid (7.8 mg, 16%).

**$^{1}$H-NMR (500 MHz, D$_2$O)** δ: 8.24 (s, 1H, H-6), 6.02 (d, *J* = 4.2 Hz, 1H, H-1’), 4.53 (dd, *J* = 10.4, 5.1 Hz, 1H, H-3’), 4.33 – 4.24 (m, 3H, H-4’-H-5’), 4.10 (dd, *J* = 9.6, 4.9 Hz, 1H, H-2’), 3.62 (s, 1H, ethynyl -CH), 3.53 (s, 3H, H-2’OMe).

**$^{13}$C NMR (125 MHz, D$_2$O)** δ 164.5 (C-4), 150.5 (C-2), 145.1 (C-6), 99.0 (C-5), 87.3 (C-1’), 83.6 (C-Ethynyl), 83.3 (C-5’), 82.6 (C-2’), 67.8 (C-3’), 64.6 (C-4’), 58.2 (C-2’OMe).

**$^{31}$P-NMR (203 MHz, D$_2$O)** δ -10.8 (brs), -11.5 (d, *J* = 19.5 Hz), -22.9 (brs).

**HRMS (ESI)**: *m/z* calcd. For $C_{12}H_{16}N_2O_{15}P_3$ [M-H]$^-$: 520.98; found: 520.9769.

***<u>Compound 18</u>***

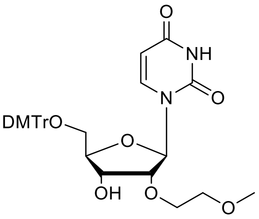


Chemical Formula: $C_{33}H_{36}N_2O_9$
Exact Mass: 604,24
Molecular Weight: 604,66 g.mol$^{-1}$

2'-*O*-(2-Methoxyethyl)-uridine (100 mg, 330 µmol) was dissolved in 0.6 mL of dry pyridine and 4,4’-Dimethoxytrityl chloride (135.0 mg, 1.2 equiv.) was dissolved in 0.65 mL of dry pyridine and added dropwise under stirring at room temperature. After 2.5 hours the solvent was removed, following extraction in DCM/$H_2O$. The combined organic phases were dried over $Na_2SO_4$ and concentrated in vacuo. The final product was obtained after purification in flash column chromatography (DCM/MeOH; 100 to 95:5) as an orange oil (180.0 mg, 90%).

***<u>Compound 19</u>***

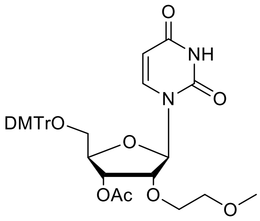


Chemical Formula: $C_{35}H_{38}N_2O_{10}$
Exact Mass: 646,25
Molecular Weight: 646,69 g.mol$^{-1}$

Compound **18** (180 mg, 1.0 equiv.) was dissolved in dry pyridine (1.4 mL) and acetic anhydride (0.28 mL, 10.0 equiv.) was added dropwise at room temperature and the reaction was stirred overnight. Then, the reaction was quenched with MeOH (5.0 mL), stirring 10 min, and removed the solvent under vacuo. Then, the residue was extracted with EtOAc/$H_2O$, the collected organic phases were dried over $Na_2SO_4$ and concentrated in vacuo, to afford a white solid. (150.0 mg, 78%).

***Compound 20***

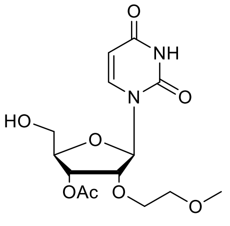

Chemical Formula: $C_{14}H_{20}N_2O_8$
Exact Mass: 344,12
Molecular Weight: 344,32 g.mol$^{-1}$

A solution of Dichloroacetic acid (2%) (3.0 mL) in DCM was added dropwise to **19** (150.0 mg, 1.0 equiv.) and stirred 30 min at room temperature. The residue was dried under vacuo and purified by flash chromatography (EtOAc/hexane; 70:30) to afford a white solid (42.0 mg, 52%).

***Compound 21: 2'-O-(2-Methoxyethyl)-uridine triphosphorylation***

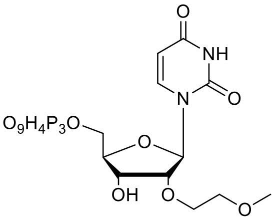

Chemical Formula: $C_{12}H_{21}N_2O_{16}P_3$
Exact Mass: 542,01
Molecular Weight: 542,22 g.mol$^{-1}$

Compound **20** (41.0 mg, 1.0 equiv.) was dried under reduced pressure overnight prior reaction, then dissolved in the mixture of dry pyridine (0.25 mL) and dioxane (0.50 mL) under argon. To this solution, 2-chloro-1,3,2-benzodioxaphosphorin-4-one (44.0 mg, 1.5 equiv.) dissolved in dry dioxane (0.25 mL) was added dropwise at 0 °C and the reaction mixture was stirred for 4.5 hours at room temperature under argon. Next, the solution of tributylammonium pyrophosphate (117.0 mg, 1.8 equiv.), in the mixture (1:3) of dry tributylamine (0.15 mL) and anhydrous DMF (0.45 mL) was added dropwise at 0 °C and the reaction mixture was stirred at room temperature for 1 hour under argon. The mixture was then oxidized by the addition of iodine (55 mg, 1.5 equiv.) in pyridine (0.9 mL) and water (0.3 mL) at 0 °C. After 20 min of stirring, the excess iodine was quenched with sodium thiosulfate solution (10% w/v in water) and the reaction mixture was concentrated under reduced pressure at 30 °C. The residue was dissolved in 2 mL of $H_2O$ and precipitated by adding it dropwise to a 2% $NaClO_4$ solution in acetone (20 mL) then centrifuged and acetone was decanted. The precipitate was dried under reduced pressure. The crude product was purified by HPLC using a preparative ion exchange column (Buffer A: 10 mM TEAB, Buffer B: 1M TEAB) to afford a white solid (3.93 mg, 5%).

**$^1$H NMR (500 MHz, $D_2O$)** δ 8.00 (d, *J* = 8.2 Hz, 1H, H-6), 6.02 (d, *J* = 4.4 Hz, 1H, H-5), 5.97 (dd, *J* = 8.1 Hz, 1H, H-1'), 4.51 (dd, *J* = 9.6, 4.9 Hz, 1H, H-3'), 4.31 – 4.22 (m, 4H, H-4', H-5'), 4.19 (dd, *J* = 4.8, 9.6 Hz, 1H, H-2'), 3.88 – 3.84 (m, 2H, O-$CH_2$), 3.67 – 3.61 (m, 2H, O-$CH_2$-O), 3.35 (s, 3H, O-$CH_3$).

**$^{13}$C NMR (125 MHz, $D_2O$)** δ 166.25 (C-4), 151.64 (C-2), 141.66 (C-6), 102.59 (C-1'), 87.00 (C-5), 83.32 (C-4'), 83.28 (C-4'), 81.52 (C-2'), 71.10 (-O$CH_2$-), 69.44 (-$CH_2$O-), 68.11 (C-3'), 64.54 (OMe).

**$^{31}$P NMR (203 MHz, $D_2O$)** δ -8.8 (brs), -11.3 (d, *J* = 19.8 Hz), -22.4 (brs).

**HRMS (ESI)**: *m/z* calcd. For $C_{12}H_{17}N_2O_{15}P_3$ [M-H]$^-$: 541.00; found: 541.0031.

***Compound 22***

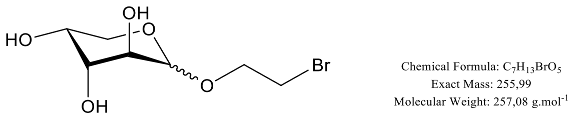


Following the protocol **A**, the compound was obtained as a mixture of α/β anomers (90/10) as a yellow oil (0.43 g, 51%).

**$^1$H NMR (400 MHz, $D_2O$) (α anomer)** δ: 5.06 (d, 1H, *J* = 3.6 Hz, H-1), 4.11-4.03 (m, 3H, H-4, H-5, H-6), 4.01-3.93 (m, 2H, H-3, H-6), 3.90 (dd, 1H, *J* = 10.2, 3.6 Hz, H-2), 3.75-3.63 (m, 3H, H-5, H-7).

**$^{13}$C NMR (100 MHz, $D_2O$) (α anomer)** δ: 98.9 (C-1), 69.0-68.8 (C-3, C-4), 68.3 (C-6), 68.3 (C-2), 63.1 (C-5), 31.4 (C-7).

**HRMS (ESI)**: *m/z* calcd. For $C_8H_{14}BrO_7$ $[M+HCOO]^-$: 300.9928; found: 300.9926.

***Compound 23***

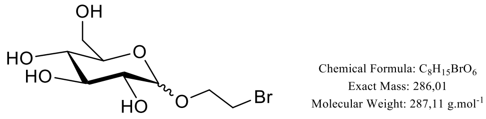


Following the protocol **A**, the compound was obtained as a mixture of α/β anomers (75/25) as a yellow oil (0.52 g, 65%).

**$^1$H NMR (400 MHz, $D_2O$) (α anomer)** δ: 5.05 (d, 1H, *J* = 3.8 Hz, H-1), 4.10 (quint, 1H, *J* = 5.5 Hz, H-7), 4.02-3.75 (m, 2H, H-6), 3.96-3.76 (m, 2H, H-3, H-4), 3.74-3.67 (m, 2H, H-8), 3.63 (dd, 1H, *J* = 9.9, 3.8 Hz, H-2), 3.48 ($t_{app}$, 1H, *J* = 9.4 Hz, H-5).

**$^{13}$C NMR (100 MHz, $D_2O$) (α anomer)** δ: 98.3 (C-1), 73.1 (C-3), 72.1 (C-4), 71.4 (C-2), 69.6 (C-5), 68.2 (C-7), 60.6 (C-6), 31.4 (C-8).

**HRMS (ESI)**: *m/z* calcd. For $C_9H_{16}BrO_8$ $[M+HCOO]^-$: 331.0034; found: 331.0025.

***Compound 24***

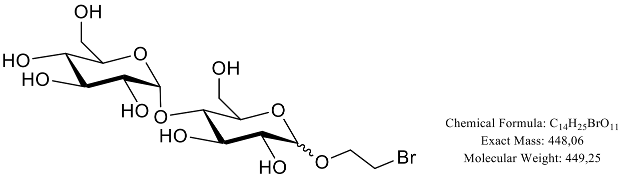


Following the protocol **C**, the compound was obtained as a mixture of α/β anomers (30/70) as a yellow oil (1.64 g, 56%).

**$^1$H NMR (400 MHz, $D_2O$) (α anomer)** δ: 5.45 (d, 1H, *J* = 3.9 Hz, H-1’), 5.05 (d, 1H, *J* = 3.8 Hz, H-1), 4.11-4.02 (m, 3H, H-3, H-3’, H-7), 4.06-3.87 (m, 3H, H-6 or H-6’, H-7), 3.87-3.61 (m, 8H, H-2, H-2’, H-4, H-4’, H-6 or H-6’, H-8), 3.48-3.38 (m, 2H, H-5, H-5’).

**$^{13}$C NMR (100 MHz, $D_2O$) (α anomer)** δ: 98.8 (C-1’), 97.2 (C-1), 76.1-72.5-71.9-71.8 (C-3, C-3’, C-4, C-4’), 70.9-70.3 (C-2, C-2’), 69.7-68.5 (C-5, C-5’), 67.4 (C-7), 59.8-59.8 (C-6, C-6’), 30.5 (C-8).

**HRMS (ESI)**: *m/z* calcd. For $C_{14}H_{25}BrO_{11}$ $[M+NH_4]^+$: 466.0918; found: 466.0915.

***Compound 25***

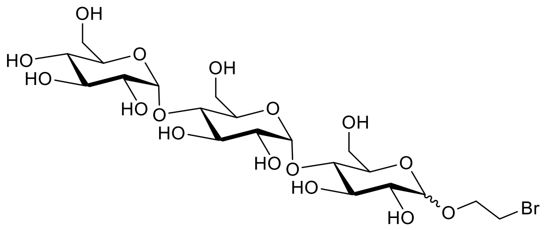

Chemical Formula: $C_{20}H_{35}BrO_{16}$
Exact Mass: 610,1108
Molecular Weight: 611,3880 g.mol$^{-1}$

Following the protocol **C**, the compound was obtained as a mixture of α/β anomers (35/65) as a yellow oil (0.15 g, 90%).

**$^1$H NMR (400 MHz, MeOD) (α anomer)** δ: 5.20-5.10 (m, 2H, H-1’, H-1’’), 4.96-4.75 (m, 1H, H-1), 4.15-3.72 (m, 9H, H-3, H-3’, H-3’’, H-6 or H-6’ or H-6’’, H-7), 3.72-3.38 (m, 13H, H-2, H-2’, H-2’’, H-4, H-4’, H-4’’, H-5, H-5’, H-5’’, H-6 or H-6’ or H-6’’, H-8).

**$^{13}$C NMR (100 MHz, MeOD) (α anomer)** δ: 102.7-102.6 (C-1’, C-1’’), 100.4 (C-1), 81.2-77.7-74.9-74.5-73.8-73.7 (C-3, C-3’, C-3’’, C-4, C-4’, C-4’’), 73.3-73.0-72.7 (C-2, C-2’, C-2’’), 71.6-71.5-70.9 (C-5, C-5’, C-5’’), 69.7 (C-7), 62.1-62.1-62.0 (C-6, C-6’, C-6’’), 31.3 (C-8).

**HRMS (ESI)**: *m/z* calcd. For $C_{20}H_{39}BrO_{16}N$ $[M+NH_4]^+$: 628.1447; found: 628.1442.

***Compound 26***

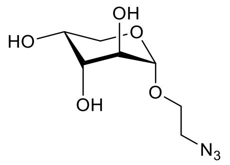

Chemical Formula: $C_7H_{13}N_3O_5$
Exact Mass: 219,09
Molecular Weight: 219,20 g.mol$^{-1}$

Following the protocol **B**, the compound was obtained as a mixture of α/β anomers (90/10) and separate by HPLC as a white solid (α = 0.26 g; β = 0.03 g, 61%).

**$^1$H NMR (400 MHz, $D_2O$)** δ: 5.03 (d, 1H, *J* = 3.6 Hz, H-1), 4.10-4.05 (m, 1H, H-4), 4.01 (dd, 1H, *J* = 12.8, 1.2 Hz, H-5), 3.99-3.88 (m, 2H, H-2, H-3), 3.98-3.91 (m, 1H, H-6), 3.80-3.72 (m, 1H, H-6), 3.74 (dd, 1H, *J* = 12.8, 2.2 Hz, H-5), 3.66 (ddd, 1H, *J* = 13.6, 7.4, 3.0 Hz, H-7), 3.54 (ddd, 1H, *J* = 13.6, 6.0, 3.0 Hz, H-7).

**$^{13}$C NMR (100 MHz, $D_2O$)** δ: 98.9 (C-1), 69.0-68.7-68.2 (C-2, C-3, C-4), 66.8 (C-6), 62.9 (C-5), 50.5 (C-7).

**HRMS (ESI)**: *m/z* calcd. For $C_8H_{14}N_3O_7$ $[M+HCOO]^-$: 264.0837; found: 264.0837.

***Compound 27***

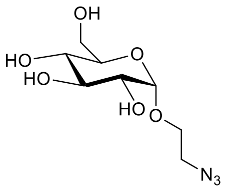

Chemical Formula: $C_8H_{15}N_3O_6$
Exact Mass: 249,10
Molecular Weight: 249,22 g.mol$^{-1}$

Following the protocol **B**, the compound was obtained as a mixture of α/β anomers (75/25) and separate by HPLC as a white solid (α = 0.17 g; β = 0.05 g, 55%).

**$^1$H NMR (400 MHz, $D_2O$)** δ: 5.00 (d, 1H, *J* = 3.8 Hz, H-1), 3.96 (ddd, 1H, *J* = 10.7, 7.4, 3.1 Hz, H-6), 3.91 (dd, 1H, *J* = 12.0, 2.0 Hz, H-7), 3.84-3.71 (m, 4H, H-3, H-4, H-6, H-7), 3.70-3.48 (m, 2H, H-8), 3.60 (dd, 1H, *J* = 9.6, 3.7 Hz, H-2), 3.46 ($t_{app}$, 1H, *J* = 9.5 Hz, H-5).

**$^{13}$C NMR (100 MHz, $D_2O$)** δ: 98.7 (C-1), 73.1 (C-3), 73.0 (C-2), 72.0 (C-4), 69.6 (C-5), 66.9 (C-7), 60.7 (C-6), 50.6 (C-8).

**HRMS (ESI)**: *m/z* calcd. For $C_8H_{19}N_4O_6$ $[M+NH_4]^+$: 267.1299; found: 267.1300.

***Compound 28***

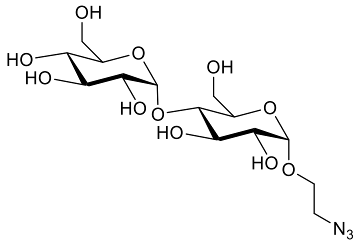


Following the protocol **B**, the compound was obtained as a mixture of α/β anomers (20/80) and separate by HPLC as a white solid (α = 0.18 g; β = 0.68 g, 87%).

**$^1$H NMR (400 MHz, $D_2O$)** δ: 5.29 (d, 1H, *J* = 3.8 Hz, H-1'), 4.88 (d, 1H, *J* = 3.8 Hz, H-1), 3.92 (dd, 1H, *J* = 9.7, 9.5 Hz, H-3 or H-3'), 3.86-3.68 (m, 6H, H-3 or H-3', H-6 or H-6', H-7), 3.67-3.46 (m, 7H, H-2, H-2', H-4, H-4', H-6 or H-6', H-8), 3.41 (ddd, 1H, *J* = 13.5, 5.9, 3.5 Hz, H-8), 3.33 ($t_{app}$, 1H, *J* = 9.4 Hz, H-5).

**$^{13}$C NMR (100 MHz, $D_2O$)** δ: 99.7 (C-1'), 98.1 (C-1), 77.2-72.9 (C-4, C-4'), 73.3-72.6 (C-3, C-3'), 71.8-71.0 (C-2, C-2'), 70.5-69.3 (C-5, C-5'), 66.6 (C-7), 60.5-60.5 (C-6, C-6'), 50.6 (C-8).

**HRMS (ESI):** *m/z* calcd. For $C_{14}H_{25}N_3NaO_{11}$ $[M+Na]^+$: 434.1381; found: 434.1382.

***Compound 29***

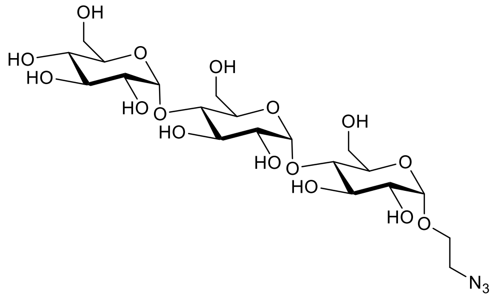


Following the protocol **B**, the compound was obtained as a mixture of α/β anomers (40/60) and separate by HPLC as a white solid (α = 12.0 mg; β = 16.5 mg, 75%).

**$^1$H NMR (400 MHz, $D_2O$)** δ: 5.43 (d, 1H, *J* = 3.8 Hz, H-1''), 5.41 (d, 1H, *J* = 3.9 Hz, H-1'), 5.00 (d, 1H, *J* = 3.8 Hz, H-1), 4.04 (dd, 1H, *J* = 9.8, 9.0 Hz, H-3' or H-3''), 4.00 (dd, 1H, *J* = 9.9, 9.2 Hz, H-3' or -3''), 3.97-3.83 (m, 8H, H-3, H-4 or H-4' or H-4'', H-5', H-5'', H-6 or H-6' or H-6'', H-7), 3.83-3.58 (m, 10H, H-2, H-2', H-4 or H-4' or H-4'', H-6 or H-6' or H-6'', H-7, H-8), 3.53 (ddd, 1H, *J* = 13.6, 6.1, 3.2 Hz, H-8), 3.45 ($t_{app}$, 1H, *J* = 9.4 Hz, H-5).

**$^{13}$C NMR (100 MHz, $D_2O$)** δ: 99.8-99.6 (C-1', C-1''), 98.1 (C-1), 77.4-76.9 (C-4 or C-4' or C-4''), 73.4-73.3-72.9-72.7 (C-3 or C-3' or C-3'' or C-4 or C-4' or C-4''), 71.8-71.6-71.2 (C-2, C-2', C-2''), 71.0-70.5-69.4 (C-5, C-5', C-5''), 66.7 (C-7), 60.5-60.5-60.5 (C-6, C-6'), 50.4 (C-8).

**HRMS (ESI):** *m/z* calcd. For $C_{20}H_{35}N_3NaO_{16}$ $[M+Na]^+$: 596.1910; found: 596.1897

***Compound 5***

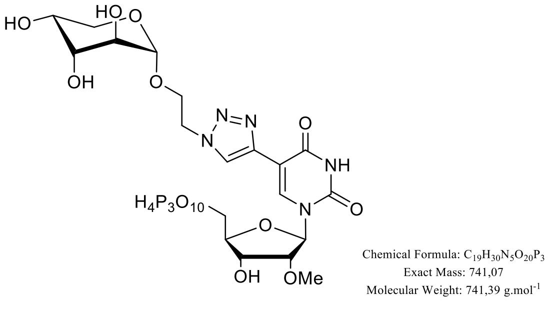


Following the protocol **D**, the compound was obtained as a white solid (0.56 mg, 11%).

**$^1$H NMR (400 MHz, $D_2O$)** δ: 8.47 (s, 1H, H-8), 8.32 (s, 1H, H-6), 6.09 (d, 1H, *J* = 4.9 Hz, H-1’), 4.93 (brs, 1, H-1’’), 4.85-4.74 (m, 2H, H-9), 4.59 ($t_{app}$, 1H, *J* = 5.0 Hz, H-3’), 4.34-4.24 (m, 3H, H-4’, H-5’), 4.13-4.09 (m, 1H, H-10), 3.97 (td, 1H, *J* = 10.7, 3.9 Hz, H-10), 3.85-3.82 (m, 1H, H-4’’), 3.78-3.75 (m, 2H, H-2’’, H-3’’), 3.54 (s, 3H, $H_{Me}$), 3.45 (dd, 1H, *J* = 12.8, 2.0 Hz, H-5’’), 3.15 (dd, 1H, *J* = 12.8, 0.9 Hz, H-5’’).

**$^{13}$C NMR (100 MHz, $D_2O$)** δ: 165.4 (C=O), 151.3 (C=O), 139.1 (C-6), 125.3 (C-8), 106.5 (C-7), 98.5 (C-1’’), 87.9 (C-1’), 83.6 (C-4’), 82.5 (C-2’), 70.5 (C-3’), 68.9 (C-4’’), 68.5-68.2 (C-2’’, C-3’’), 68.0 (C-3’), 66.0 (C-10), 65.2 (C-5’), 62.7 (C-5’’), 58.1 (C-OMe), 50.3 (C-9).

**$^{31}$P NMR (162 MHz, $D_2O$)** δ: -9.3 (s), -11.3 (d, *J* = 19 Hz), -22.2 (s).

**HRMS (ESI):** *m/z* calcd. For $C_{19}H_{30}N_5O_{20}P_3$ $[M+Na]^+$: 764.0592; found: 764.0209.

***Compound 6***

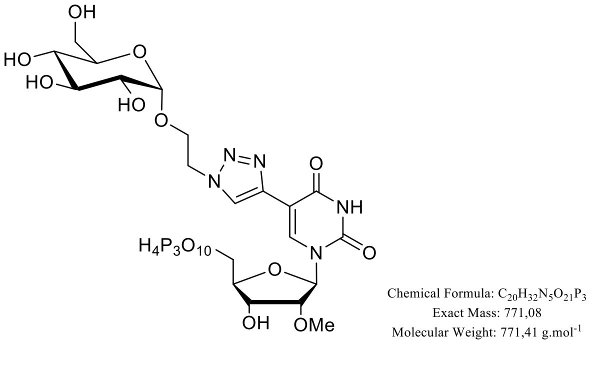


Following the protocol **D**, the compound was obtained as a white solid (1.52 mg, 19%).

**$^1$H NMR (400 MHz, $D_2O$)** δ: 8.47 (s, 1H, H-8), 8.31 (s, 1H, H-6), 6.08 (d, 1H, *J* = 4.8 Hz, H-1’), 4.91 (d, 1, *J* = 3.7 Hz, H-1’’), 4.87-4.71 (m, 2H, H-9), 4.62-4.55 (m, 1H, H-3’), 4.37-4.25 (m, 3H, H-4’, H-5’), 4.22 ($t_{app}$, 1H, *J* = 5.0 Hz, H-2’), 4.18-4.11 (m, 1H, H-10), 4.04-3.98 (m, 1H, H-10), 3.64-3.57 (m, 3H, H-3’’, H-6’’), 3.54 (s, 3H, H-OMe), 3.49 (dd, 1H, *J* = 9.8, 3.8 Hz, H-2’’), 3.32 ($t_{app}$, 1H, *J* = 9.7 Hz, H-4’’), 2.90-2.80 (m, 1H, H-5’’).

**$^{13}$C NMR (100 MHz, $D_2O$)** δ: 162.4 (C=O), 150.3 (C=O), 138.6 (C-6), 125.1 (C-8), 106.5 (C-7), 98.4 (C-1’’), 87.9 (C-1’), 83.6 (C-4’), 82.2 (C-2’), 73.1 (C-3’’), 72.1 (C-5’’), 71.2 (C-2’’), 69.0 (C-4’’), 67.8 (C-3’), 66.0 (C-10), 65.1 (C-5’), 60.1 (C-6’’), 58.5 (C-OMe), 50.5 (C-9).

**$^{31}$P NMR (162 MHz, $D_2O$)** δ: -5.80 (brs), -10.7 (brs), -19.9 (brs).

**HRMS (ESI):** *m/z* calcd. For $C_{20}H_{30}N_5O_{21}P_3$ $[M-2H]^{-2}$: 384.5329; found: 384.5341

***Compound 7***

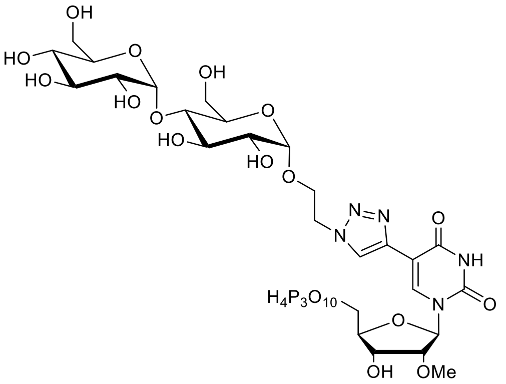

Chemical Formula: $C_{26}H_{42}N_5O_{26}P_3$
Exact Mass: 933,13
Molecular Weight: 933,55 g.mol$^{-1}$

Following the protocol **D**, the compound was obtained as a white solid (4.95 mg, 46%).

**$^1$H NMR (400 MHz, $D_2O$)** δ: 8.51 (s, 1H, H-8), 8.36 (s, 1H, H-6), 6.06 (d, 1H, *J* = 4.4 Hz, H-1'), 5.10 (d, 1H, *J* = 3.8 Hz, H-1''), 4.92 (d, 1H, *J* = 3.5 Hz, H-1''), 4.82-4.76 (m, 2H, H-9), 4.62-4.56 (m, 1H, H-3'), 4.36-4.27 (m, 3H, H-4', H-5'), 4.23 ($t_{app}$, 1H, *J* = 5.0 Hz, H-2'), 4.18-3.94 (m, 2H, H-10), 3.78 ($t_{app}$, 1H, *J* = 9.4 Hz, H-3''), 3.76-3.54 (m, 5H, H-3''', H-6'', H-6'''), 3.56 (s, 3H, H-OMe), 3.54 -3.39 (m, 4H, H-2'', H-2''', H-4'', H-4'''), 3.34 (dd, 1H, *J* = 10.0, 9.4 Hz, H-5'''), 2.59-2.53 (m, 1H, H-5'').

**$^{13}$C NMR (100 MHz, $D_2O$)** δ: 162.7 (C=O), 151.0 (C=O), 138.3 (C-6), 124.7 (C-8), 105.8 (C-7), 100.6 (C-1'''), 97.2 (C-1''), 87.8 (C-1'), 83.4 (C-4'), 82.2 (C-2'), 78.4-73.0-72.9-72.7-72.0-70.9 (C-2'', C-2''', C-3'', C-3''', C-4'', C-4'''), 70.1 (C-5''), 69.0 (C-5'''), 67.9 (C-3'), 65.2 (C-10), 64.9 (C-5'), 60.2-59.8 (C-6'', C-6'''), 58.1 (C-OMe), 50.1 (C-9).

**$^{31}$P NMR (203 MHz, $D_2O$)** δ: -7.7 (s), -11.2 (d, *J* = 15.7 Hz), -21.5 (s)

**HRMS (ESI):** *m/z* calcd. For $C_{26}H_{40}N_5O_{26}P_3$ $[M-2H]^{-2}$: 465.5593; found: 465.5594.

***Compound 8***

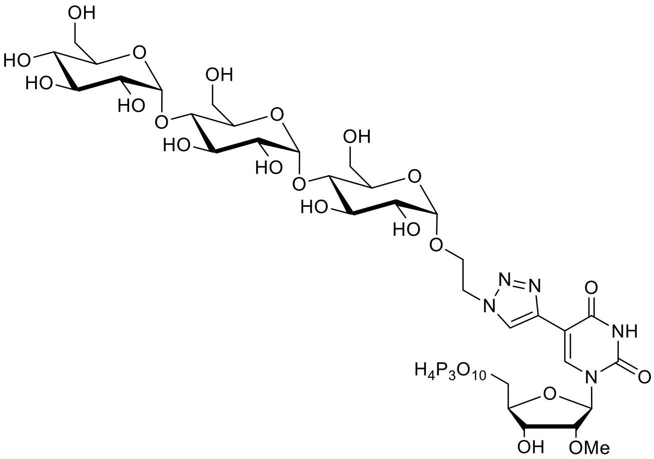

Chemical Formula: $C_{32}H_{52}N_5O_{31}P_3$
Exact Mass: 1095,19
Molecular Weight: 1095,69 g.mol$^{-1}$

Following the protocol **D**, the compound was obtained as a white solid (0.33 mg, 4%).

**$^1$H NMR (400 MHz, $D_2O$)** δ: 8.49 (s, 1H, H-8), 8.34 (s, 1H, H-6), 6.10 (t, 1H, *J* = 4.7 Hz, H-1'), 5.40 (d, 1H, *J* = 3.8 Hz, H-1''''), 5.22 (d, 1H, *J* = 3.9 Hz, H-1'''), 4.91 (d, 1H, *J* = 3.9 Hz, H-1''), 4.79-4.74 (m, 2H, H-9), 4.61-4.57 (m, 1H, H-3'), 4.36-4.29 (m, 3H, H-4', H-5'), 4.23 ($t_{app}$, *J* = 5.1 Hz, H-2'), 4.18-4.10 (m, 1H, H-10), 4.04-3.99 (m, 1H, H-10), 3.89-3.41 (m, 17H, H-2'', H-2''', H-2'''', H-3'', H-3''', H-3'''', H-4'', H-4''', H-4'''', H-5''', H-5'''', H-6'', H-6''', H-6''''), 3.56 (s, 3H, H-OMe), , 2.89-2.83 (m, 1H, H-5'').

**$^{13}$C NMR (100 MHz, $D_2O$)** δ: 162.8 (C=O), 150.9 (C=O), 139.2 (C-5), 138.4 (C-6), 124.8 (C-8), 105.9 (C-7), 99.8-99.6-97.6(C-1'''' C-1''', C-1''), 87.4 (C-1'), 83.5 (C-4'), 82.3 (C-2'), 77.1-76.5-73.3-73.0-72.8-72.7-

71.8-71.8-71.2-71.1-70.1 (C-2’’, C-2’’’, C-2’’’’, C-3’’, C-3’’’, C-3’’’’, C-4’’, C-4’’’, C-4’’’’, C-5’’’, C-5’’’’), 70.1-69.3 (C-3’, C-5’’), 67.9 (C-10), 65.8 (C-5’), 60.5-60.2-59.9 (C-6’’, C-6’’’, C-6’’’’), 58.1 (C-OMe), 50.1 (C-9).

**$^{31}$P NMR (162 MHz, $D_2O$)** δ: -10.7 (s), -11.4 (d, *J* = 20.0 Hz), -23.1 (s)

**HRMS (ESI):** *m/z* calcd. For $C_{32}H_{50}N_5O_{31}P_3$ $[M-2H]^{-2}$: 546.5857; found: 546.5853.

#### 4. NMR spectra (1H, 13C, 31P) of all compounds

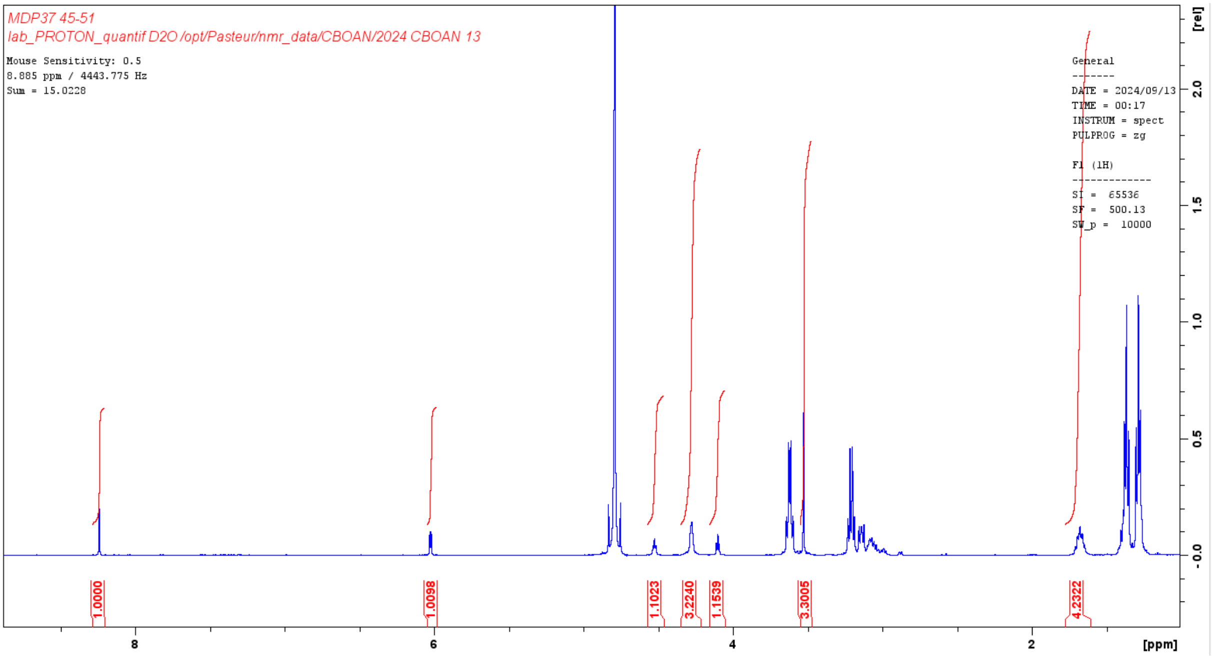


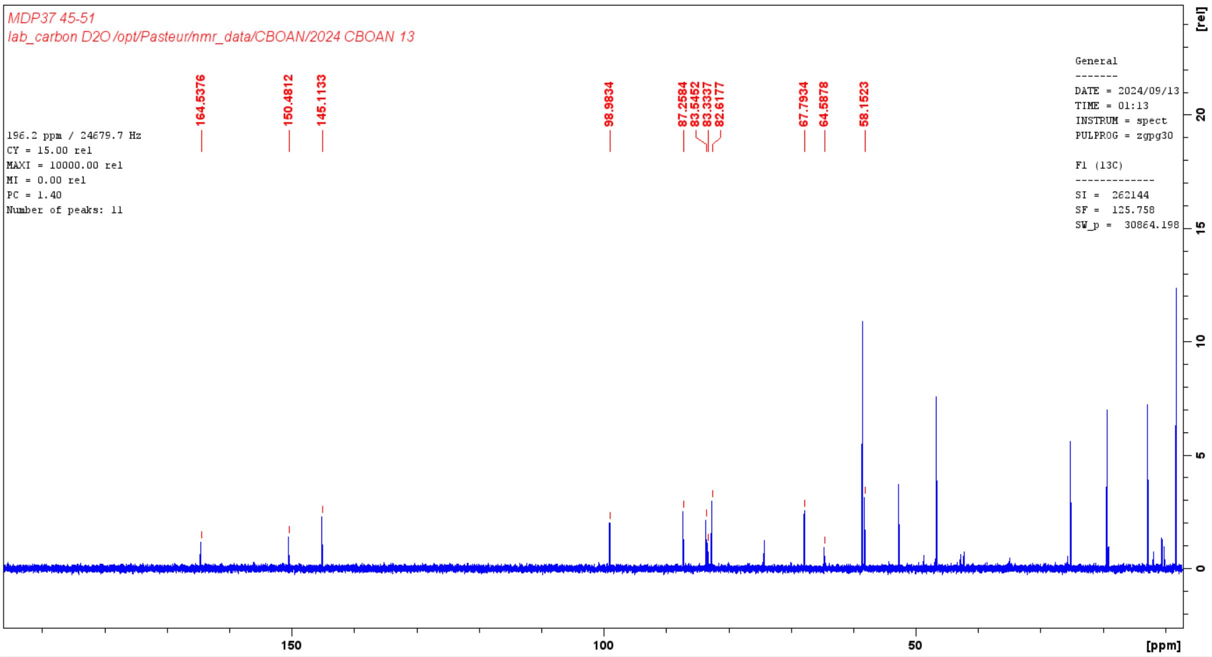

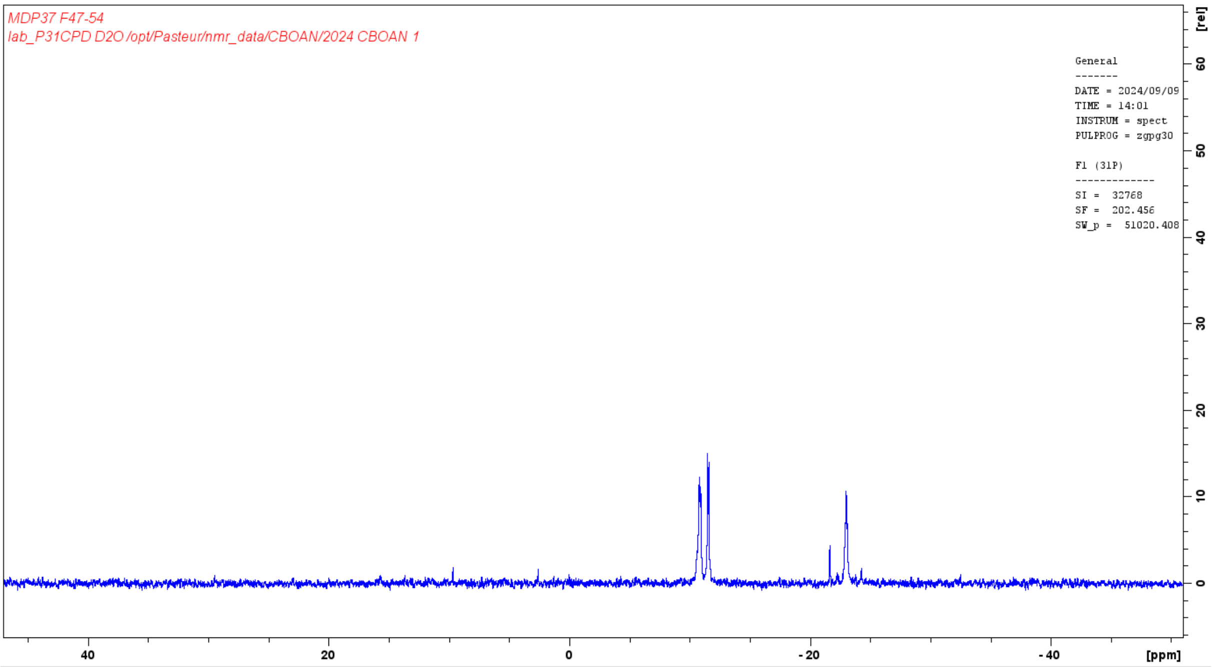


NMR spectrum of **5-ethynyl-2’-*O*-methyl-UTP**

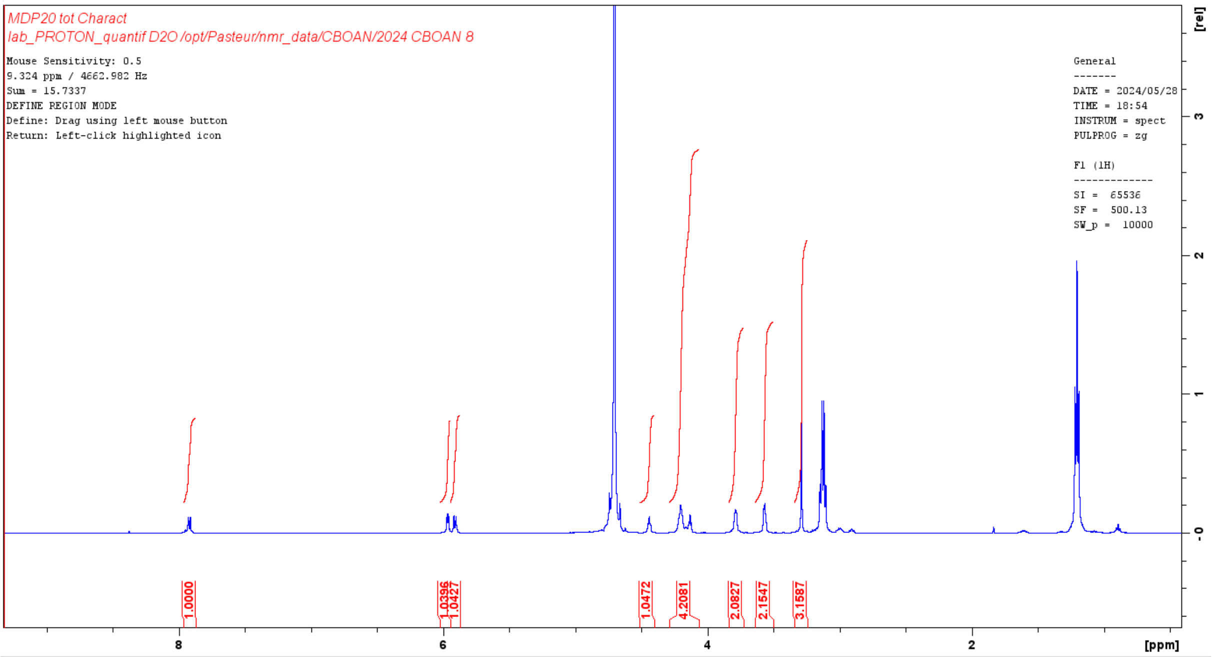

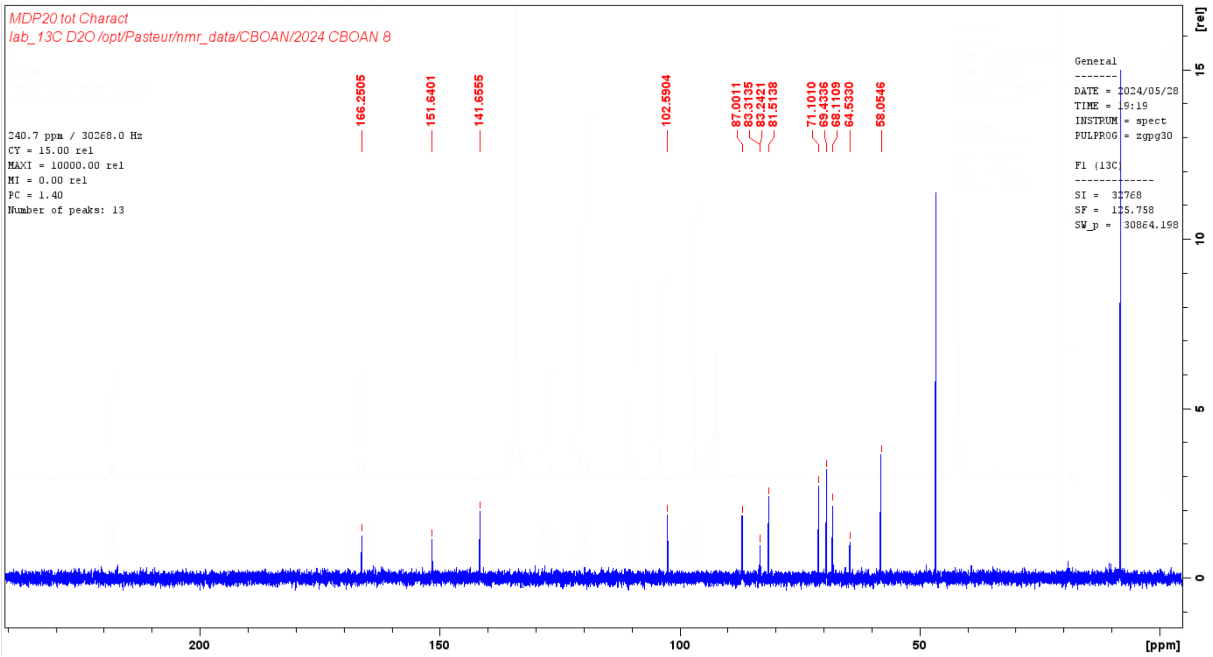

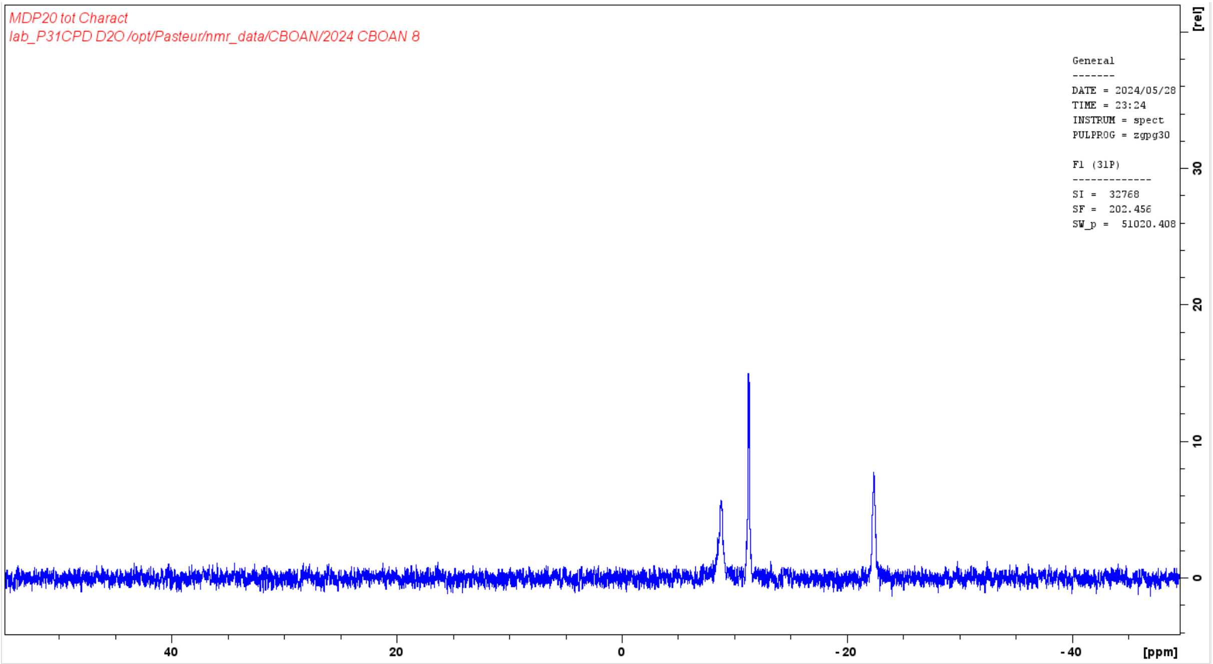


NMR spectrum of **2'-*O*-(2-Methoxyethyl)-uridine triphosphorylation**

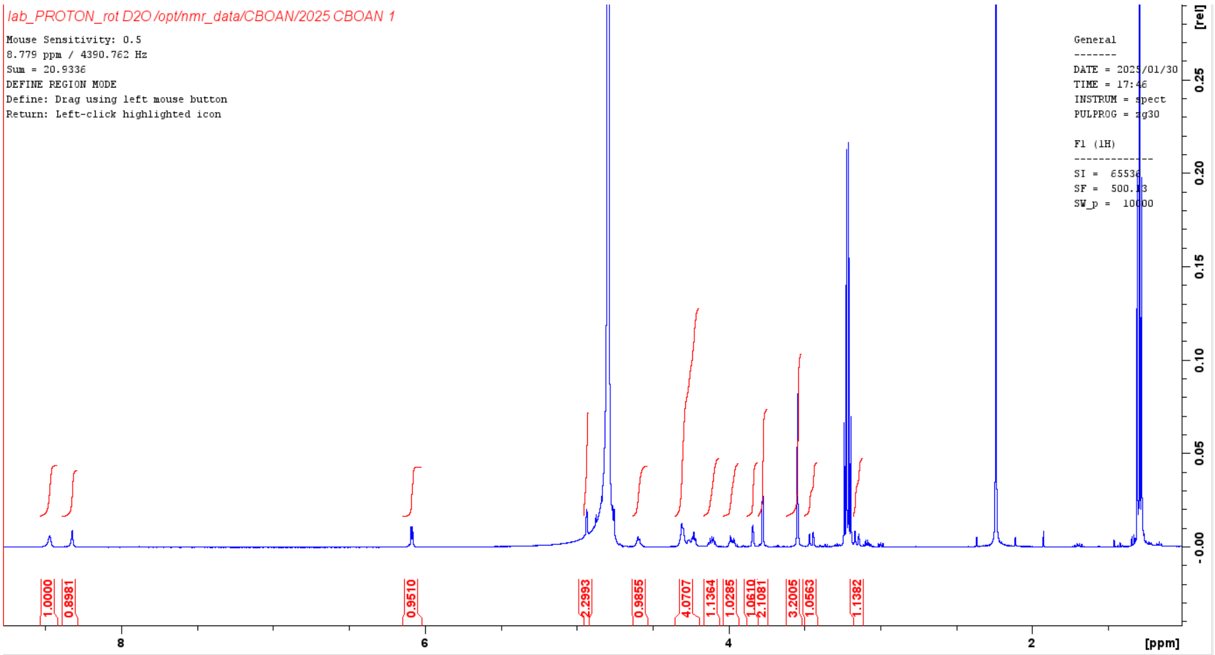


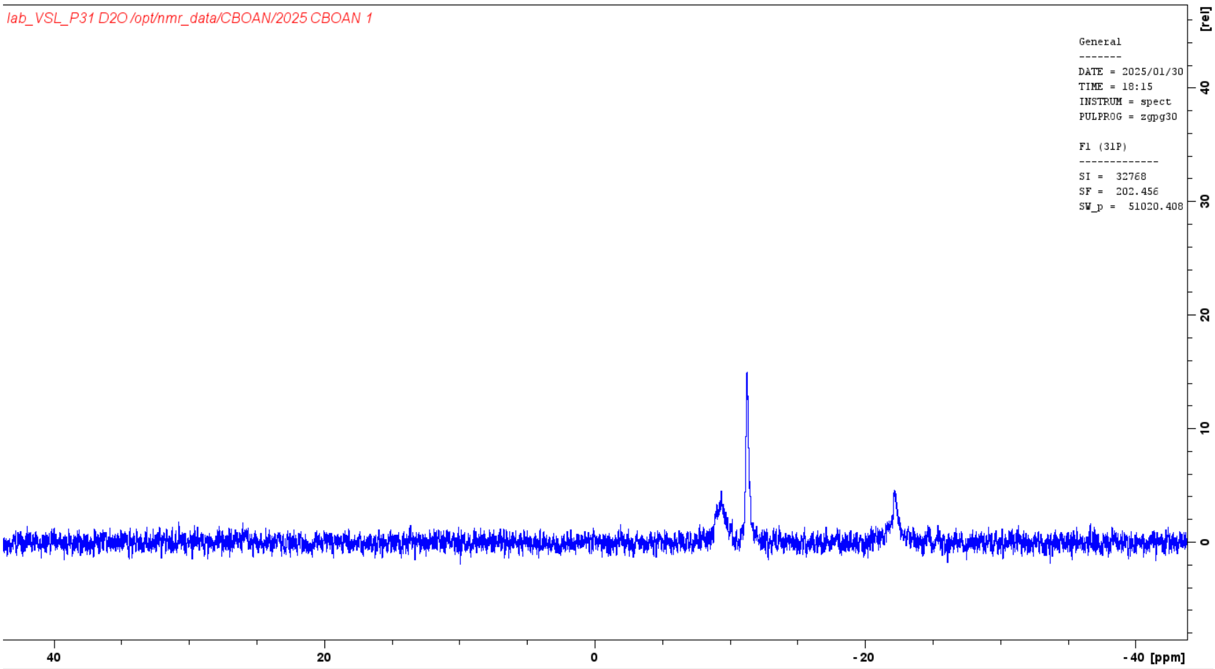


NMR spectrum of **5**

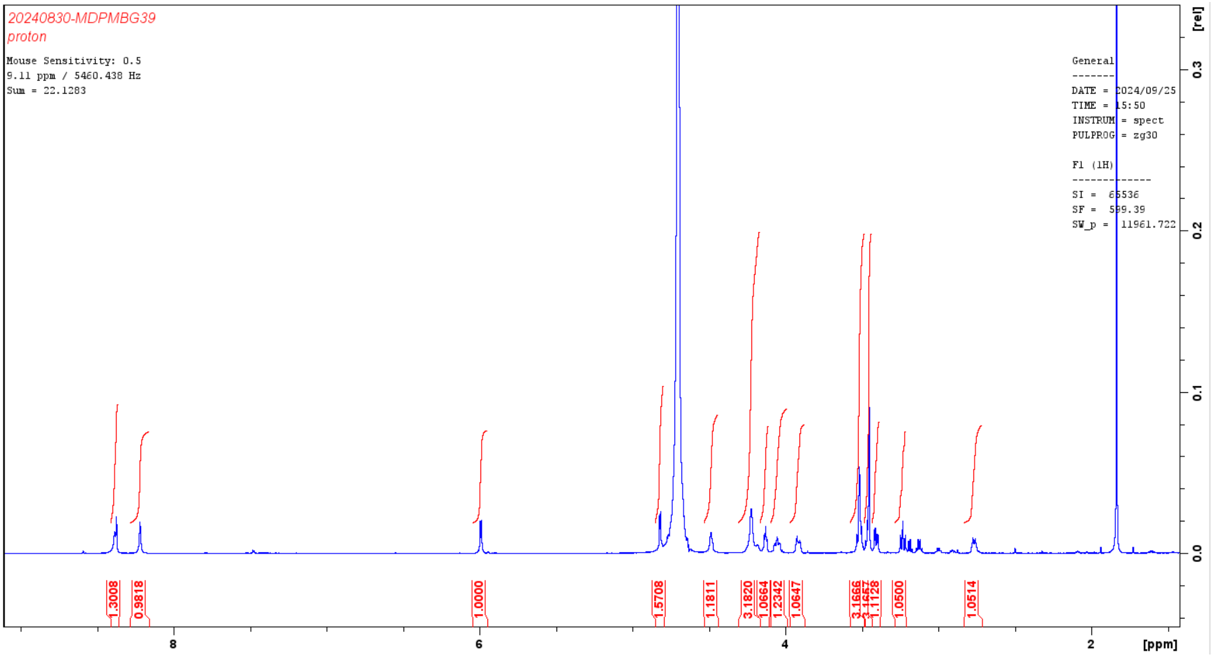
20240830-MDPMBG39
proton
Mouse Sensitivity: 0.5
9.11 ppm / 5460.438 Hz
Sum = 22.1283
General
DATE = 2024/09/25
TIME = 15:50
INSTRUM = spect
PULPROG = zg30
F1 (1H)
SI = 65536
SF = 599.39
SW_p = 11961.722
1.3008
0.9818
1.0000
1.5708
1.1811
3.1820
1.0664
1.2342
1.0647
3.1666
1.1128
1.0500
1.0514
[rel]
[ppm]

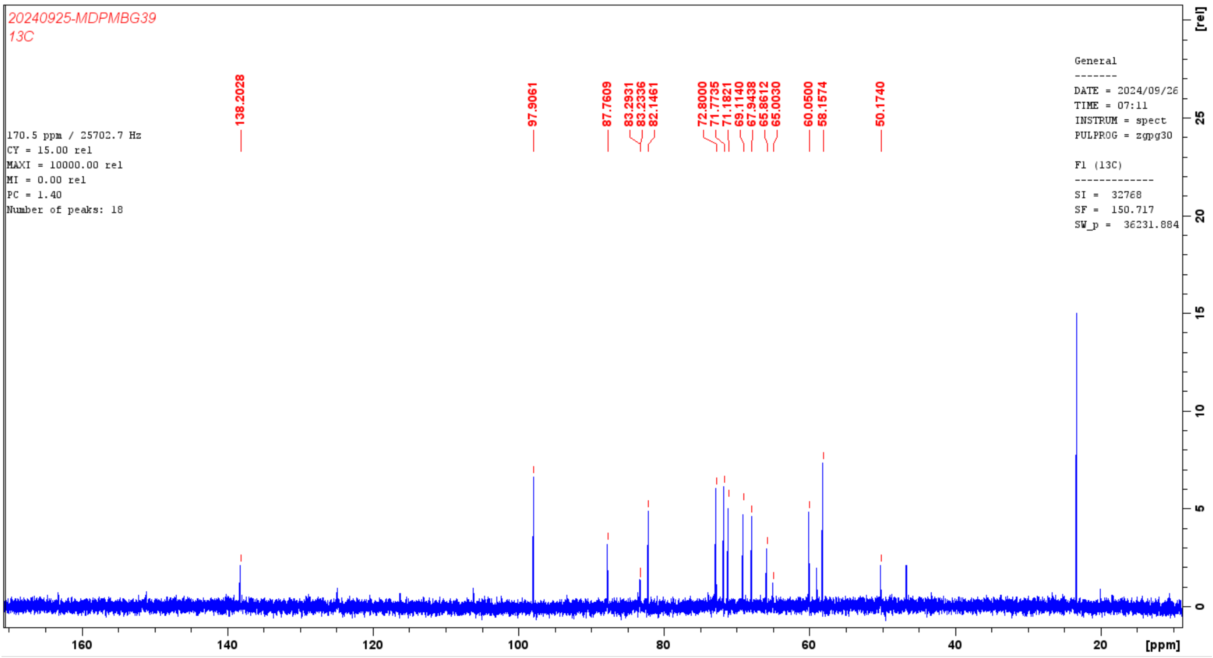
20240925-MDPMBG39
13C
170.5 ppm / 25702.7 Hz
CY = 15.00 rel
MAXI = 10000.00 rel
MI = 0.00 rel
PC = 1.40
Number of peaks: 18
General
DATE = 2024/09/26
TIME = 07:11
INSTRUM = spect
PULPROG = zgpg30
F1 (13C)
SI = 32768
SF = 150.717
SW_p = 36231.884
138.2028
97.9061
87.7609
83.2931
83.2336
82.1461
72.8000
71.7735
71.1821
69.1140
67.9438
65.8612
65.0030
60.0500
58.1574
50.1740
[rel]
[ppm]

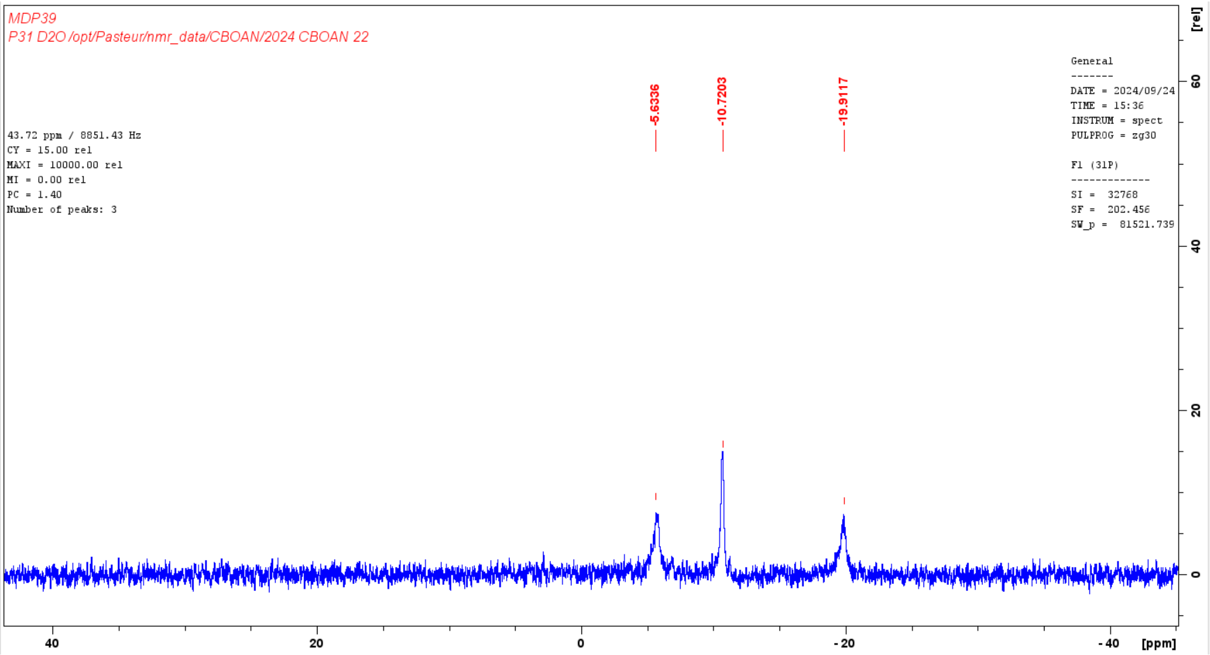


NMR spectrum of **6**

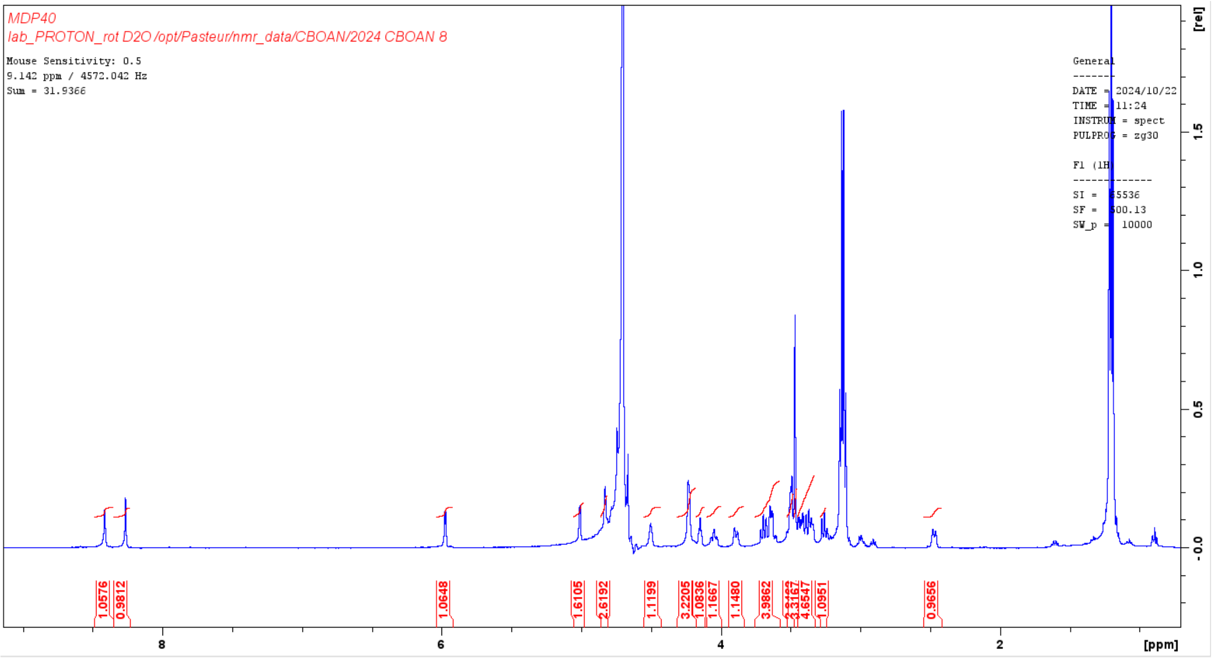

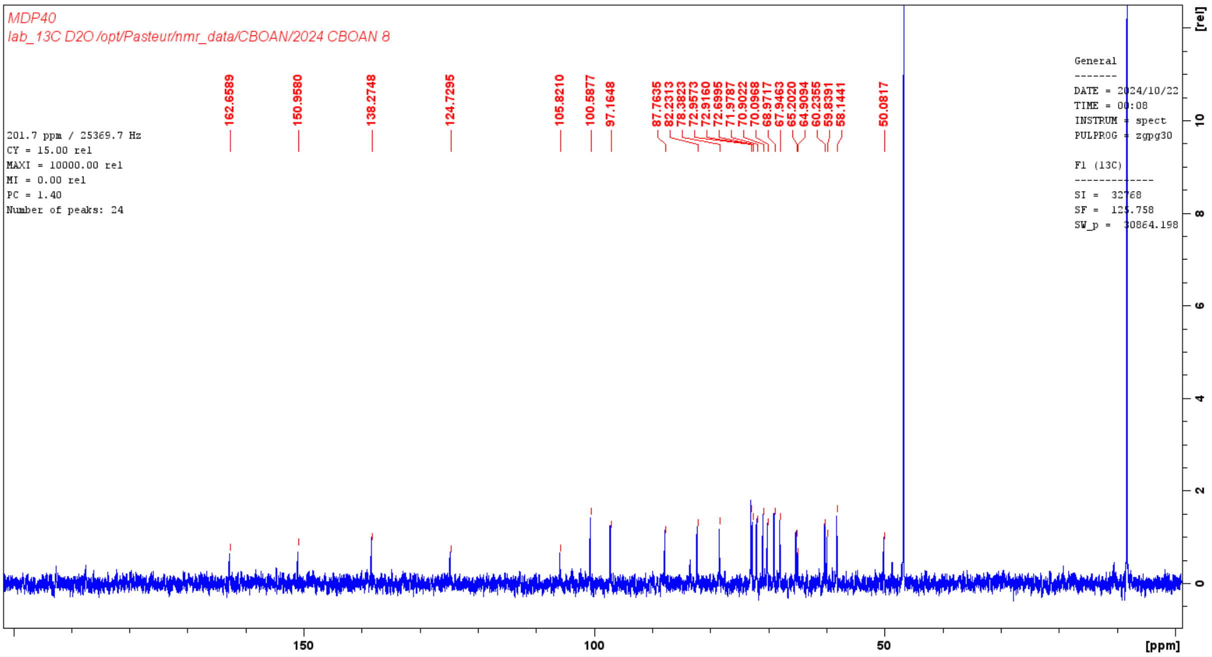


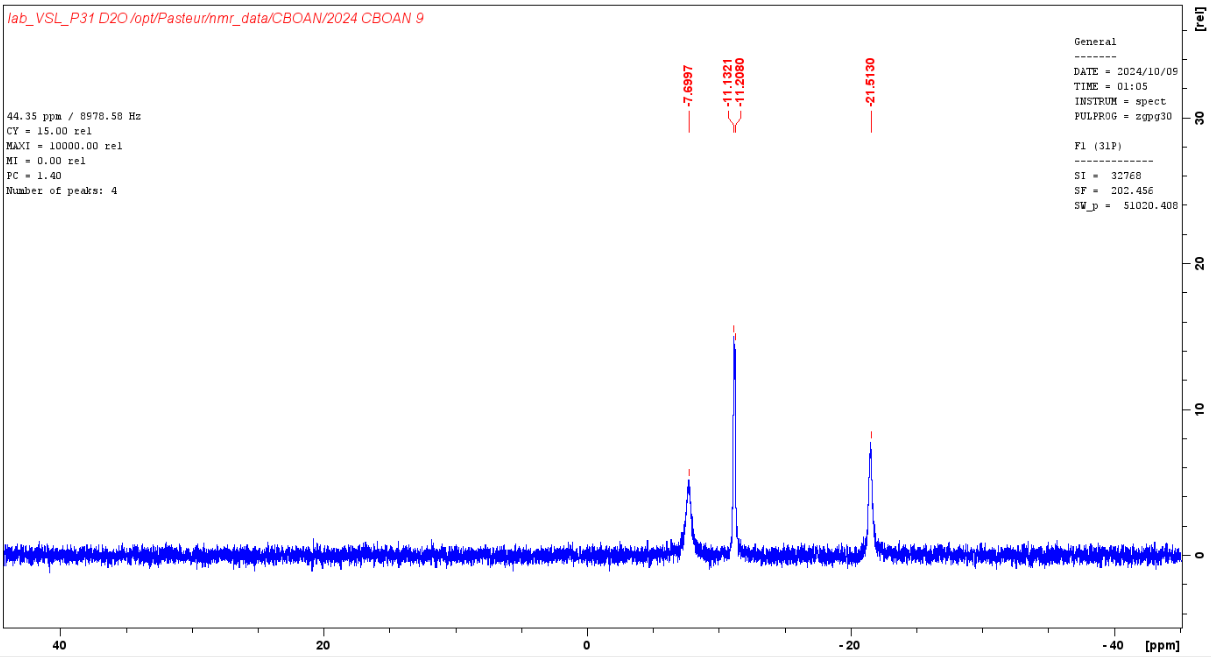


NMR spectrum of **7**

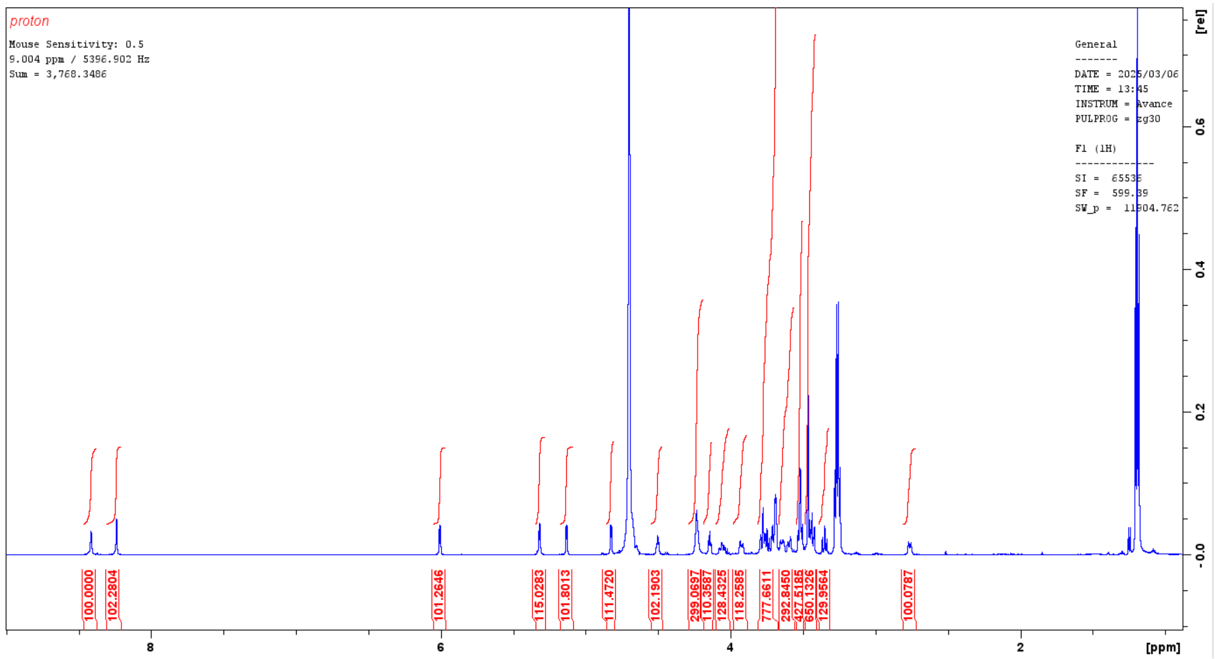

proton
Mouse Sensitivity: 0.5
9.004 ppm / 5396.902 Hz
Sum = 3,768.3486
General
DATE = 2025/03/06
TIME = 13:45
INSTRUM = Avance
PULPROG = zg30
F1 (1H)
SI = 65536
SF = 599.89
SW_p = 11904.762
100.0000
102.2804
101.2646
115.0283
101.8013
111.4720
102.1903
299.0697
110.3587
128.4325
118.2585
777.6611
292.8450
427.5185
650.1326
129.9564
100.0787
[ppm]
[rel]


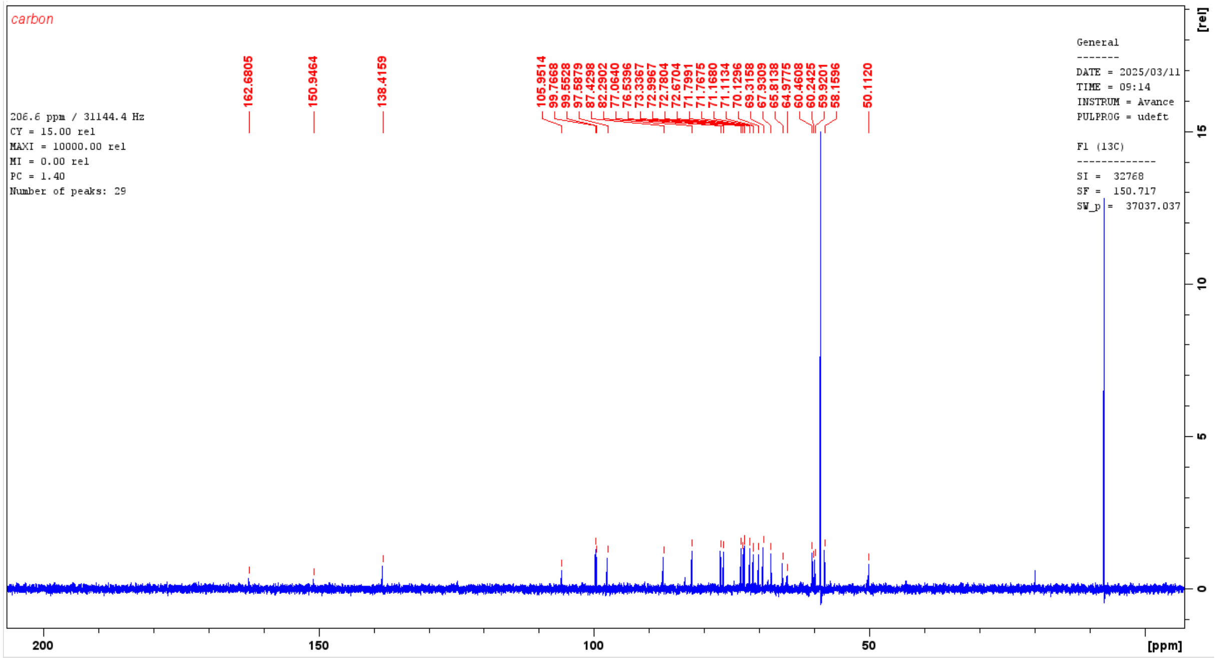

carbon
206.6 ppm / 31144.4 Hz
CY = 15.00 rel
MAXI = 10000.00 rel
MI = 0.00 rel
PC = 1.40
Number of peaks: 29
General
DATE = 2025/03/11
TIME = 09:14
INSTRUM = Avance
PULPROG = udeft
F1 (13C)
SI = 32768
SF = 150.717
SW_p = 37037.037
162.6805
150.9464
138.4159
105.9514
99.7668
99.5528
97.5879
87.4298
82.2902
77.0640
76.5396
73.3367
72.9967
72.7804
72.6704
71.7991
71.7675
71.1680
71.1134
70.1296
69.3158
67.9309
65.8138
64.9775
60.4608
60.2425
59.9201
58.1596
50.1120
[ppm]
[rel]

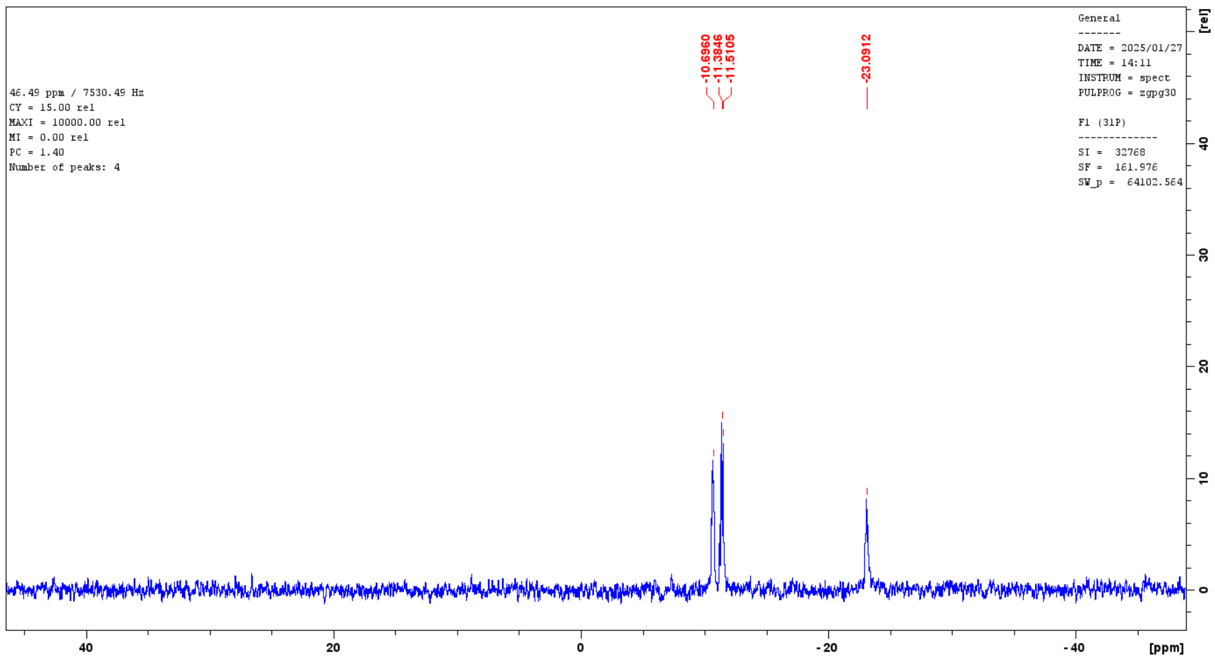

NMR spectrum of **8**

**Protein sequences**

***Thermococcus gorgonarius*** **(Tgo) DGLK protein sequence**

MILDTDYITEDGKPVIRIFKKENGEFKIDYDRNFEPYIYALLKDDSAIEDVKKITAERHGTTVRVVRAEKV
KKKFLGRPIEVWKLYFTHPQDQPAIRDKIKEHPAVVDIYEYDIPFAKRYLIDKGLIPMEGDEELKMLAFAI
ATLYHEGEEFAEGPILMISYADEEGARVITWKNIDLPYVDVVSTEKEMIKRFLKVVKEKDPDVLITYNGDN
FDFAYLKKRSEKLGVKFILGREGSEPKIQRMGDRFAVEVKGRIHFDLYPVIRRTINLPTYTLEAVYEAIFG
QPKEKVYAEEIAQAWETGEGLERVARYSMEDAKVTYELGKEFFPMEAQLSRLVGQSLWDVSRSSTGNLVEW
FLLRKAYERNELAPNKPDERELARRRESYAGGYVKEPERGLWENIVYLDFRSLGPSIIITHNVSPDTLNRE
GCEEYDVAPQVGHKFCKDFPGFIPSLLGDLLEERQKVKKKMKATIDPIEKKLLDYRQRLIKILANSFYGYY
GYAKARWYCKECAESVTAWGRQYIETTIREIEEKFGFKVLYADTDGFFATIPGADAETVKKKAKEFLDYIN
AKLPGLLELEYEGFYKRGFFVTKKKYAVIDEEDKITTRGLEIVRRDWSEIAKETQARVLEAILKHGDVEEA
VRIVKEVTEKLSKYEVPPEKLVIYKQITRDLKDYKATGPHVAVAKRLAARGIKIRPGTVISYIVLKGSGRI
GDRAIPFDEFDPAKHKYDAEYYIENQVLPAVERILRAFGYRKEDLRYQKTRQVGLGAWLKPKT

***Thermococcus sp. 9°N*** **(9°N) DGLK protein sequence**

MILDTDYITENGKPVIRVFKKENGEFKIEYDRTFEPYFYALLKDDSAIEDVKKVTAKRHGTVVKVKRAEKV
QKKFLGRPIEVWKLYFNHPQDQPAIRDRIRAHPAVVDIYEYDIPFAKRYLIDKGLIPMEGDEELTMLAFAI
ATLYHEGEEFGTGPILMISYADGSEARVITWKKIDLPYVDVVSTEKEMIKRFLRVVREKDPDVLITYNGDN
FDFAYLKKRCEELGIKFTLGRDGSEPKIQRMGDRFAVEVKGRIHFDLYPVIRRTINLPTYTLEAVYEAVFG
KPKEKVYAEEIAQAWESGEGLERVARYSMEDAKVTYELGREFFPMEAQLSRLIGQSLWDVSRSSTGNLVEW
FLLRKAYKRNELAPNKPDERELARRRGGYAGGYVKEPERGLWDNIVYLDFRSLGPSIIITHNVSPDTLNRE
GCKEYDVAPEVGHKFCKDFPGFIPSLLGDLLEERQKIKRKMKATVDPLEKKLLDYRQRLIKILANSFYGYY
GYAKARWYCKECAESVTAWGREYIEMVIRELEEKFGFKVLYADTDGLHATIPGADAETVKKKAKEFLKYIN
PKLPGLLELEYEGFYVRGFFVTKKKYAVIDEEGKITTRGLEIVRRDWSEIAKETQARVLEAILKHGDVEEA
VRIVKEVTEKLSKYEVPPEKLVIHKQITRDLRDYKATGPHVAVAKRLAARGVKIRPGTVISYIVLKGSGRI
GDRAIPADEFDPTKHRYDAEYYIENQVLPAVERILKAFGYRKEDLRYQKTKQVGLGAWLKVKGK

***Pyrococcus sp. GB-D DV*** **(DV) DGLK protein sequence**

MILDADYITEDGKPIIRIFKKENGEFKVEYDRNFRPYIYALLKDDSQIDEVRKITAERHGKIVRIIDAEKV
RKKFLGRPIEVWRLYFEHPQDQPAIRDKIREHSAVIDIFEYDIPFAKRYLIDKGLIPMEGDEELKLLAFAI
ATLYHEGEEFAKGPIIMISYADEEEAKVITWKKIDLPYVEVVSSEREMIKRFLKVIREKDPDVIITYNGDS
FDLPYLVKRAEKLGIKLPLGRDGSEPKMQRLGDMTAVEIKGRIHFDLYHVIRRTINLPTYTLEAVYEAIFG
KPKEKVYAHEIAEAWETGKGLERVAKYSMEDAKVTYELGREFFPMEAQLSRLVGQPLWDVSRSSTGNLVEW
YLLRKAYERNELAPNKPDEREYERRLRESYAGGYVKEPEKGLWEGLVSLDFRSLGPSIIITHNVSPDTLNR
EGCREYDVAPEVGHKFCKDFPGFIPSLLKRLLDERQEIKRKMKASKDPIEKKMLDYRQRLIKILANSYYGY
YGYAKARWYCKECAESVTAWGREYIEFVRKELEEKFGFKVLYIDTDGLYATIPGAKPEEIKKKALEFVDYI
NAKLPGLLELEYEGFYVRGFFVTKKKYALIDEEGKIITRGLEIVRRDWSEIAKETQAKVLEAILKHGNVEE
AVKIVKEVTEKLSKYEIPPEKLVIYKQITRPLHEYKAIGPHVAVAKRLAARGVKVRPGMVIGYIVLRGDGP
ISKRAILAEEFDLRKHKYDAEYYIENQVLPAVLRILEAFGYRKEDLRWQKTKQTGLTAWLNIKKK

##### *Thermococcus kodakarensis* (Kod) DGLK protein sequence

MILDTDYITEDGKPVIRIFKKENGEFKIEYDRTFEPYFYALLKDDSAIEEVKKITAERHGTVVTVKRVEKV
QKKFLGRPVEVWKLYFTHPQDQPAIRDKIREHPAVIDIYEYDIPFAKRYLIDKGLVPMEGDEELKMLAFAI
ATLYHEGEEFAEGPILMISYADEEGARVITWKNVDLPYVDVVSTEREMIKRFLRVVKEKDPDVLITYNGDN
FDFAYLKKRCEKLGINFALGRDGSEPKIQRMGDRFAVEVKGRIHFDLYPVIRRTINLPTYTLEAVYEAVFG
QPKEKVYAEEITTAWETGENLERVARYSMEDAKVTYELGKEFLPMEAQLSRLIGQSLWDVSRSSTGNLVEW
FLLRKAYERNELAPNKPDEKELARRRQSYEGGYVKEPERGLWENIVYLDFRSLGPSIIITHNVSPDTLNRE
GCKEYDVAPQVGHRFCKDFPGFIPSLLGDLLEERQKIKKKMKATIDPIERKLLDYRQRLIKILANSYYGYY
GYARARWYCKECAESVTAWGREYITMTIKEIEEKYGFKVIYSDTDGFFATIPGADAETVKKKAMEFLKYIN
AKLPGALELEYEGFYKRGFFVTKKKYAVIDEEGKITTRGLEIVRRDWSEIAKETQARVLEALLKDGDVEKA
VRIVKEVTEKLSKYEVPPEKLVIHKQITRDLKDYKATGPHVAVAKRLAARGVKIRPGTVISYIVLKGSGRI
GDRAIPFDEFDPTKHKYDAEYYIENQVLPAVERILRAFGYRKEDLRYQKTRQVGLSAWLKPKGT

##### G2 protein sequence

MRGSHHHHHHTDPSGLVPRGSMILDTDYITEDGKPVIRIFKKENGEFKIEYDRTFEPYFYALLKDDSAIED
VKKITAERHGTTVRVVRAEKVKKKFLGRPIEVWRLYFEHPQDQPAIRDKIREHSAVIDIFEYDIPFAKRYL
IDKGLIPMEGDEELKLLAFAIATLYHEGEEFAEGPILMISYADEEGARVITWKNIDLPYVDVVSTEKEMIK
RFLKVVKEKDPDVLITYNGDNFDFAYLKKRSEKLGVKFILGREGSEPKIQRMGDRFAVEVKGRIHFDLYPV
IRRTINLPTYTLEAVYEAIFGQPKEKVYAEEITTAWETGENLERVARYSMEDAKVTYELGKEFFPMEAQLS
RLVGQSLWDVSRSSTGNLVEWFLLRKAYERNELAPNKPDEKELARRRQSYEGGYVKEPERGLWENIVYLDF
RSLGPSIIITHNVSPDTLNREGCKEYDVAPQVGHRFCKDFPGFIPSLLGDLLEERQKIKRKMKATVDPLEK
KLLDYRQRLIKILANSFYGYYGYAKARWYCKECAESVTAWGRQYIETTIREIEEKFGFKVLYADTDGFFAT
IPGADAETVKKKAKEFLDYINAKLPGLLELEYEGFYKRGFFVTKKKYAVIDEEDKITTRGLEIVRRDWSEI
AKETQARVLEAILKHGDVEEAVRIVKEVTEKLSKYEVPPEKLVIYKQITRDLKDYKATGPHVAVAKRLAAR
GIKIRPGTVISYIVLKGSGRIGDRAIPFDEFDPAKHKYDAEYYIENQVLPAVERILRAFGYRKEDLRYQKT
RQVGLGAWLKPKT

**Nanopore sequencing data processing**

1. ***Synthetic reference sequence generation***

The four full-length construct reference sequences, each including a Unique Molecular Identifier (UMI) pair, were aligned using MUSCLE v5.1[5]. A consensus sequence was generated from this alignment by selecting the most frequent nucleotide at each position, with ties broken arbitrarily. This synthetic sequence of 2751 bp served as the alignment target for the Nanopore reads, providing a reference equidistant from all four constructs and thus avoiding alignment bias toward any single variant.

2. ***Basecalling and mapping***

Oxford Nanopore Technologies (ONT) raw signal files were basecalled using Dorado v1.2.0 (https://github.com/nanoporetech/dorado; model dna_r10.4.1_e8.2_400bps_sup@v5.2.0) with default parameters. Reads were aligned to the synthetic reference sequence using minimap2 v2.28, integrated within Dorado[6,7].

3. ***Read filtering***

Aligned reads were filtered using SAMtools v1.21[8]. Only primary alignments with MAPQ > 20 and read length within 2751 ± 20% bp (i.e., 2201–3301 bp) were retained. These criteria remove low-quality mappings and reads that are unusually short or long, including potential concatemer artifacts.

4. ***UMI extraction***

UMI pairs were extracted from filtered alignments at reference positions 121–164 and 2603–2646 using our in-house tool xumi v1.0.3 (https://github.com/Fravadona/xumi, doi:10.5281/zenodo.18905757). Only pairs where both UMIs were 44 ± 6 bp were retained, concatenated, and written as FASTA sequences identified by their source read's query name; these steps were performed using a custom AWK script.

5. ***UMI clustering***

Concatenated UMI sequences were clustered using CD-HIT-EST v4.8.1[9] at 91.5% sequence identity, in accurate mode (-g 1) and with full-length sequence identifiers preserved (-d 0). Given the $3^{64}$ sequence space of the UMI pair design (64 degenerate bases out of 88, each allowing 3 possible nucleotides), the probability of incorrectly merging distinct UMIs at this threshold is vanishingly small relative to experimental library sizes. Only clusters with depth ≥ 5 were retained.

6. ***Cluster reads extraction***

Reads belonging to each retained cluster were extracted from the alignment file using a custom Python script. The resulting alignments were organized into a single BAM file in which each cluster was represented as a separate synthetic contig reference, allowing independent variant calling. The BAM file was then sorted and indexed, and a companion FASTA file was generated by copying the synthetic reference sequence once per contig.

7. ***Variant calling***

Variants were called from the clustered alignments using FreeBayes v1.3.10[10] with default parameters, except that ploidy was set to 1, since each cluster represents reads derived from a single UMI-tagged molecule.

8. ***Variant filtering and normalization***

The VCF file was processed using BCFtools v1.21[8] and a custom AWK script. At positions with multiple candidate alleles, the best-supported allele was retained based on alternate allele observation count (AO), using QA (the sum of Phred-scaled base quality scores for the alternate allele) as a tiebreaker. Only variants where AO exceeded the reference allele observation count (RO) (i.e., AO > RO) and that were located within the mutagenized target region (194–2584) were retained.

##### 9. Consensus sequence generation

Consensus sequences for each cluster were reconstructed from the filtered VCF file using BCFtools consensus. The consensus sequences were then trimmed to retain only the region of interest (reference positions 194–2584). Sequences whose length was not divisible by 3 were discarded. The remaining consensus sequences were translated into protein sequences using the standard genetic code with a custom Python script. The generated protein sequences were used as input in downstream analyses.

#### Supplementary Figures

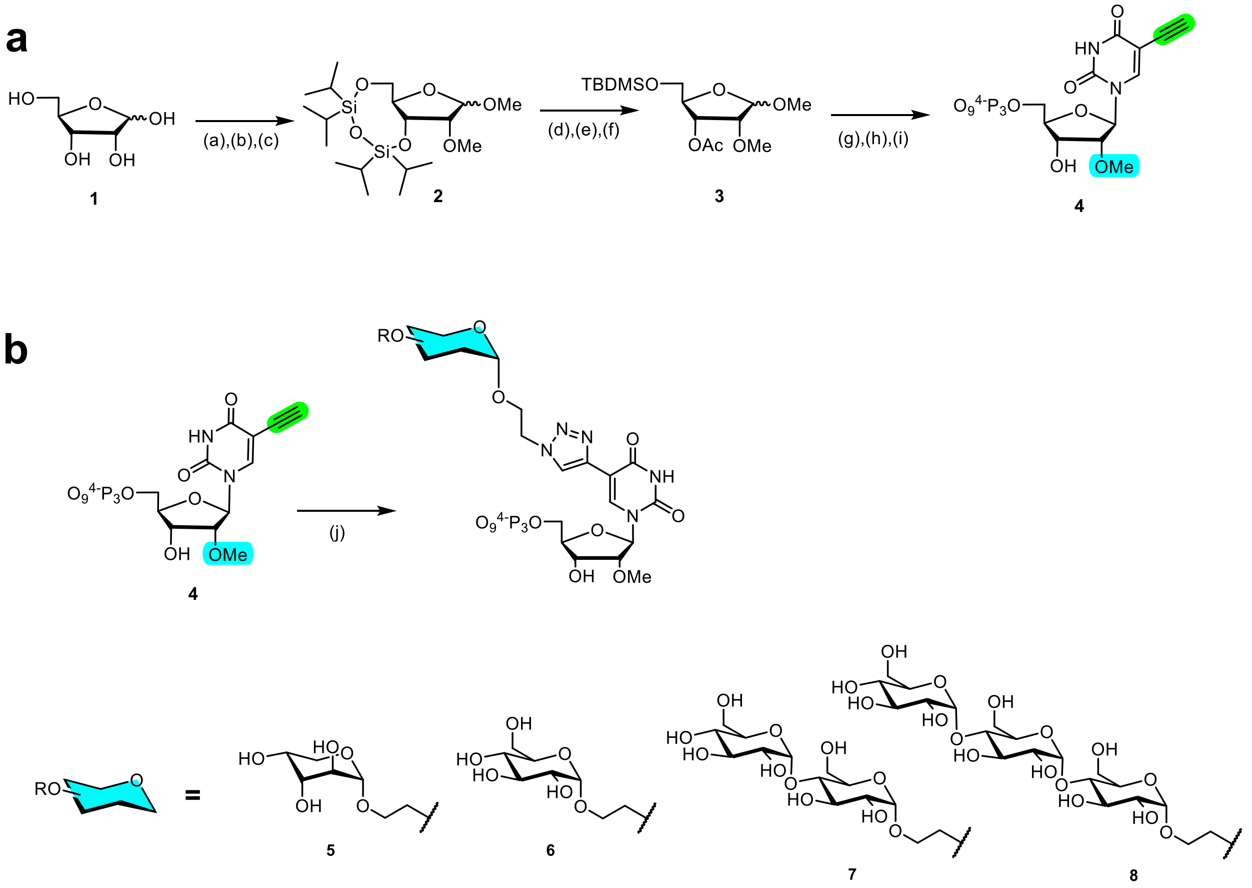


**Supplementary Figure 1 | Synthesis of modified nucleotides.**

**(a)** Multi-step synthesis of 5-ethynyl-2'-*O*-methyl-UTP **4** starting from D-ribose in 9 steps and via intermediates **2** and **3**. Reagents and conditions: (a) Dowex ($H^+$), MeOH, M.W., 100°C, 40 min, 98%; (b) $TIPDSiCl_2$, pyr., rt, 1 h, 97%; (c) MeI, NaH, rt, 5 h, 53%; (d) TBAF·$3H_2O$, THF, rt, 2 h, 83%; (e) TBDMS-Cl, TEA, DMAP, DMF, rt, overnight, 55%; (f) $AC_2O$, DMAP, pyr., rt, overnight, 96%; (g) i) BSTFA, ACN, rt, 30 min, ii) TMSOTf, ACN, 80°C, 24 h, 57%; (h) TBAF (1M/THF), THF, 0°C , 3 h, 77%; (i) 1) 2-chloro-1,3,2-benzodioxaphosphorin-4-one, pyridine, dioxane, rt, 5 h; 2) $(nBu_3NH)_2H_2P_2O_7$, DMF, $Bu_3N$, rt, 45 min; 3) $I_2$, pyr./$H_2O$,rt ,30 min; 4) aq. $NH_3$, rt, 1.5 h, 16 % over four steps.

**(b)** Synthesis of carbohydrate-modified nucleotides. *Reagents and conditions:* **(j)** azidoethyl glycosides[11], CuI, DMF/water, 25°C, 4 h.

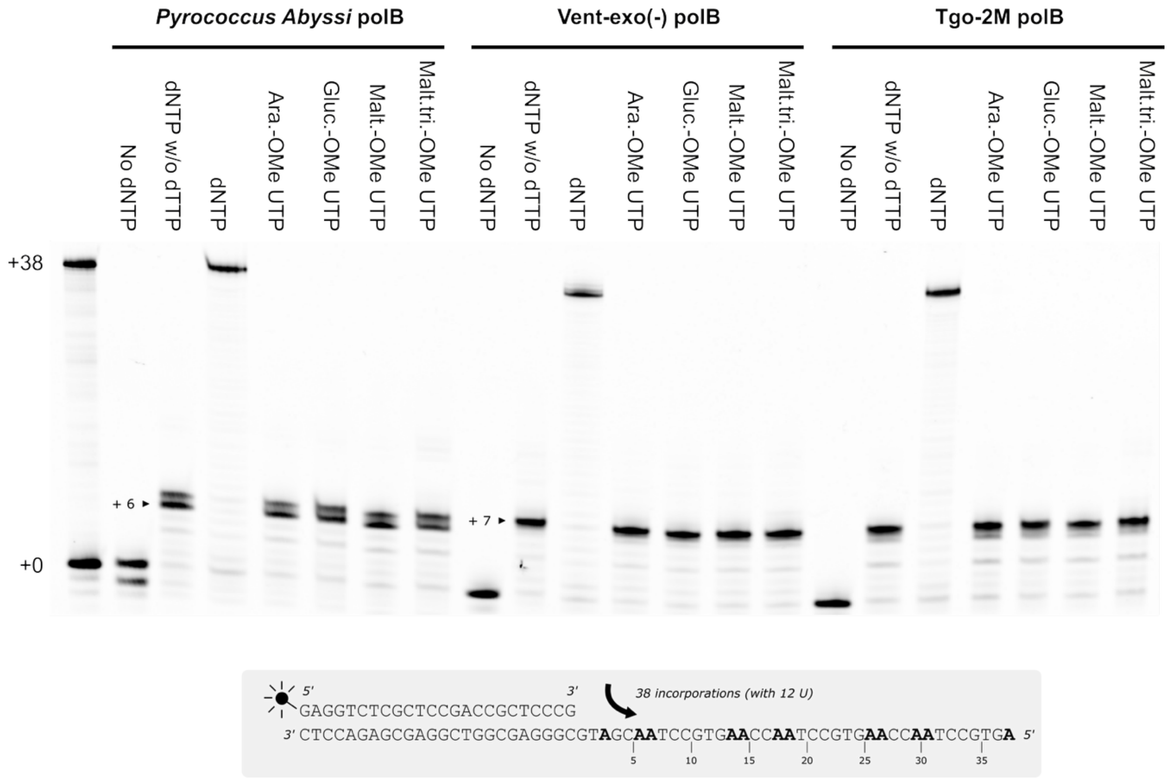


**Supplementary Figure 2 | Basal activity for glycosylated nucleotides incorporation by various family B DNA polymerases from the literature.**

Three family B DNA polymerases, the DNAP from *Pyrococcus abyssi*, the Vent(exo$^{-}$), and the Tgo-2M variant identified in Freund et al.[12], were tested for incorporation of glycosylated nucleotides (5-arabinosyl-2'-*O*-methyl-UTP, 5-glucosyl-2'-*O*-methyl -UTP, 5-maltosyl-2'-*O*-methyl -UTP or 5-maltotriosyl-2'-*O*-methyl-UTP). None of these polymerases could incorporate more than three modified nucleotides (+6 or +7 total), including 2 in a row, and this was with a high risk of errors, as activity was similar with mismatched nucleotides in the absence of dTTP. All polymerases showed full activity in the dNTP control.

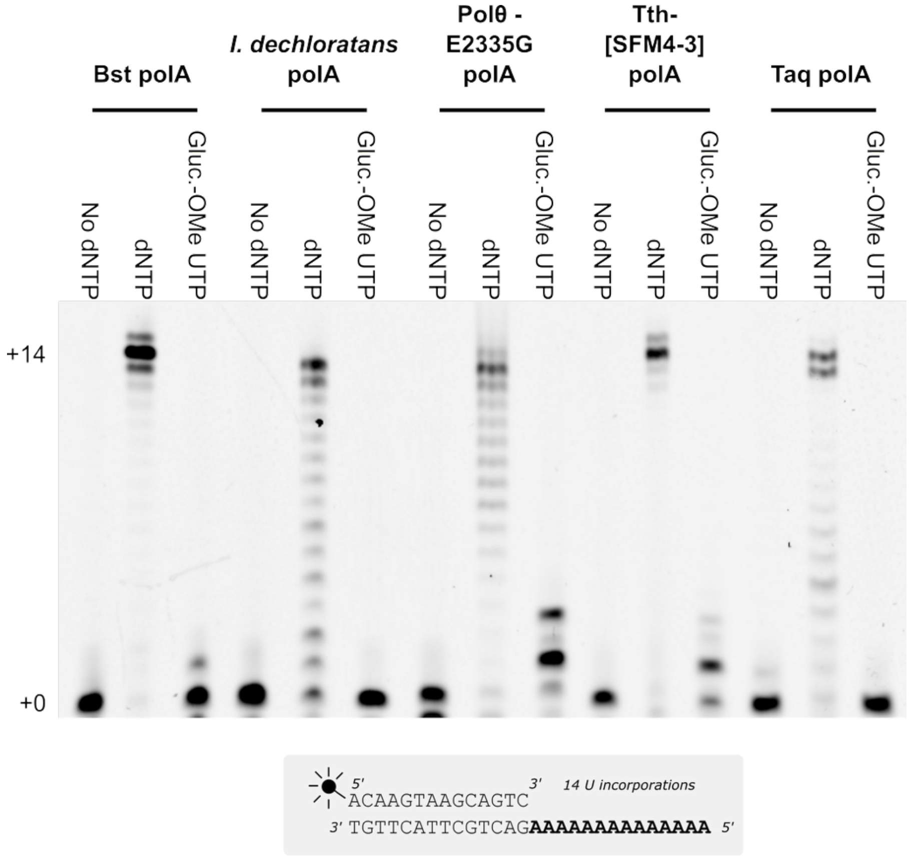


**Supplementary Figure 3 | Basal activity for 5-glucosyl-2'-*O*-methyl-UTP incorporation by various family A DNA polymerases from the literature.**

5-glucosyl-2'-OMe-UTP incorporation was assessed for five representative family A DNA polymerases: Bst polymerase from *Geobacillus stearothermophilus*, Taq polymerase from *Thermus aquaticus*, the polymerase from *Ideonella dechloratans* identified by Czernecki et al.[13], the steric-gate mutant polθ-E2335G described by Randrianjatovo-Gbalou et al.[14], and *Thermus thermophilus* DNA polymerase bearing the mutations reported by Chen et al.[15] for 2'-OMe RNA synthesis.

The mutated polθ is the most efficient enzyme but remains only weakly active, incorporating at most three modified nucleotides, followed by Tth[SFM4-3] and then Bst. All polymerases showed activity in the dNTP control.

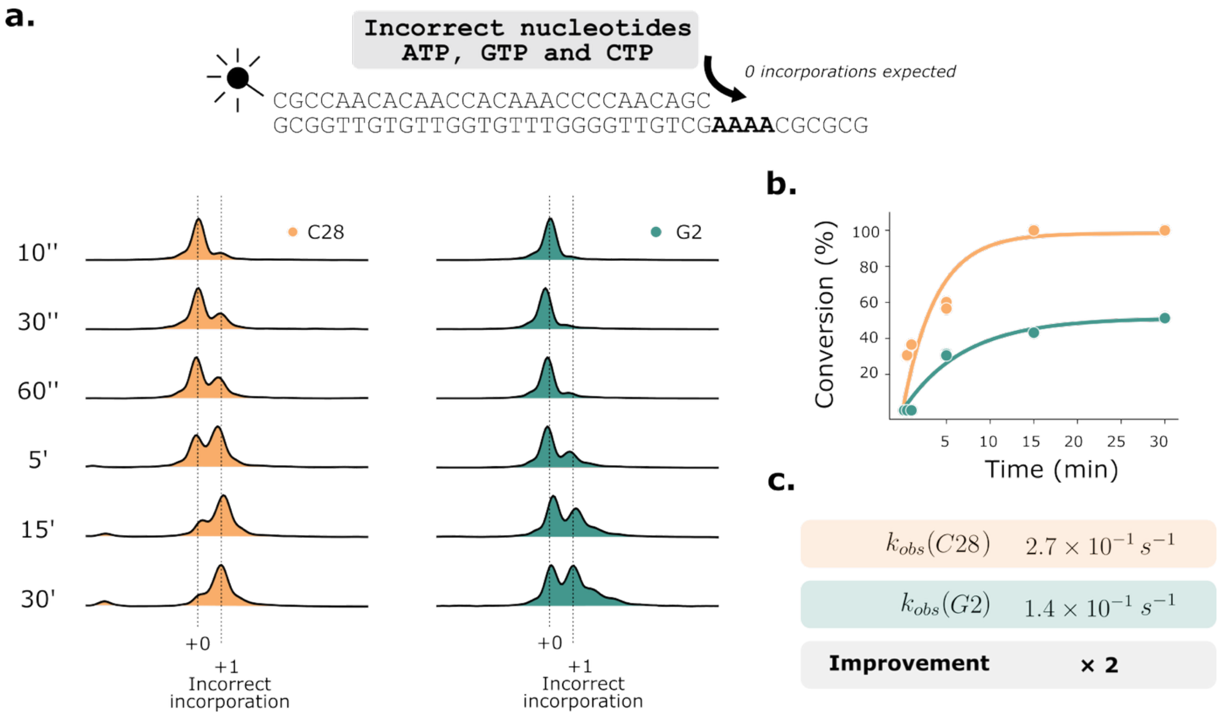


**Supplementary Figure 4 | Fidelity assessment of C28 and G2 polymerases by misincorporation assay.**

A fidelity assay was conducted by capillary electrophoresis to measure the accuracy of C28 and G2 enzymes by testing polymerase activity in the presence of incorrect ribonucleotides (ATP, GTP, and CTP).

**(a)** Time course of incorrect nucleotide incorporation after 10, 30, and 60 seconds, 5, 15, and 30 minutes. After 30 minutes, some overextension products begin to appear for both enzymes.

**(b)** Quantification of >1 extension product over time. Approximately 95% >1 product is observed for C28 after 30 minutes, whereas G2 shows only 52% >1 product at the same time point, indicating reduced misincorporation. Duplicates for each time point.

**(c)** Exponential model fitting to calculate observed rate constants ($k_{obs}$) estimates an approximately two-fold higher fidelity for G2 compared to C28.

### Supplementary Tables

| Round | Library init | Library final | Incubation 55C (h) | Fluorescent Threshold (afu) | Average Event Rate (Hz) | Drops Detected | Hits Detected |
|---|---|---|---|---|---|---|---|
| **1** | IVE001 | IVE002 | 6 | 65 | 693 | 10 934 690 | 126 161 |
| **2** | IVE002 | IVE003 | 6 | 71 | 781 | 8 309 391 | 127 338 |
| **3** | IVE003 | IVE004 | 6 | 73 | 1020 | 4 734 178 | 167 223 |
| **4** | IVE004 | IVE005 | 2 | 60 | 1020 | 10 868 905 | 57 993 |

**Supplementary Table 1 | Throughput of droplet-based directed evolution.**

Four rounds of directed evolution enable screening of millions of droplets, typically sorting more than $10^6$ variants per cycle with a 0.1 droplet occupation.

| Name | Use | Sequence |
|---|---|---|
| **30mer.V2HP.Cy3** | Droplet test | /5Cy3/ACAACCATTATCTAGAGCGATTCGTATAGGTGGTATCCGAAAGGATACCACC |
| **QP08.Iowa** | Droplet test | ATGGTTGT/3IABkFQ/ |
| **iE124** | Primer kinetics | [ATTO488]c*g*c*caacacaaccacaaaccccaac*a*g*c |
| **iE144** | Ladder kinetics +10 | [ATTO488]c*g*c*caacacaaccacaaaccccaac*a*g*cTTTTGCGCGC |
| **iE136** | Template kinetics DNA | GCGCGCAAAAGCTGTTGGGGTTTGTGGTTGTGTTG*G*C*G |
| **iR128** | 38-nt primer | [6FAM]GAGGTCTCGCTCCGACCGCTCCCG |
| **iR131** | 38-nt ladder | [6FAM]GAGGTCTCGCTCCGACCGCTCCCGCATCGTTAGGCACTTGGTTAGGCACTTGGTTAGGCACT |
| **iR135** | 38-nt template | AGTGCCTAACCAAGTGCCTAACCAAGTGCCTAACGATGCGGGAGCGGTCGGAGCGAGACCTC |
| **iR585** | 38-nt complement | GAGGTCTCGCTCCGACCGCTCCCGCATCGTTAGGCACTTGGTTAGGCACTTGGTTAGGCACT |
| **iE140** | Library barcoding | GATCCTCTCATAGTTAATTTCTCCTCTTTAATGAATTCTGBDHVBDHVBDHVBDHVCGACTGATGACGBDHVBDHVBDHVBDHVATCCGCTCACAAAGCAAATAAATTTTTTATG |
| **iE141** | Library barcoding | CAGAATTCATTAAAGAGGAGAAATTAACTATGAGAGGATC |
| **iE142** | Library barcoding | GTCGACCTGCAGCCAAGCBDHVBDHVBDHVBDHVCGACTGATGACGBDHVBDHVBDHVBDHVCCTCAGAACTCCATCTGGATTTGTTCAG |
| **iE143** | Library barcoding | GCTTGGCTGCAGGTCGAC |

**Supplementary Table 2 | Primers and templates for all polymerase studies, mutagenesis and directed evolution.**

| Library | Number of raw reads | Number of clusterized reads (clust. size >5) | Number of clusters (= denoised reads) | Number of unique variants | Number of unique variants with no stop | Share of stop into population (in %) | Mean size of clusters |
|---|---|---|---|---|---|---|---|
| **lVE001** | 8.5 M | 3.41 M | 184 001 | 86 683 | 79 223 | 7.4 | 12 |
| **lVE005** | 6.7 M | 1.49 M | 143 857 | 14 544 | 13 806 | 2.1 | 10 |
| **Total** | **15.2 M** | **4.9 M** | **327 858** | **101 227** | **93 029** | | |

**Supplementary Table 3 | Summary of the high-throughput sequencing (MinION Oxford Nanopore Technology) experiments performed throughout the evolution campaign.**

For each library, the table lists the library name, the number of raw reads before clustering, the number of reads aligned to the reference, the number of clusters (defined as unique barcode pairs supported by ≥5 reads) and their mean cluster size, the number of unique variants recovered, the number of unique variants without premature stop codons and the fraction of reads carrying stop codons in the library.

### Bibliography


1. Parmentier, G., Schmitt, G., Dolle, F. & Luu, B. A convergent synthesis of 2′-o-methyl uridine. *Tetrahedron* **50**, 5361–5368 (1994).

2. Barandun, L. J. *et al.* Replacement of Water Molecules in a Phosphate Binding Site by Furanoside-Appended *lin* -Benzoguanine Ligands of tRNA-Guanine Transglycosylase (TGT). *Chemistry* **21**, 126–135 (2015).

3. Wang, Q., Ma, X., Chen, Y., Jiang, C. & Xu, Y. Electrochemical Synthesis of 5-Selenouracil Derivatives by Selenylation of Uracils. *European J. Org. Chem.* **2020**, 4384–4388 (2020).

4. Cooke, J. W. B., Bright, R., Coleman, M. J. & Jenkins, K. P. Process Research and Development of a Dihydropyrimidine Dehydrogenase Inactivator: Large-Scale Preparation of Eniluracil Using a Sonogashira Coupling. *Org. Process Res. Dev.* **5**, 383–386 (2001).

5. Edgar, R. C. Muscle5: High-accuracy alignment ensembles enable unbiased assessments of sequence homology and phylogeny. *Nat. Commun.* **13**, 6968 (2022).

6. Li, H. Minimap2: pairwise alignment for nucleotide sequences. *Bioinformatics* **34**, 3094–3100 (2018).

7. Li, H. New strategies to improve minimap2 alignment accuracy. *Bioinformatics* **37**, 4572–4574 (2021).

8. Danecek, P. *et al.* Twelve years of SAMtools and BCFtools. *Gigascience* **10**, (2021).

9. Fu, L., Niu, B., Zhu, Z., Wu, S. & Li, W. CD-HIT: accelerated for clustering the next-generation sequencing data. *Bioinformatics* **28**, 3150–3152 (2012).

10. Garrison, E. & Marth, G. Haplotype-based variant detection from short-read sequencing. *ArXiv* http://arxiv.org/abs/1207.3907 (2012).

11. Dalla Pozza, M. *et al.* Enzymatic synthesis of glyco-DNA equipped with oligosaccharides and charged monosaccharides. Preprint at https://doi.org/10.26434/chemrxiv.15001602/v1 (2026).

12. Freund, N. *et al.* A two-residue nascent-strand steric gate controls synthesis of 2′-O-methyl- and 2′-O-(2-methoxyethyl)-RNA. *Nat. Chem.* **15**, 91–100 (2023).

13. Czernecki, D., Nourisson, A., Legrand, P. & Delarue, M. Reclassification of family A DNA polymerases reveals novel functional subfamilies and distinctive structural features. *Nucleic Acids Res.* **51**, 4488–4507 (2023).

14. Randrianjatovo-Gbalou, I. *et al.* Enzymatic synthesis of random sequences of RNA and RNA analogues by DNA polymerase theta mutants for the generation of aptamer libraries. *Nucleic Acids Res.* **46**, 6271–6284 (2018).

15. Chen, T. *et al.* Evolution of thermophilic DNA polymerases for the recognition and amplification of C2′-modified DNA. *Nat. Chem.* **8**, 556–562 (2016).
